# A chromosome-scale strawberry genome assembly of a Japanese variety, Reikou

**DOI:** 10.1101/2021.04.23.441065

**Authors:** Kenta Shirasawa, Hideki Hirakawa, Shinobu Nakayama, Shigemi Sasamoto, Hisano Tsuruoka, Chiharu Minami, Akiko Watanabe, Yoshie Kishida, Mitsuyo Kohara, Manabu Yamada, Tsunakazu Fujishiro, Akiko Komaki, Sachiko N Isobe

## Abstract

Cultivated strawberry (*Fragaria* × *ananassa*) is an octoploid species (2n = 8x= 56) that is widely consumed around the world as both fresh and processed fruit. In this study, we report a chromosome-scale strawberry genome assembly of a Japanese variety, Reikou. The Illumina short reads derived from paired-end, mate-pair, and 10X Genomics libraries were assembled using Denovo MAGIC 3.0. The generated phased scaffolds consisted of 32,715 sequences with a total length of 1.4 Gb and an N50 length of 3.9 Mb. A total of 63 pseudomolecules including chr0 were created by aligning the scaffolds onto the Reikou S1 linkage maps with the IStraw90 Axiom SNP array and ddRAD-Seq. Meanwhile, genomes of diploid *Fragaria* species were resequenced and compared with the most similar chromosome-scale scaffolds to investigate the possible progenitor of each subgenome. Clustering analysis suggested that the most likely progenitors were *F. vesca* and *F. iinumae*. The phased pseudomolecules were assigned the scaffolds names with Av, Bi, and X, representing sequence similarity with *F. vesca* (Av), *F. iinumae* (Bi), and others (X), respectively. The result of a comparison with the Camerosa genome suggested the possibility of subgenome structure differences between the two varieties.

## Introduction

Polyploidization is often observed in plants and has contributed to evolution and domestication by increasing plant size and stress tolerance. It has also played important roles in human life and has been used for crops such as wheat, cotton, banana, potato, sweetpotato, and strawberry. Polyploidy is defined as “the heritable condition of processing more than two complete sets of chromosomes”[1]. Because of their complex structures, polyploid species have been neglected in genetic and genomic studies. However, recent advances in next-generation sequencing (NGS) and bioinformatics approaches provide new insights into polyploid plant genomes. When Kyriakidou et al. (2018)[2] reviewed progress in polyploid plant genome sequence assembly, a total of 47 assembled sequences were available as of May 2018. One of the breakthroughs in polyploid genome assembly technology is the development of bioinformatic analysis for the phasing of heterozygous genome sequences, such as Falcon phase[3], Trio binning[4], and Denovo MAGIC™ software (https://www.nrgene.com/solutions/denovomagic/).

Cultivated strawberry (*Fragaria* × *ananassa*) is an octoploid species (2n = 8x= 56) that is widely consumed around the world as fresh or processed fruit. The genome size of strawberry is estimated as 1C = 708–720 Mb[5, 6]. Strawberry was generated by artificial interspecific crosses between *F. chiloensis* and *F. virginiana* in 18th century Europe[7]. It was first considered an allo-autopolyploidy species (AABBBBCC[8], AAA’A’BBBB[9], then the allopolyploidy structure (AAA’A’BBB’B’) was suggested based on isozyme segregation[10]. Later, the genome structure of AvAvBiBiB1B1B2B2 was proposed by Tenessen et al. (2014)[11], reflecting the possible progenitor genomes of *F. vesca*.

The genome of cultivated strawberry was first sequenced at the beginning of the next-generation sequencing (NGS) era by using Roche 454, Illumina Solexa, and Life Technologies[12]. The sequenced material was Reikou, a Japanese variety, that was bred at the Chiba Prefectural Agriculture and Forestry Research Center in 1979. Reikou was chosen because it was a founder of many important Japanese varieties, such as Nyoho, Tochiotome, Benihoppe, and Fukuoka S6 (http://www.naro.affrc.go.jp/org/narc/evotree/tree_view/strawberry/index.html). It was one of the first trial of *de novo* genome assembly in polyploidy species; however, the assembled sequences were fragmented because of immature technologies in sequencing and assembly tools.

Recently, a chromosome-scale genome assembly in cultivated strawberry was reported by Edger et al. (2019)[13]. The Illumina paired-end (PE), mate-pair (MP), and 10X Genomics libraries were sequenced for a strawberry variety, Camarosa, which has been widely cultivated around the world. The Illumina reads were assembled by Denovo MAGIC 3 (NRGene), and the chromosome-scale scaffolds were created with the Hi-C reads. PacBio reads were also used to fill gaps. Edger et al. (2019) compared the assembled genome and transcript sequences of diploid *Fragaria* spp sequences on the basis of the assembled sequences and hypothesized that two octoploid progenitors of cultivated strawberry, *F. chiloensis* and *F. virginiana*, were derived from four diploid species, *F. iinumae, F. nipponica, F. viridis*, and *F. vesca*.

The genome sequences assembled by Edger et al. (2019) have enhanced the genetic and genomic study of cultivated strawberry. Meanwhile, homoeologous recombination was reported in several polyploid species, such as Brassica *napus*[14], cotton[15], and coffee[16]. Therefore, there is a possibility that homoelologous recombination has occurred in strawberry too and that the subgenome structure varies among the accessions. Although Edger et al. (2019) suggested the origin of strawberry based on their assembled genome, Liston et al. (2020)[17] immediately argued against that hypothesis on the basis of the results of a chromosome-scale phylogenomic and phylogenetic analysis based on an *F. moschata* linkage map. This debate suggests that a single reference genome might not be enough to draw conclusions about the features of the strawberry genome or its evolutional history. Hence, we report another chromosome-scale strawberry genome assembly of a Japanese variety, Reikou. Camarosa was a variety bred at the University of California, USA, in 1992, with a different breeding history than Reikou. Thus, the Reikou genome is expected to suggest another aspect of the strawberry genome structure by comparison with the Camarosa genome and to advance the discussion of strawberry genetics and genomics.

## Materials and methods

### Sampling strategy

Japanese strawberry variety Reikou, a founder of elite cultivars in Japan, was used for genome sequencing analysis (Supplementary Table S1). A self-pollinated S1 population of Reikou, consisting of 157 progenies, was used for ddRAD-Seq analysis and linkage map construction. The materials were grown at the Chiba Prefectural Agriculture Research Center. A total of 14 wild diploid species, *F. bucharica, F. chinensis, F. daltoniana, F. glacilis, F. hayatae, F. iinumae, F. mandshurica, F. nilgirrensis, F. nipponica, F. nipponica* (accession Yakushima), *F. nubicola, F. pentaphylla, F. vesca* (accession Hawaii 4), and *F. viridis* (Supplementary Table S2), were used for whole-genome resequencing analysis. Leaves of these 14 species were provided by Prof. Tomohiro Yanagi of Kagawa University. DNA was isolated from young leaves with the DNeasy Plant Mini Kit (Qiagen, Hilden, Germany).

### Whole-genome shotgun and linked sequencing analysis for genome assembly

Whole-genome shotgun paired-end (PE) libraries (insert sizes of 470 bp and 800 bp) and mate-pair (MP) libraries (3, 6, and 9 kb) were constructed from the Reikou genome DNA. Nucleotide sequences of the libraries were determined with theHiSeq 2500 DNA sequencing system (Illumina, San Diego, CA) in paired-end, 260-bp mode for the 470-bp paired-end library and 160-bp mode for the other four libraries. A linked sequencing library based on 10X Genomics Chromium technology (10x Genomics, Pleasanton, CA) was constructed and sequenced on Illumina HiSeqX in paired-end, 260-bp mode. The sequences were assembled with DenovoMAGIC 3.0 software, which was a De Bruijn-graph-based assembler, by the NRGene company (Ness Ziona, Israel), and phased scaffold sequences were generated.

### Construction of chromosome-scale pseudomolecule sequences

Since simplex SNPs were expected to be mapped on the genetic map based on the Reikou S1 mapping population constructed in our previous study[18], the alleles of the simplex SNPs could be identified by a similarity search of the flanking sequences of SNPs in the IStraw90 Axiom SNP array[19] against the assembled sequences. Theoretically, one franking sequence should hit eight homoeologous genome sequences, one of which would have the same bases located on the map as the simplex SNPs. Because the simplex SNPs were in coupling phase between scaffold sequences on a single haplotype, the sequences were connected with 10,000 Ns to obtain two phased pseudomolecule sequences with suffixes of a and b.

The qualities of the two-phased pseudomolecule sequences were furthermore investigated by aligned onto a ddRAD-Seq linkage map generated from the same Reiko S1 mapping population. The total genomic DNA of the Reikou S1 progeny used in our previous study[18] was digested with two restriction enzymes, *PstI* and *MspI*. The ddRAD-Seq libraries were constructed and sequenced on a HiSeq 2000 platform (Illumina) in paired-end 93 bp mode. The reads obtained from the ddRAD-Seq were processed and mapped by Bowtie 2[20] on the two-phased pseudomolecule sequences to call SNP candidates as described in Shirasawa et al. (2016)[21]. From the ddRAD-Seq data, high-quality biallelic SNPs were selected using the following criteria: include only genotypes supported by reads of ≥ 5 (--minDP 5); include only sites with quality values ≥ 999 (--minQ 999); exclude sites with ≥ 75% missing data (--max-missing 0.75); fit the expected ratio of 1:2:1 via Chi-square tests (PD≥D0.01); and the presence of a heterozygote in the parental line (Reikou itself). Linkage analysis of the SNPs was performed with Lep-Map3[22] for the construction of phased linkage maps. The pseudomolecule sequences constructed on the IStraw90 Axiom SNP array-based map were re-aligned on the ddRAD-Seq maps to replace the misplaced scaffolds. The revised pseudomolecule sequence was designated as the FAN_r2.3.pseudomolecule.

### Gene prediction

Putative protein-encoding genes were predicted with ab-initio-, evidence-, and homology-based gene prediction methods in a MAKER pipeline[23] Publicly available RNA-Seq reads of *Fragaria* × *ananassa* (SRX1895539, SRX1895540, and ERX935139 to ERX935144) and *F. vesca* (SRX225676) were mapped on the FAN_2.3 sequences with Tophat2[24] to presume gene positions on the genome sequences. The gene position information was used in AUGUSTUS[25] and GeneMark[26], both of BREAKER1[27] to generate training datasets. On the basis of another training preset of SNAP[28] for Arabidopsis as well as the two sets, genes were predicted from the FAN_2.3 sequences with the MAKER pipeline, for which unigene sets of the RNA-Seq and peptide sequences of *F. vesca* (v1.1.a2), *Prybys persica* (v2.0.a1), and *Malus*□×□*domestica* (v1.0p), all of which are registered in the Genome Database for Rosaceae[29], were employed. Finally, genes with an annotation edit distance[30] score of ≤0.5 and a length of >100 amino acids were selected as a final dataset consisting of 167,721 gene models. A sequence similarity search of genes was performed with BLASTN[31] with an E-value cutoff of 1E-50.

### Comparison with the diploid Fragaria sequences and the assembled genomes

Whole-genome shotgun resequencing analysis was performed across the 14 diploid *Fragaria* species. PE libraries with an insert size of 500 bp were constructed with the TruSeq Nano DNA Prep Kit (Illumina) and sequenced on NextSeq 500 (Illumina) in paired-end, 151-bp mode. The obtained sequences were mapped onto the pseudomolecule sequences (FAN_r2.3.pseudomolecule) by Bowtie 2[20], and SNPs were called the same as in the above ddRAD-Seq analysis. The called candidate SNPs were filtered out according to the following criteria: include only genotypes supported by reads ≥ 10 (--minDP 10); include only sites with quality values ≥ 999 (--minQ 999); exclude sites with ≥ 50% missing data (--max-missing 0.5); include only sites with minor allele frequencies ≥ 0.2 (--maf 0.2). Dendrograms of the 14 diploid species and *F*. × *ananassa* were drawn with TASSEL 5[18, 32].

Comparative analyses of the genome sequences between the two phased sequences of the FAN_r2.3.pseudomolecule and between the two species, *F. × ananassa* (FAN_r2.3.pseudomolecule) and *F. vesca* (v1.1), were performed with NUCmer of the MUMmer package [23]. Comparisons between the FAN_r2.3.pseudomolecule and the Camerosa genome[13] were performed by using D-Genesis[33].

## Results and Discussion

### Phased genome assembly with Illumina and 10X Genomics technologies

Whole-genome shotgun sequencing libraries for the Illumina and 10X Genomics platforms were constructed from genome DNA of a Japanese variety, Reikou. For Illumina sequencing, five libraries with insert sizes of 470 bp, 800 bp, 3 kb, 6 kb, and 9 kb were constructed and sequenced. After trimming the adapter sequences and low-quality bases, a total of 330.4 Gb of high-quality paired-end sequence reads was obtained (Supplementary Table S1). The obtained data size corresponded to 466x (4n) and 233x (8n) genome coverage when the estimated genome size is 708 Mb[5]. The paired-end reads were assembled with DenovoMAGIC 3.0 (NRGene) to generate an assembly of the Reikou genome (Table 1). To extend the sequences, another sequencing library for 10X Genomics was constructed and sequenced to obtain a total of 122.7 Gb, 86.7x genome (8n) coverage (Supplementary Table S1). The reads were integrated with the Illumina assembly to extend the sequence contiguity. The resultant genome assembly, consisting of 32,715 sequences, had a total length of 1.4 Gb with an N50 length of 3.9 Mb (Table 1). The total length of 1.406 was almost the same as the estimated genome size of 2C (1.416 Gb) [5], suggesting that the assembly represented phased genome sequences corresponding to 8n.

**Table 1.** Assembly statistics

|  | Denovo MAGIC<br>assembly with PE<br>reads | Denovo MAGIC<br>assembly with PE+<br>10X Genomics<br>reads | FAN_2.3<br>scaffold<br>sequences | FAN_2.3.<br>pseudomolecule<br>sequences |
| --- | --- | --- | --- | --- |
| Number of<br>scaffold<br>sequences | 7,563 | 32,715 | 32,771 | 63 |
| Total size of<br>scaffold<br>sequences (bp) | 1,108,561,665 | 1,406,451,310 | 1,406,430,992 | 1,733,510,992 |
| Scaffold N50 (bp) | 3,886,453 | 3,887,271 | 3,370,964 | 28,854,432 |
| Longest scaffold<br>(bp) | 14,978,214 | 17,025,478 | 10,978,170 | 607,790,169 |
| GC (%) | 39.2 | 39.3 | 39.3 | 39.3 |
| Gap represented<br>by N (%) | 3 | 2.2 | 2.2 | 20.7 |

### Construction of chromosome-scale pseudomolecule sequences based on genetic maps

Haplotype-based genetic maps, on which only coupling-phase alleles are located, would be useful for assigning phased genome sequences to chromosomes for the construction of chromosome-scale pseudomolecule sequences. In our previous study[18], a genetic map consisting of 31 linkage groups for Reikou itself has been established with 11,002 simplex SNPs from the IStraw90 Axiom SNP array[19], segregating the S1 mapping population of Reikou; however, the map was not phased, meaning that coupling-phase alleles and repulsion-phase alleles were simultaneously mapped on a single linkage group. To generate haplotype-based genetic maps, i.e., phased maps, from the unphased genetic map, it is necessary to identify coupling-phase alleles. Therefore, probe sequences of the SNP array were searched against the phased assembly sequences to find eight theoretical homoeologous sequences, only one of which should be identical to those on the map. As expected, in 7,232 SNPs, one probe sequence hit 6.1 genome sequences on average, one of which had the same base as the mapped SNP alleles. As a result, the genetic map was split into 31 pairs of 62 haplotype-based linkage groups.

The phased genome assembly was assigned on the haplotype-based genetic map. During the assignment process, 40 misassembly points were broken to reduce misassembly errors. In addition, 80 sequences (20,318 bp) were removed due to the possibility of contamination of sequences from organelles and bacteria. The resultant set of sequences, consisting of 32,771 sequences covering 1,406,430,992 bases, was designated as FAN_r2.3 (Table 1). Out of the 32,771 sequences, 464 spanning 1,121,700,823 bases (79.8%) were assigned to the 62 linkage groups as pseudomolecule sequences. The remaining 32,307 scaffold sequences (284,730,169 bp) were grouped as chromosome 0 (ch0).

These pseudomolecule sequences were further validated with another genetic map based on ddRAD-Seq technology. From the same S1 population as above, 1.8 million high-quality sequence reads on average per individual were obtained (Supplementary Table S1). The reads were aligned on the two haplotype sequences with 87.9% and 87.8% alignment rates. Totals of 8097 and 8189 selected high-confidence SNPs were mapped together on 62 linkage groups. In this analysis, five misoriented scaffold sequences were found and inverted to the correct direction. The resultant sequence dataset was designated as the FAN_2.3.pseudomolecule (Table 1). Benchmarking Universal Single-Copy Orthologues (BUSCO) analysis revealed that the FAN_2.3.pseudomolecule had 1,346 (93.5%) complete orthologues [181 (12.6%) single-copy and 1,165 (80.9%) duplicated] and 14 (1.0%) fragmented orthologues, indicating that the sequences had good coverage of the gene space of the strawberry genome.

### Assignment of chromosome numbers and subgenome names to the pseudomolecule sequences

Whole-genome shotgun sequence data were obtained from 14 wild *Fragaria* species (Supplementary Table S2) as well as from *F*. × *ananassa* (Reikou, DRA accession number DRR013874). High-quality reads were mapped on the constructed pseudomolecules (FAN_2.3.pseudomolecule). The average map rate of the reads on the entire pseudomolecule sequences was 81.4%. In each sequence, on the other hand, the map rates of *F. vesca*, a probable A-genome ancestor, and 12 other species (the exception being *F. iinumae*) were prominent in a couple of sequences (but 4 in ch6), while those of *F. iinumae* (B genome) were predominant in another couple of sequences (Supplementary Table S2).

From the mapping alignments, a total of 864,147 SNPs were detected in the 15 species with respect to the FAN_2.3.pseudomolecule (Supplementary Table S3). Based on the SNPs, a dendrogram was constructed for each pseudomolecule sequence. The dendrograms were classified into three types, depending on the shapes of the branches: 1) *F. × ananassa* and *F. vesca* in the same branches, 2) *F. × ananassa* and *F. iinumae* in the same branches, and 3) *F. × ananassa, F. vesca*, and *F. iinumae* in the same branches (Supplementary Figure S1).

The results of the mapping and SNP analyses suggested that each pseudomolecule sequence would be related to either genome of the wild species. Therefore, it was possible to assign the chromosome numbers and subgenome names to the pseudomolecule sequences. For four pairs of haplotype sequences of seven chromosomes, homologous pairs of Av (A genome from *F. vesca*), Bi (B genome from *F. iinumae*), and X1 and X2 (unknown genomes from anonymous species) were assigned. In addition, ch1X3 was an extra pair of sequences in ch1, and ch6Av2 and ch6×3 were extra pairs of sequences in ch6. Each haplotype was distinguished with suffixes a and b.

Edger et al. (2019)[13] suggested that the possible progenitors of strawberry were *F. vesca, F. iinumae, F. nipponica*, and *F virdis* based on sequence similarity with the wild *Fragaria* species. However, later reports suggested that *F. vesca* and *F. iinumae* were possible progenitors, whereas *F. nipponica* and *F. virdis* were not[17, 34, 35]. The result of our clustering analysis also denied the possibility of *F. nipponica* and *F. virdis* as progenitors of the strawberry genome.

The structure of the strawberry genome based on the FAN_r2.3.pseudomolecule basically matched that of *F. vesca* (v1.1) but included possible chromosome rearrangements, e.g., inversions and deletions over the genomes (Figure 1A, Supplementary Figure S2). Comparative analysis of genome structures between the two haplotypes indicated that most parts were well conserved (Figure 1B, Supplementary Figure S3) but had 2,774,528 SNPs and 607,776 indels (sizes of 1 bp to 86 bp).

**Figure 1.**
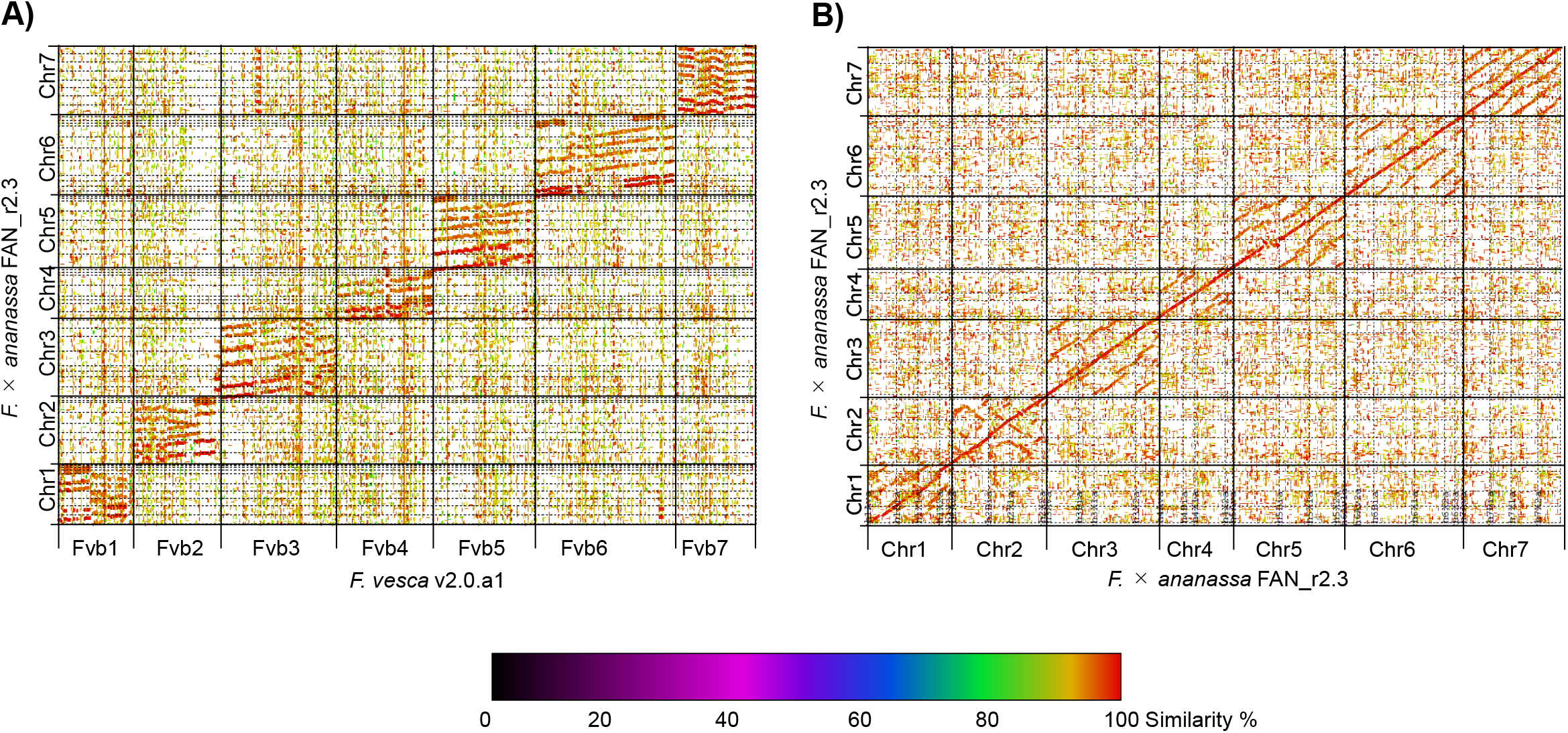
Genome-wide synteny relationships in strawberry genomes. A) Synteny in the genomes of *F*. × *ananassa* (FAN_r2.3.pseudomolecule) and *F. vesca* (F. vesca v1.1) B) Synteny between two phased sequences of the *F*. ×*ananassa* genome (FAN_r2.3.pseudomolecule)

### Gene prediction and copy number variations in the octoploid genome

A total of 167,721 protein-coding genes, whose mean length was 1,272 bases, were predicted from the FAN_2.3.pseudomolecule sequences (Table 2). The predicted genes included 1,407 (97.7%) complete [47 (3.3%) single-copy and 1,360 (94.4%) duplicated] and 14 (1.0%) fragmented BUSCO. Of those, 43,931 (26.2%), 29,389 (17.5%), and 59,922 (35.7%) genes were on Av, Bi, and X genomes, respectively, while the remaining 34,479 (20.6%) genes were on ch0, a group of scaffold sequences unassigned to any chromosomes.

**Table 2.**
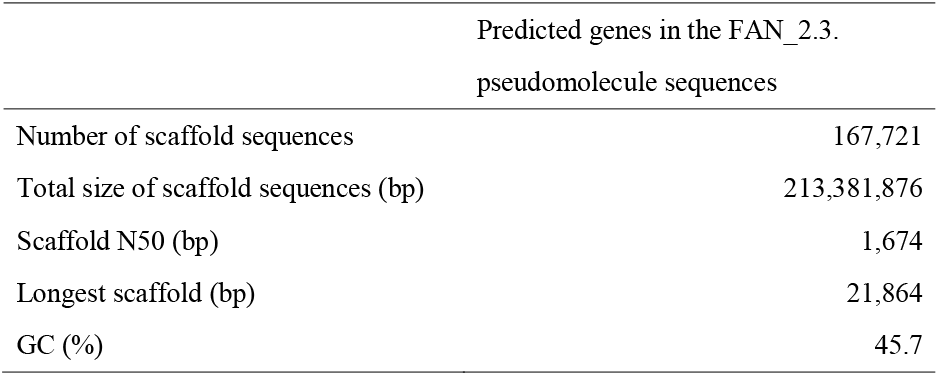
Assembly statistics of the predicted genes

The copy numbers of the genes were investigated. The sequence similarities of the 167,721 genes in the FAN_2.3.pseudomolecule were searched with genes predicted in the *F. vesca* genome (v1.1.a2). As expected, one *F. vesca* gene hit an average of 8.4 genes in the FAN_2.3.pseudomolecule (Figure 2A), since each pseudomolecule sequence represented eight genomes of the octoploid. Among 27,985 *F. vesca* genes hit to either Av, Bi, or X sequences, 23,669 (84.6%), 16,749 (59.8%), and 22,635 (80.9%) were found in Av, Bi, and X genomes, respectively (Figure 2B). While 12,046 (43.0%) genes were shared by all of the Av, Bi, and X sequences, 2,743 (9.8%), 759 (2.7%), and 1,461 (5.2%) genes were proper to Av, Bi, and X sequences, respectively.

**Figure 2.**
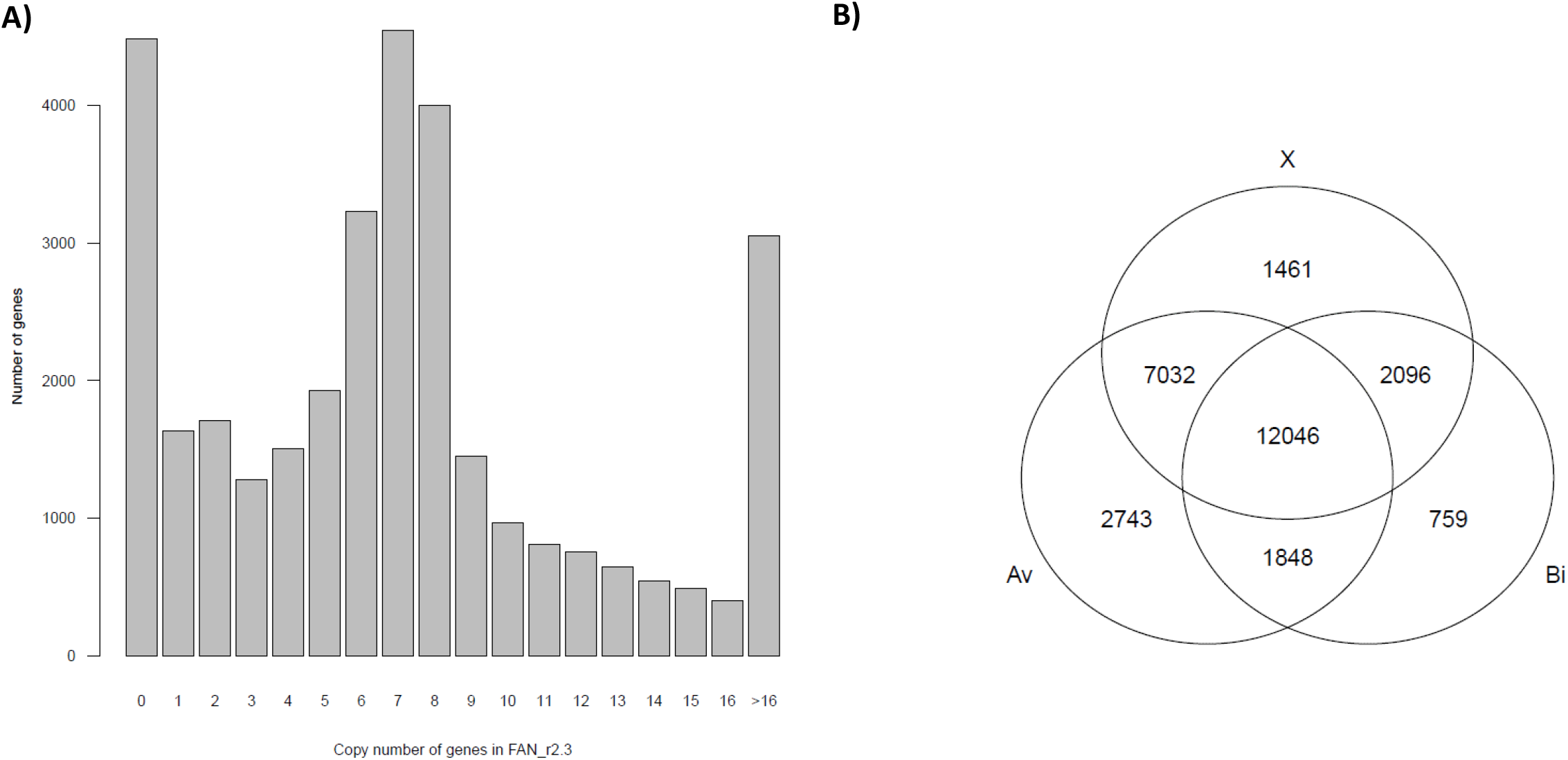
Gene duplication in the *F*. ×*ananassa* genome. A) Copy number of genes in the *F*. ×*ananassa* genome (FAN_r2.3) estimated with the sequence similarity analysis of genes of F. vesca (v1.1.a2) B) Number of genes shared by three subgenomes, Av, Bi, and X

### Comparison with the Camerosa genome

The structures of the FAN_r2.3 pseudomolecule sequences were compared with those of another *F*. × *ananassa* assembled genome, Camerosa v1.0[13] (Figure 3). The camerosa genome consisted of unphased sequences, whereas the Reikou genome (FAN_r2.3) consisted of phased sequences. Each pair of Reikou sequences showed the highest similarity (dark green dots) to the one Camerosa sequence in most regions, suggesting the correspondence of subgenomes between the two assembled genomes. The subgenome sequences between the two genomes often showed nonstraight linear correlations, representing different structure between the two genomes. The results suggested two possibilities: 1) the subgenome structure was different between the two varieties, 2) either genome was misassembled.

**Figure 3.**
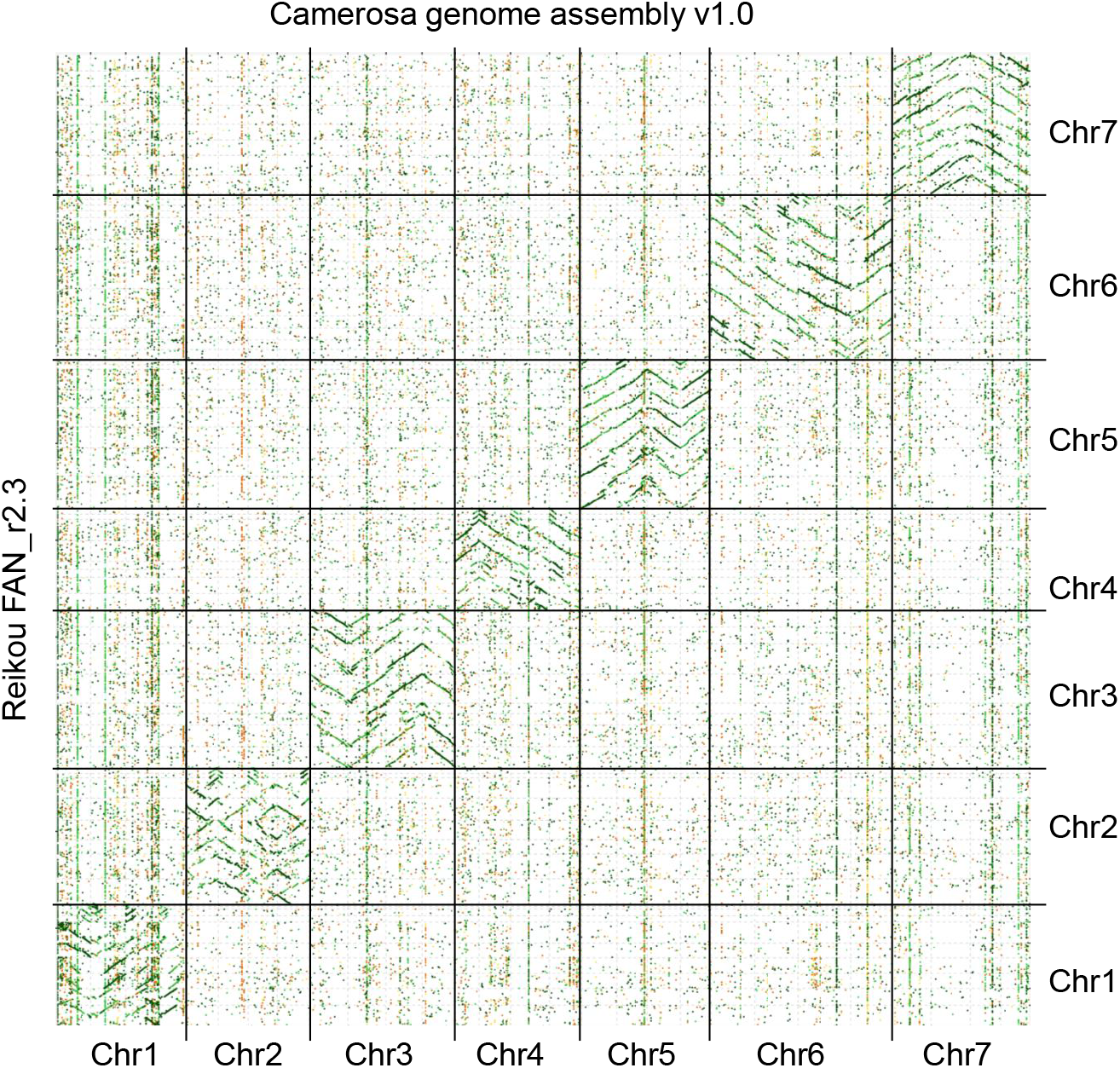
Comparison of the genome structure between Reikou (FAN_r2.3) and Camerosa (v1.0)

### Conclusion

In this study, we reported *de novo* whole genome sequence assembly in a Japanese strawberry variety, Reikou. The phased pseudomolecules were assigned the scaffolds names with Av, Bi, and X, representing sequence similarities with the wild *Fragaria* species, *F. vesca* and *F. iinumae* and others. The distinction between subgenome sequences with ancestorial similarities agreed with the previous genetic and cytological studies, suggesting that *F. vesca* and *F. iinumae* were the most likely progenitors. The result of the comparison with the Camerosa genome suggested the possibility of subgenome structure differences between the two varieties.

## Data availability

The sequence reads are available from the DDBJ Sequence Read Archive (DRA) under the accession numbers DRA007606-DRA007609 and the BioProject ID PRJDB5536. The genome assembly data, including scaffold and pseudomolecule sequences and gene models are available at Strawberry GARDEN (http://strawberry-garden.kazusa.or.jp/).

## Acknowledgments

We thank Prof. Tomohiro Yanagi for providing the plant materials of diploid Fragaria species. This work is partially supported by the Research Program on Development of Innovative Technology, National Agricultural Research Organization, BRAIN (30020B) and by SIP Technologies for Smart Bio-Industry and Agriculture (DDB2006).

## Notes

http://strawberry-garden.kazusa.or.jp/

### Competing Interest Statement

The authors have declared no competing interest.

