## Supplementary Figure S1 for "A chromosome-scale strawberry genome assembly of a Japanese variety, Reikou"

Dendrograms based on SNPs detected in each haplotype across the 14 strawberry line

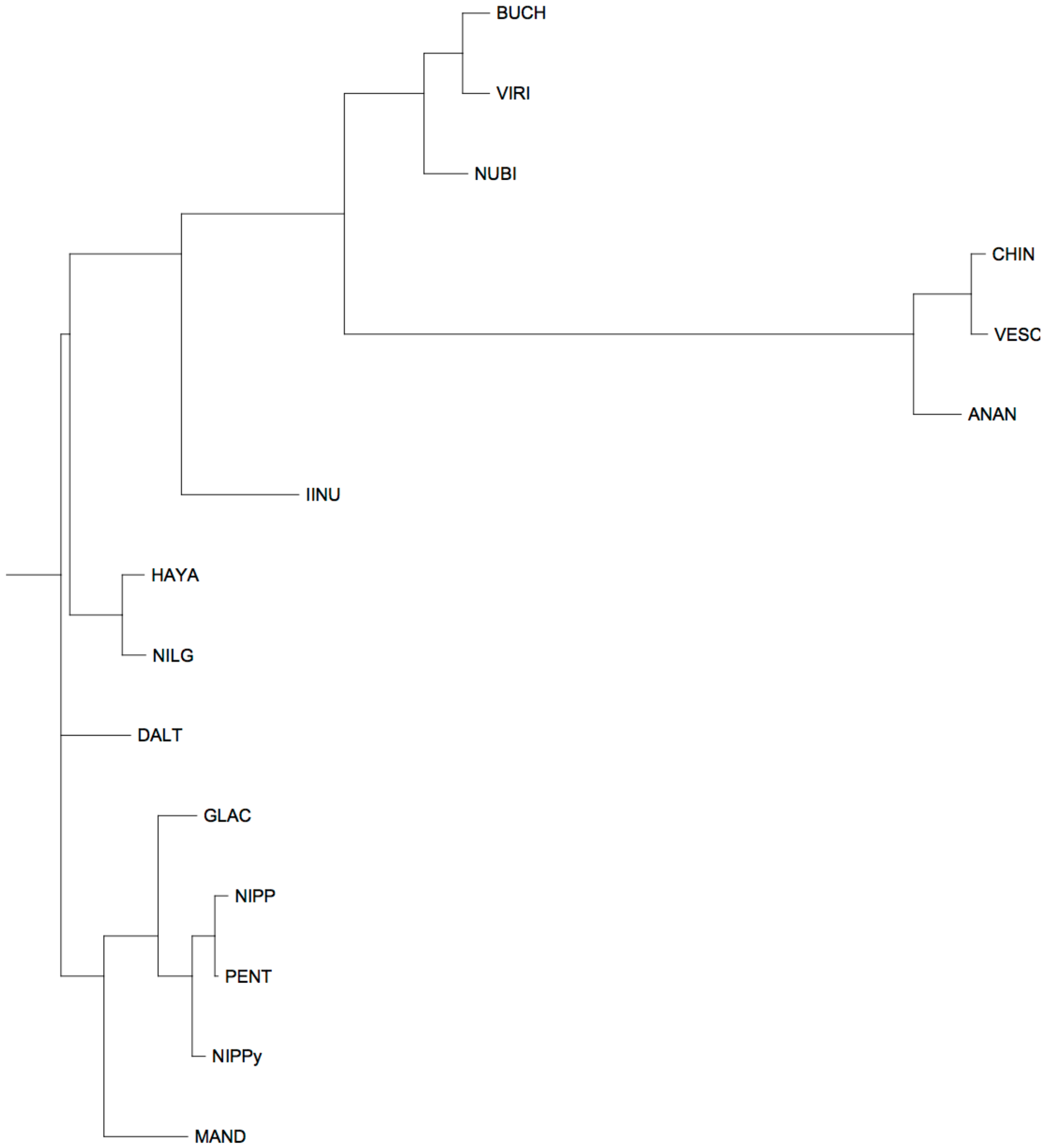


FAN_r2.3ch1Ava

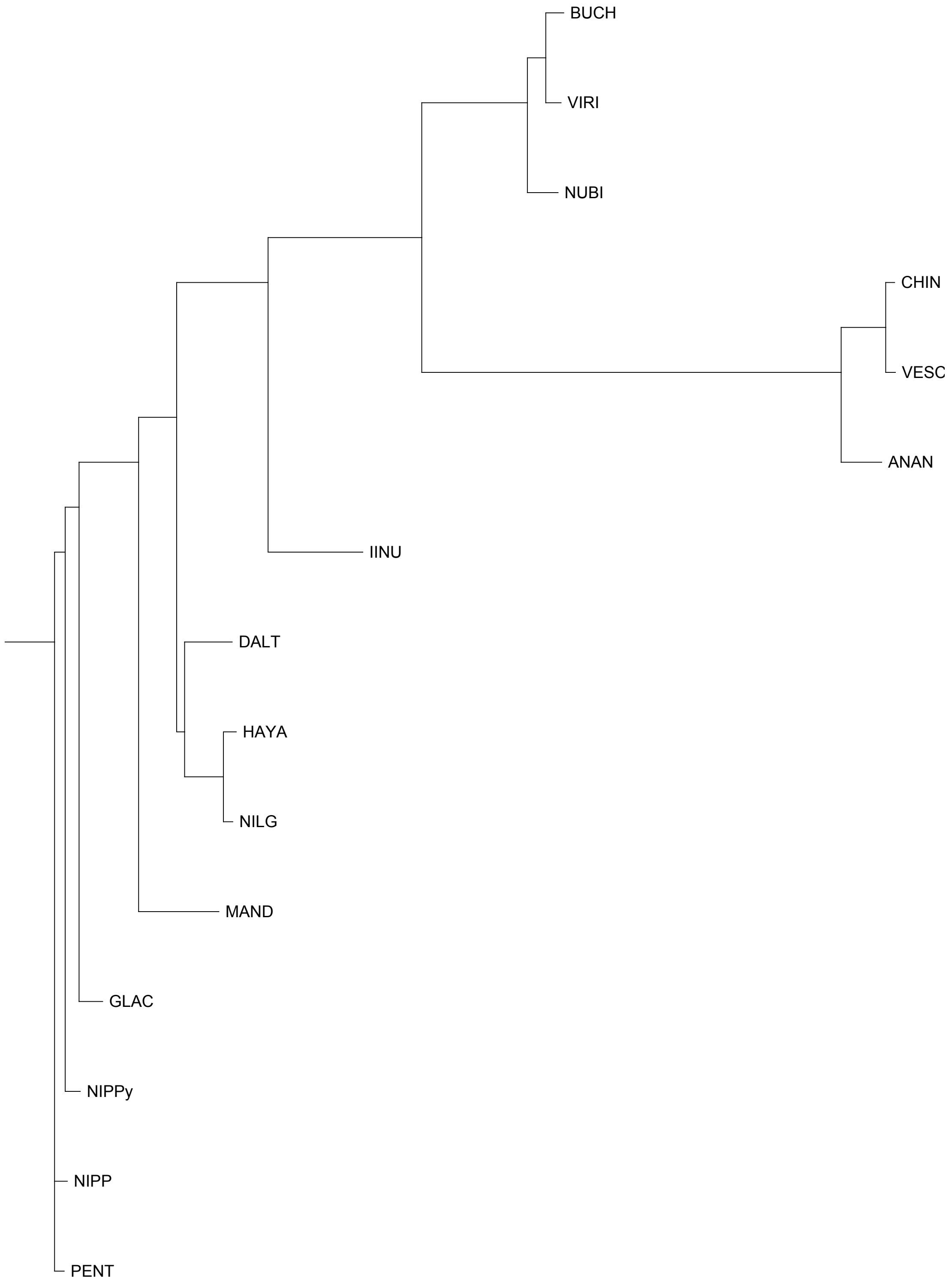


FAN_r2.3ch1Avb

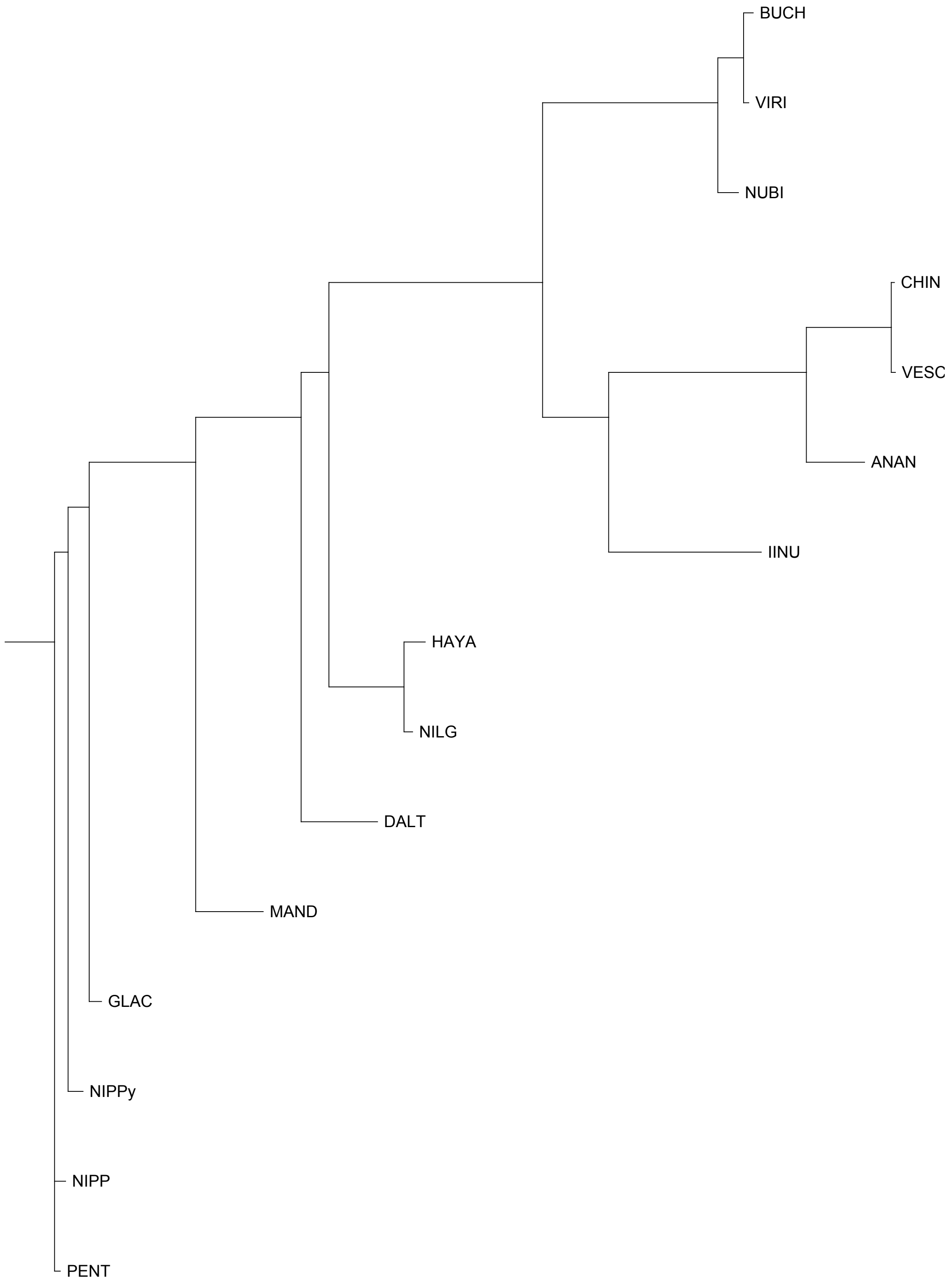


FAN_r2.3ch1Bia

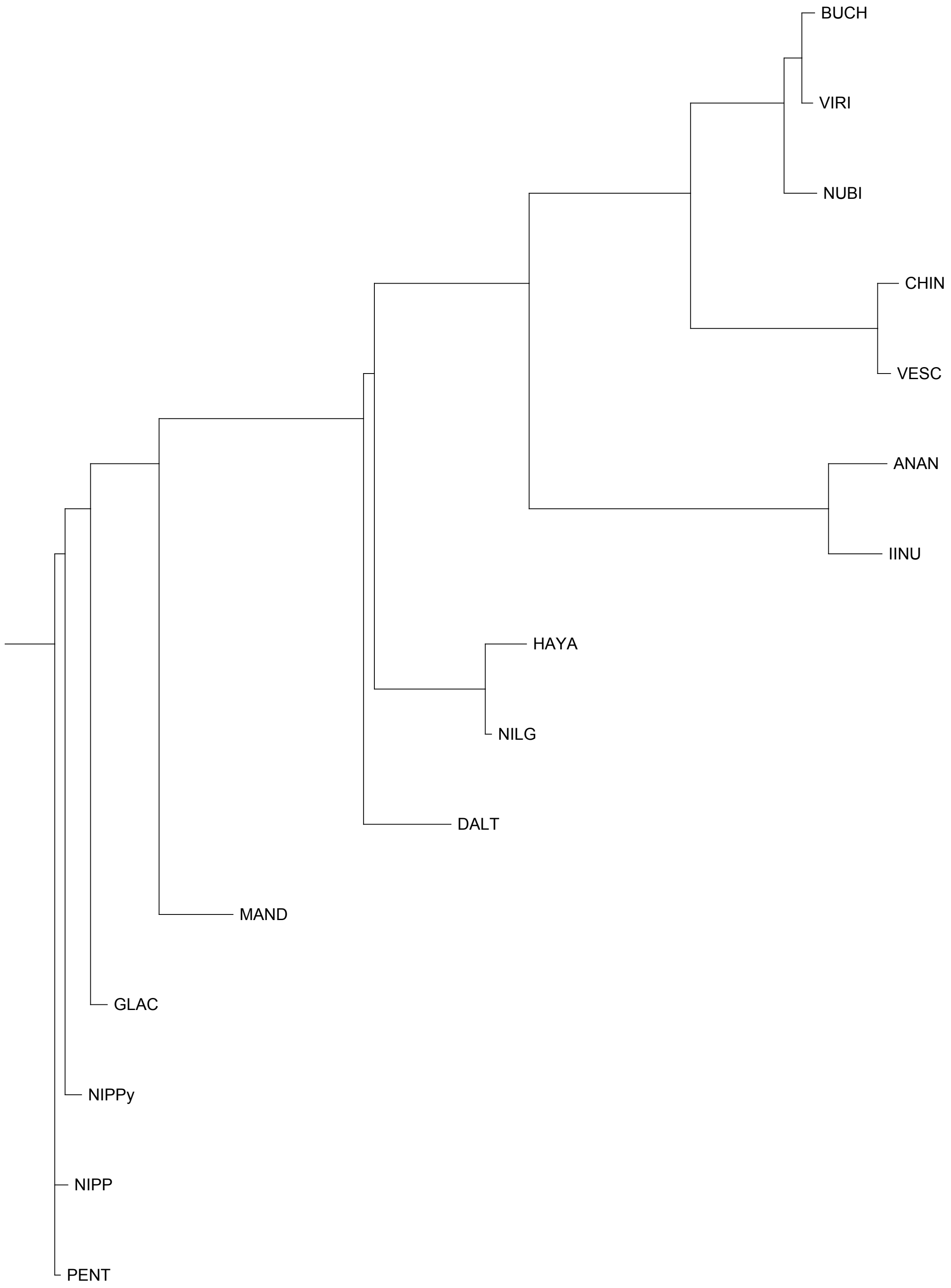


FAN_r2.3ch1Bib

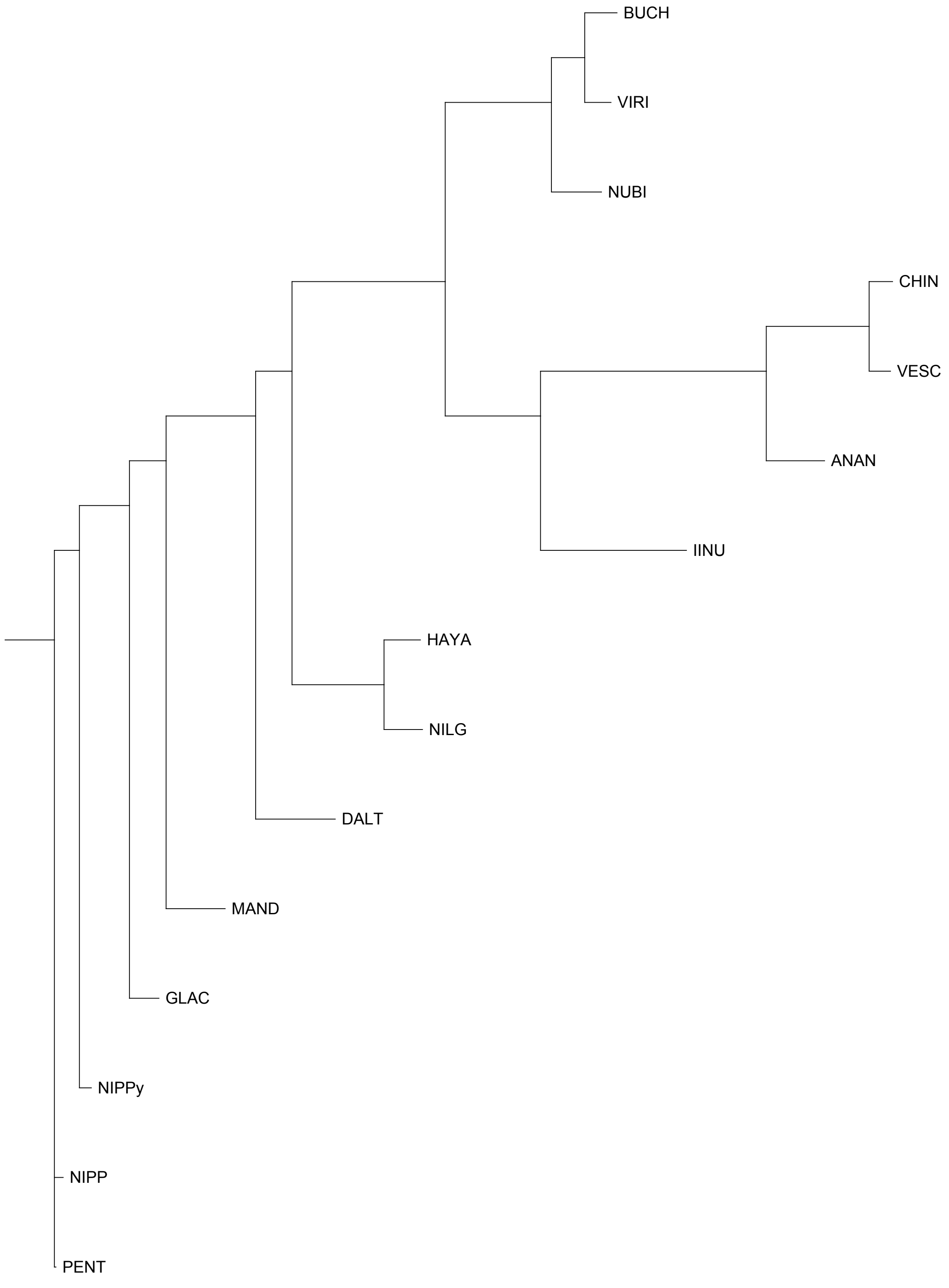


FAN_r2.3ch1X1a

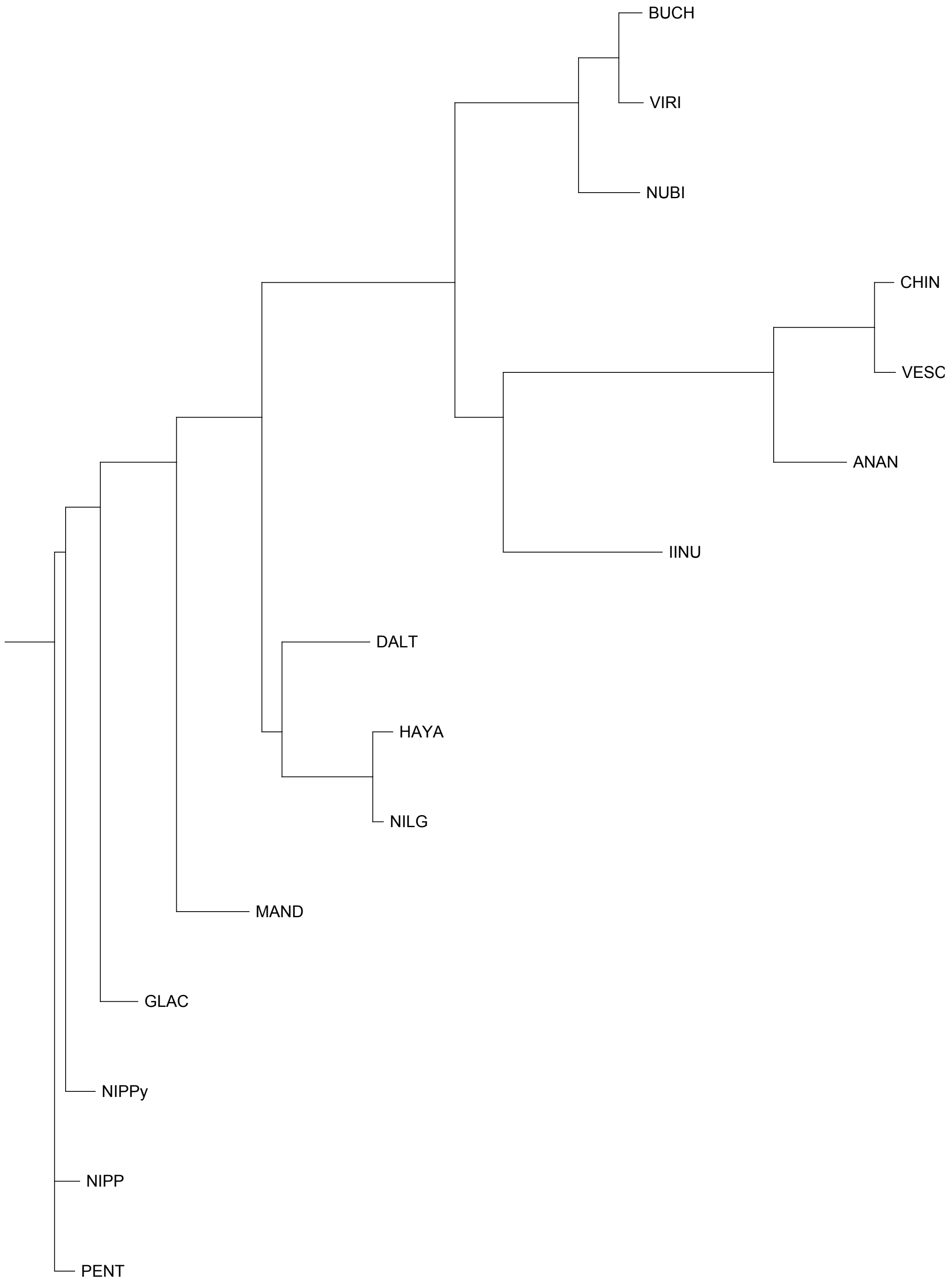


FAN_r2.3ch1X1b

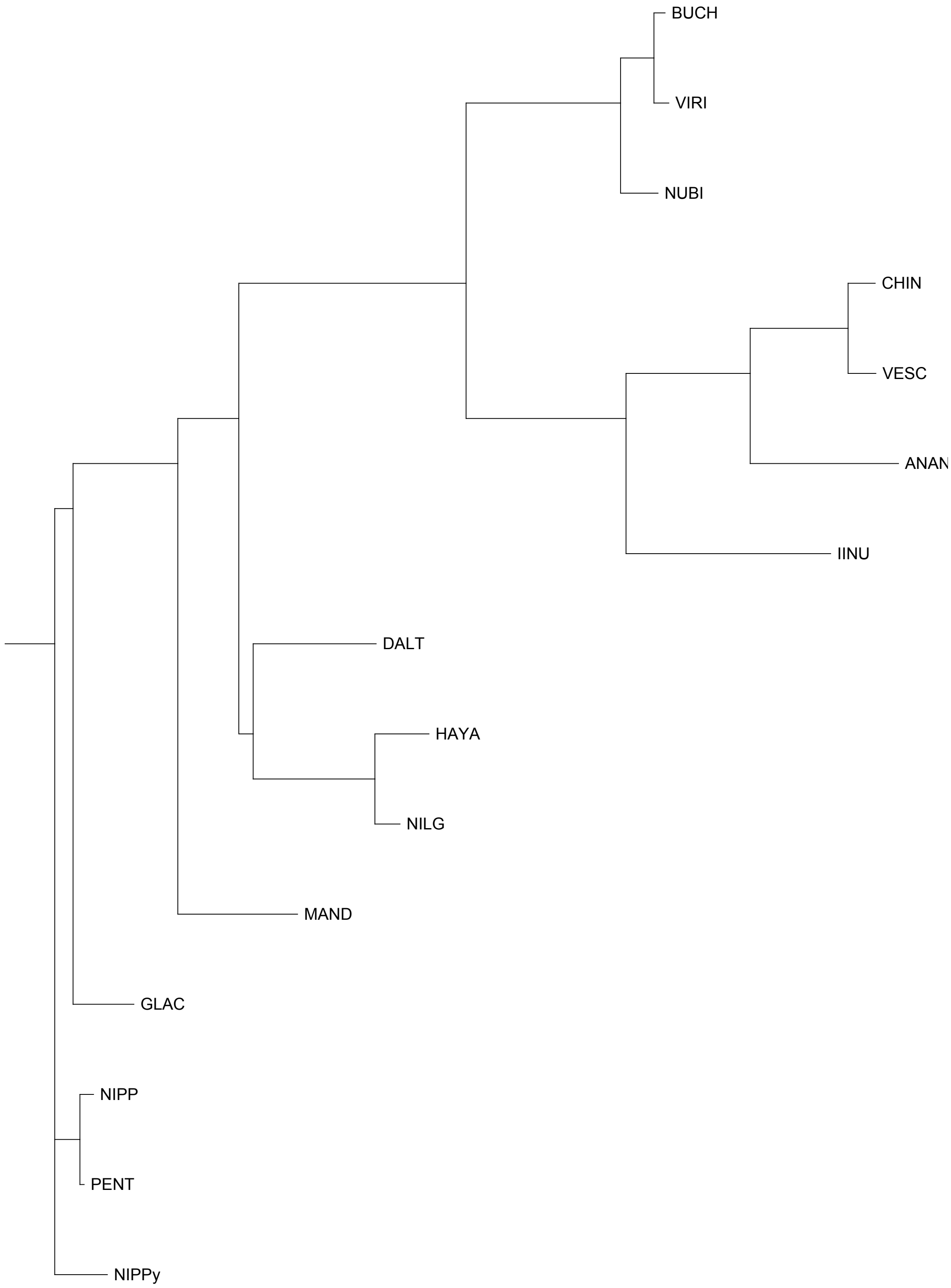


FAN_r2.3ch1X2a

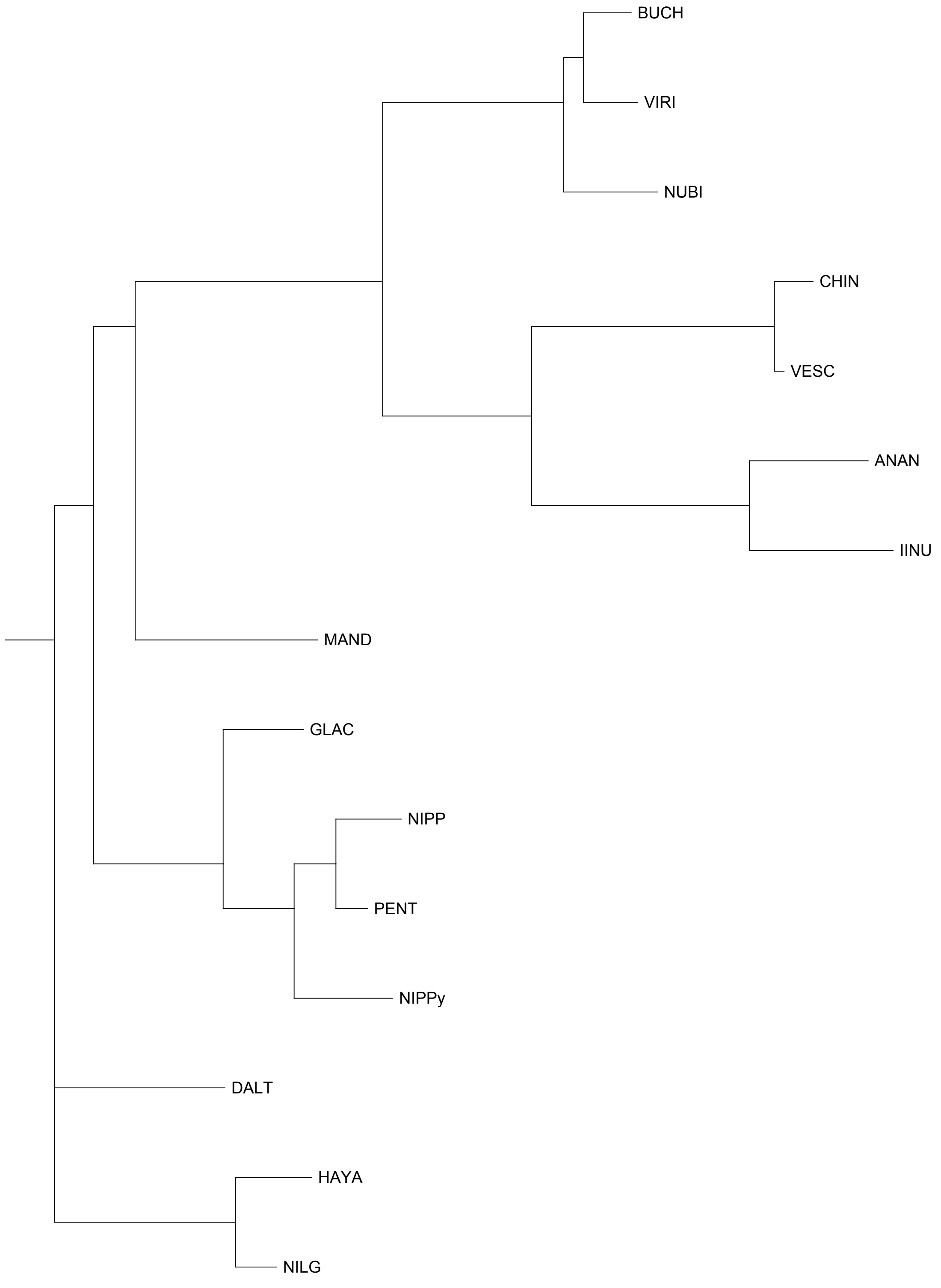


FAN_r2.3ch1X2b

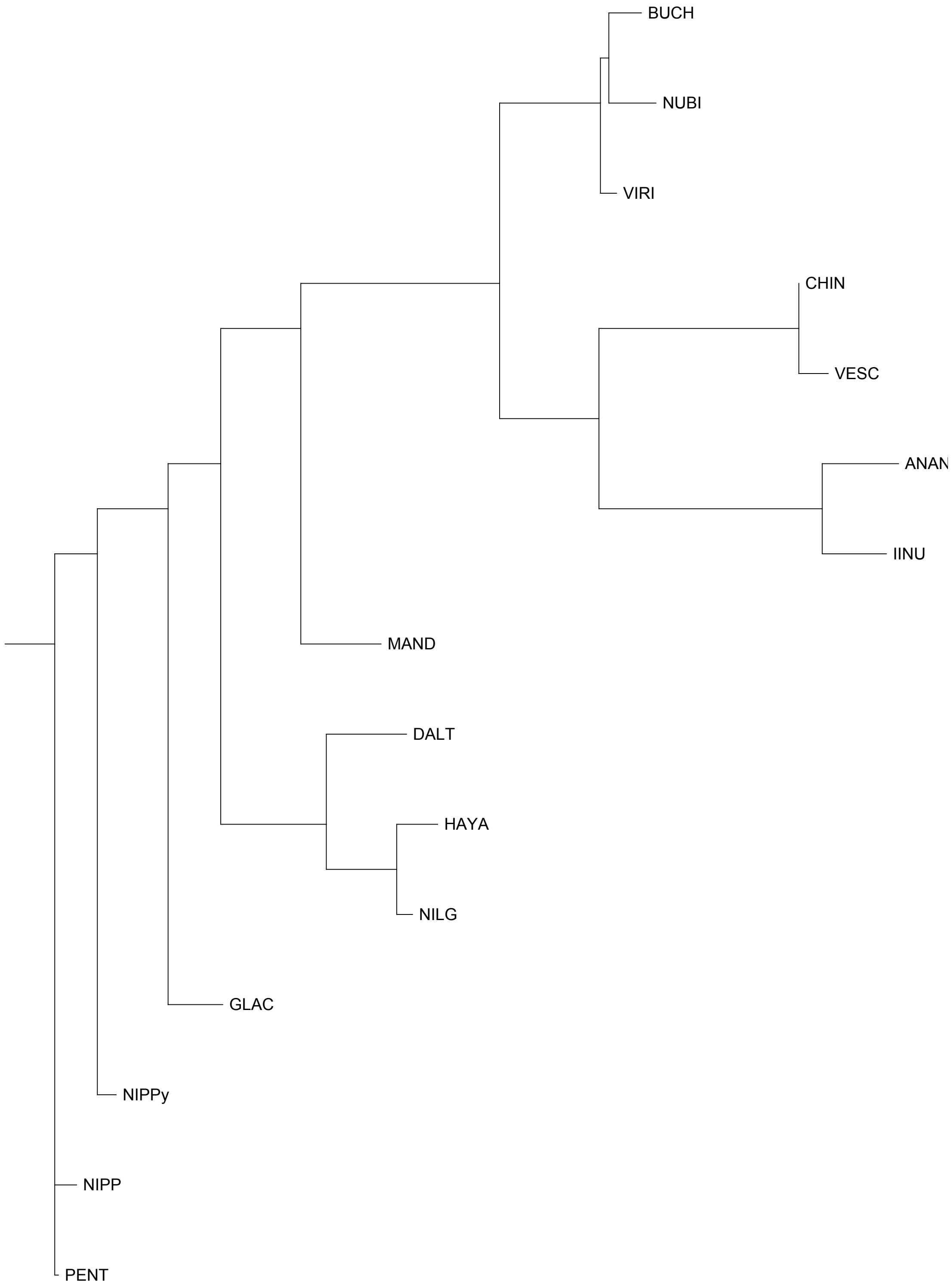


FAN_r2.3ch1X3a

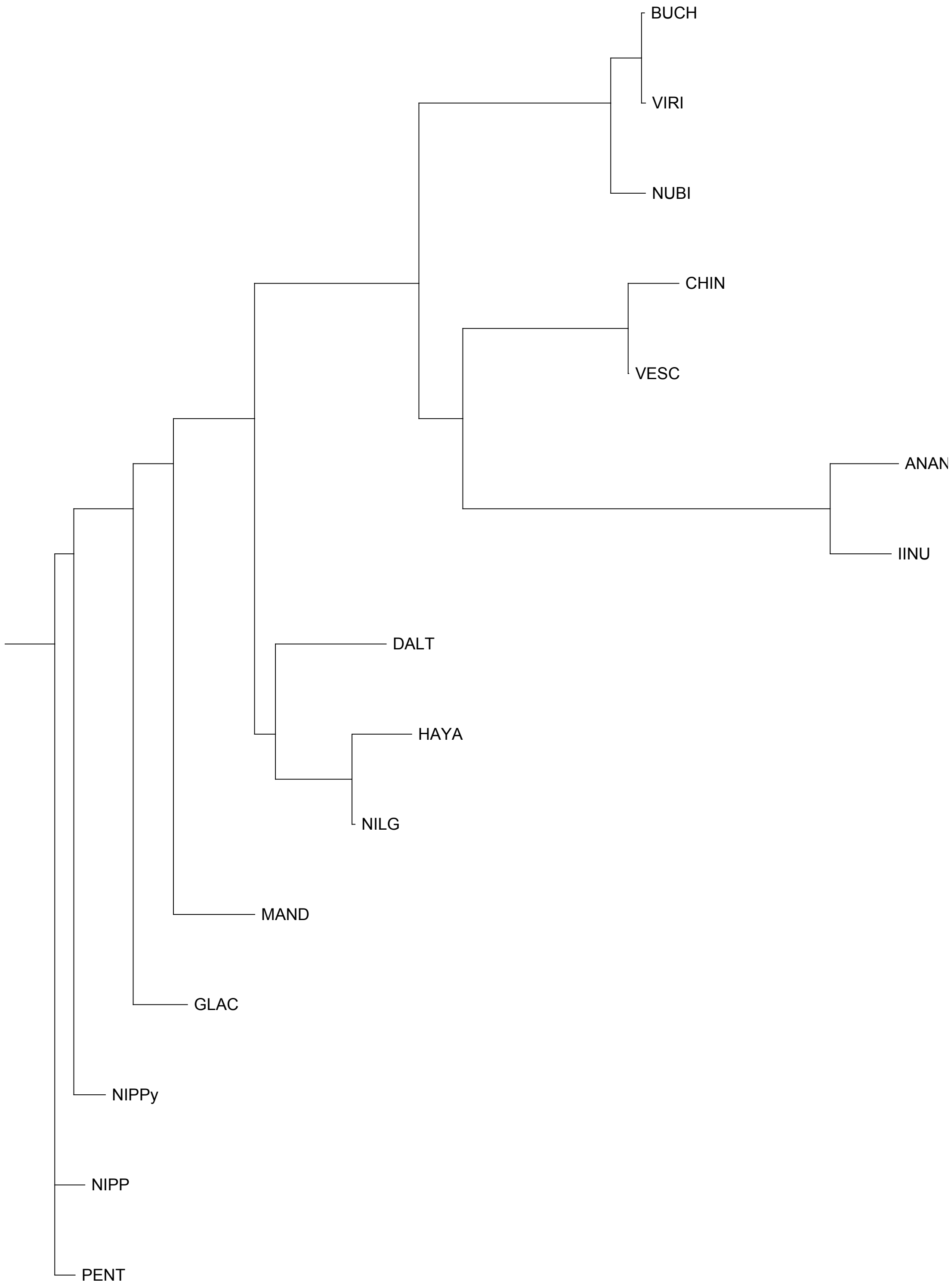


FAN_r2.3ch1X3b

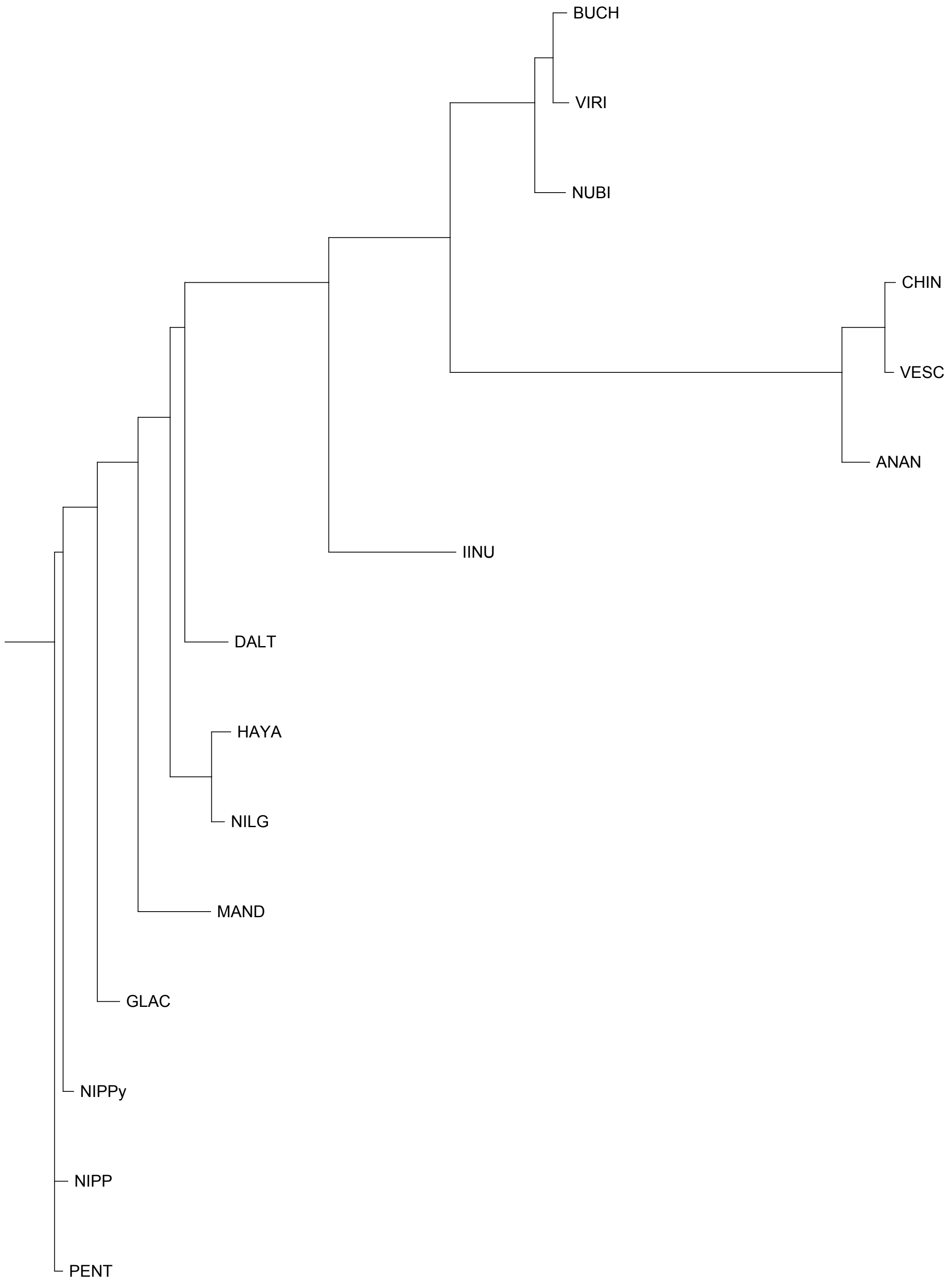


FAN_r2.3ch2Ava

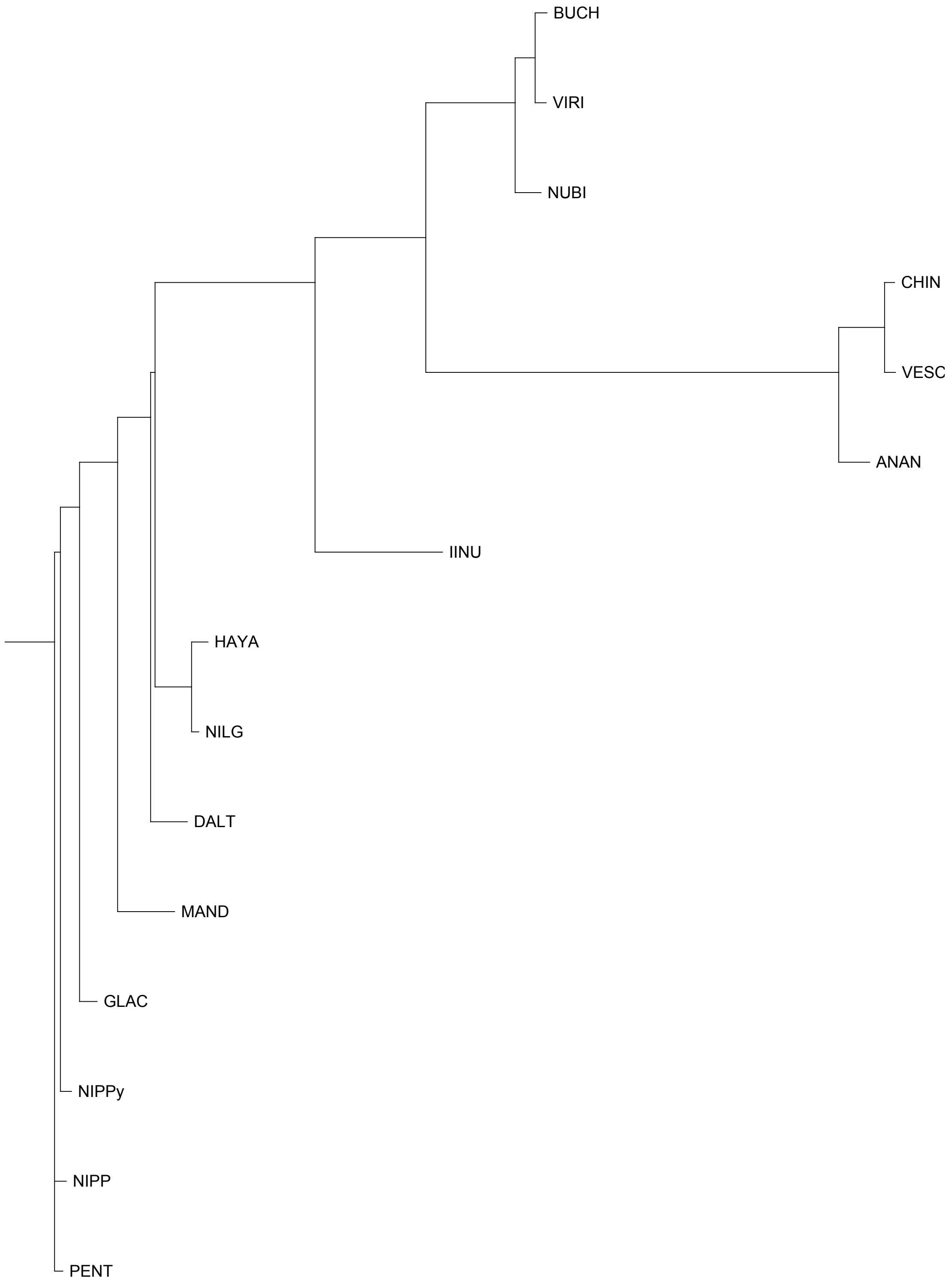


FAN_r2.3ch2Avb

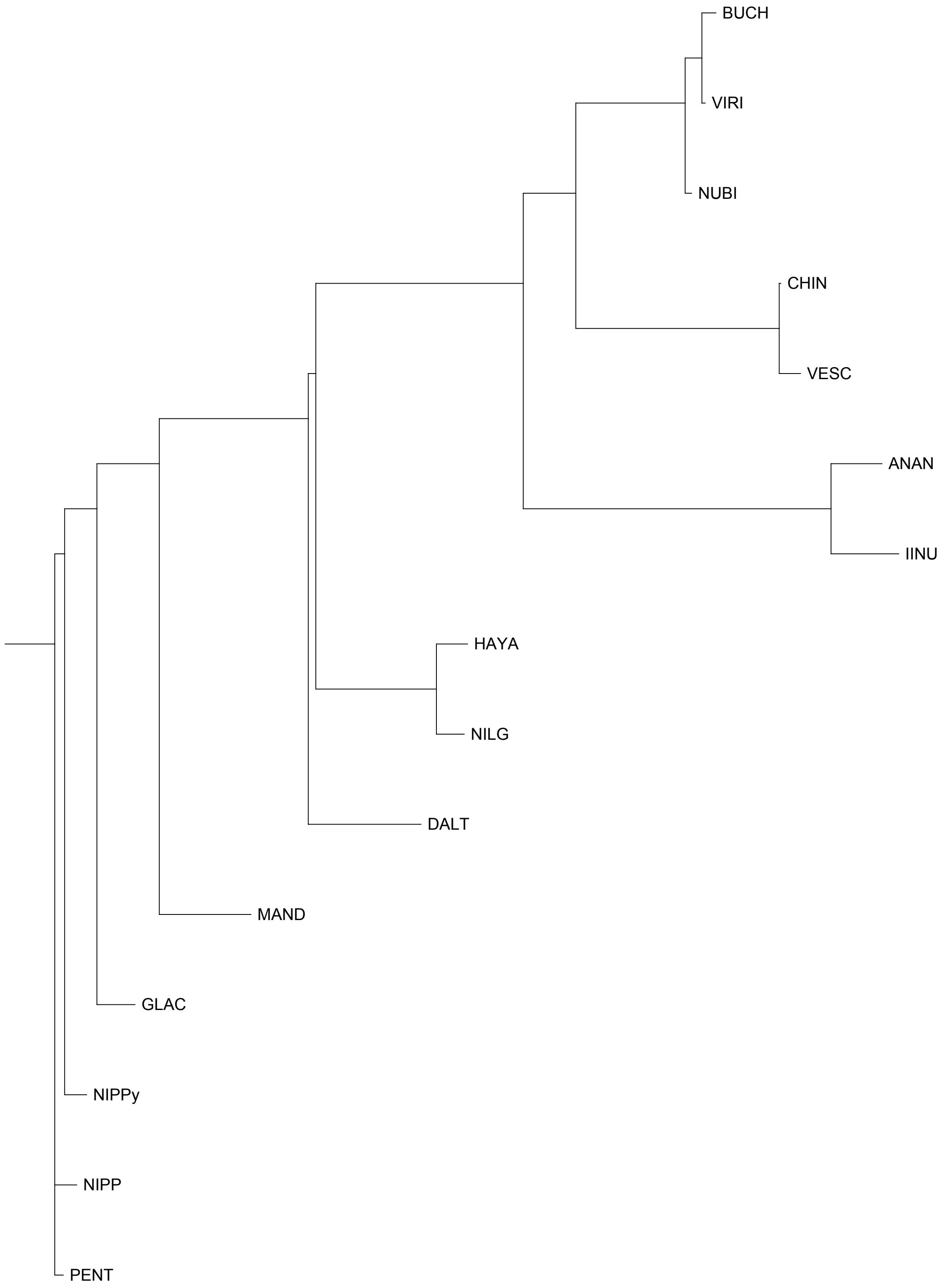


FAN_r2.3ch2Bia

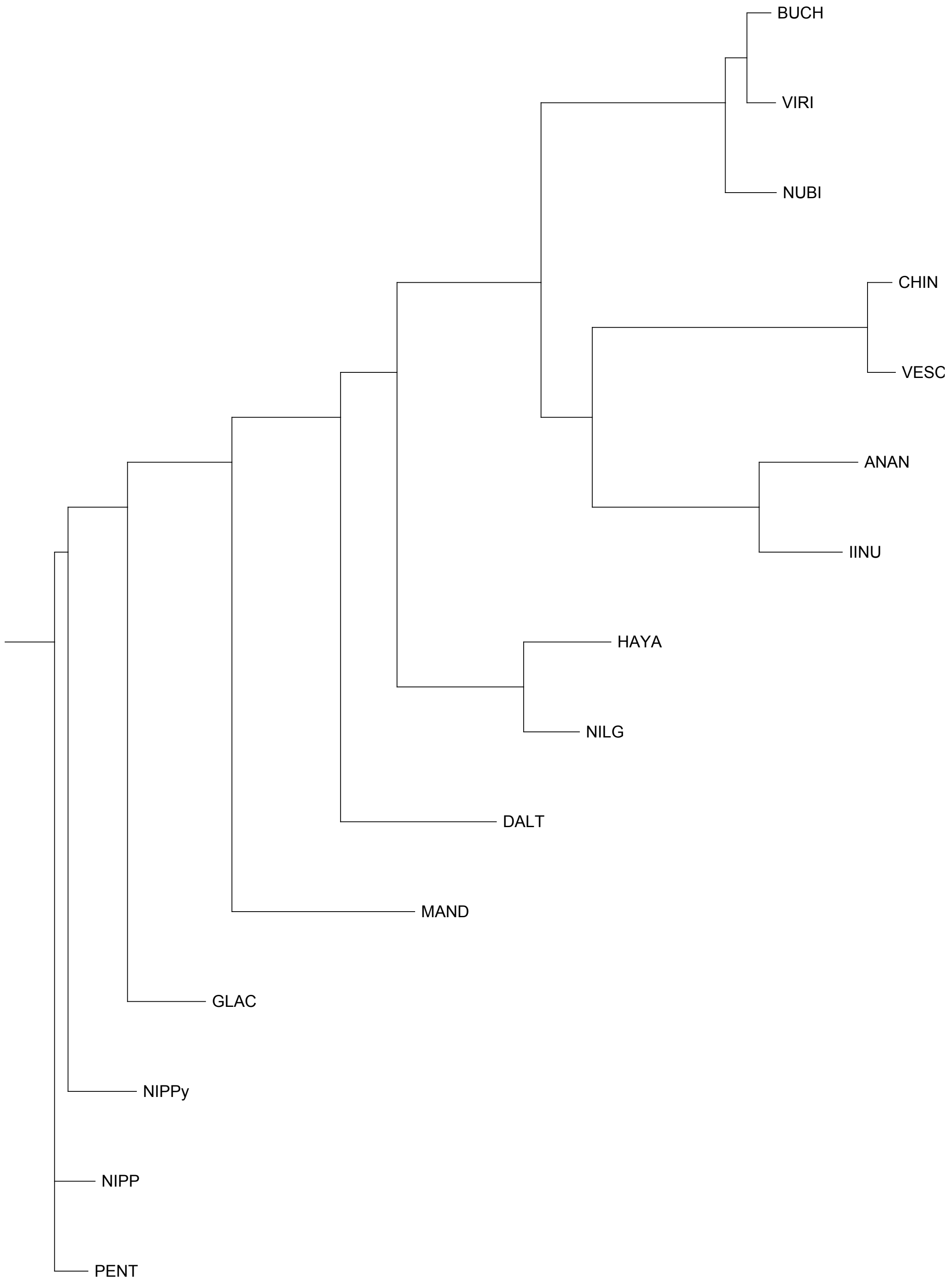


FAN_r2.3ch2Bib

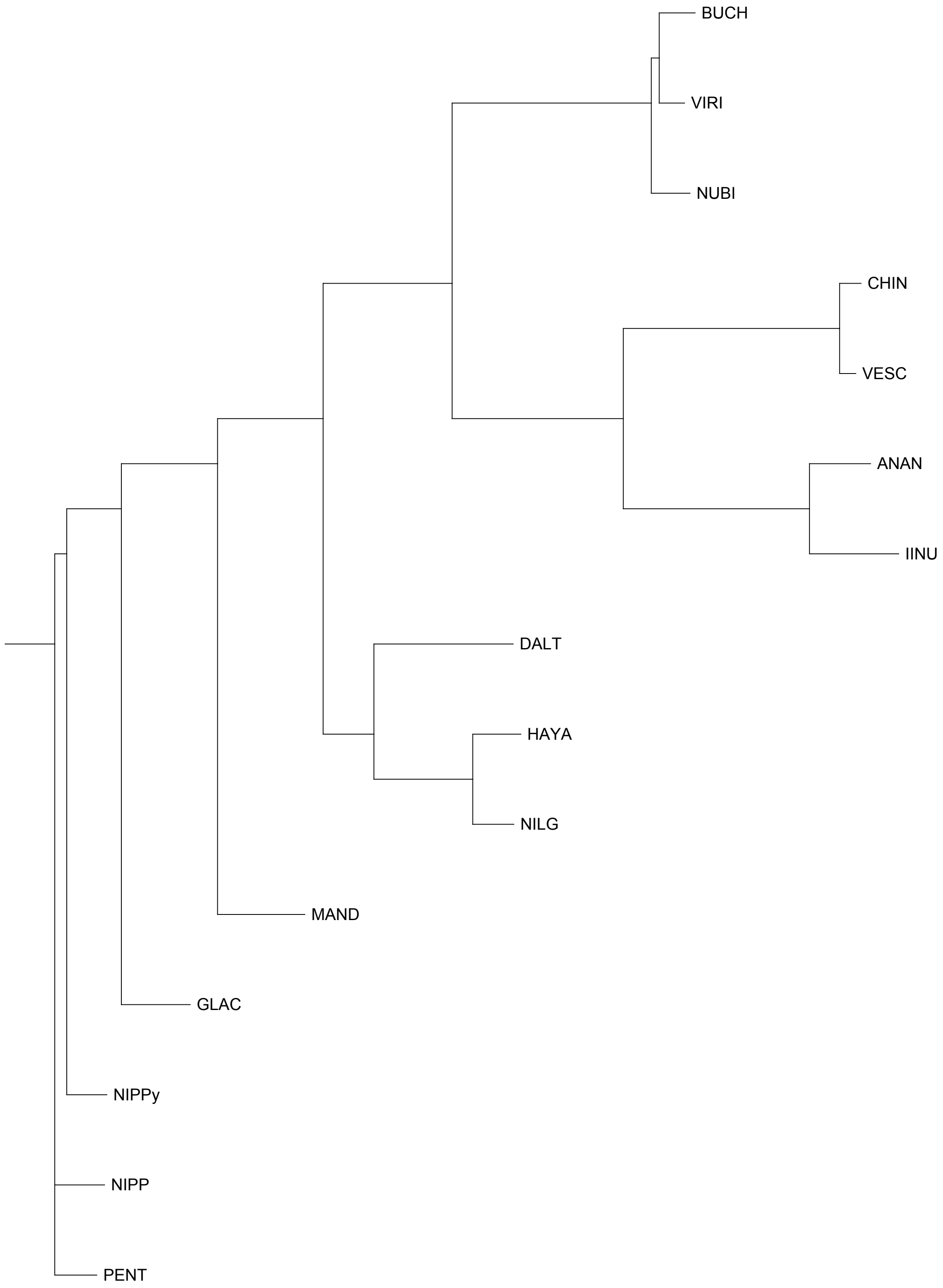


FAN_r2.3ch2X1a

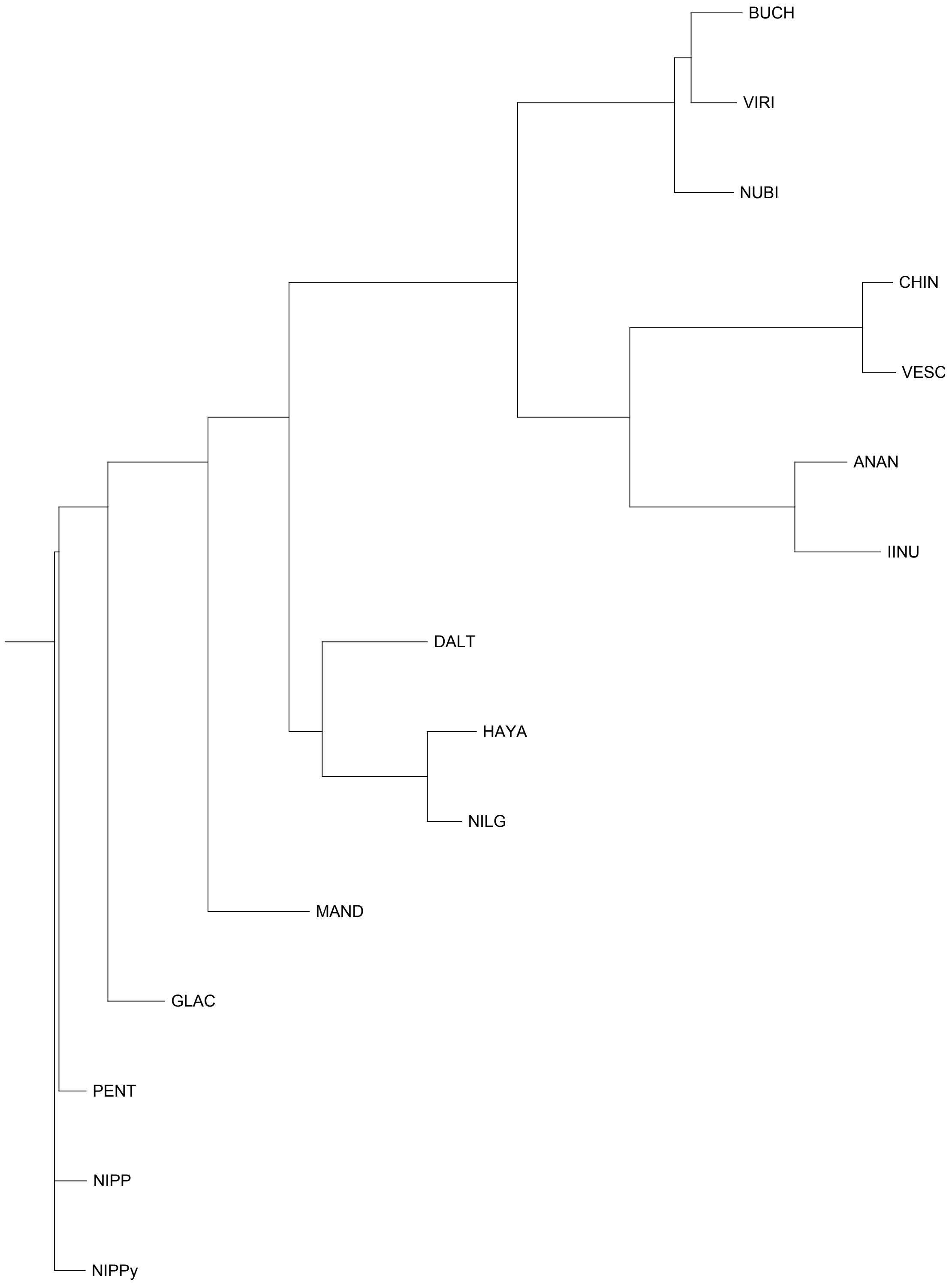


FAN_r2.3ch2X1b

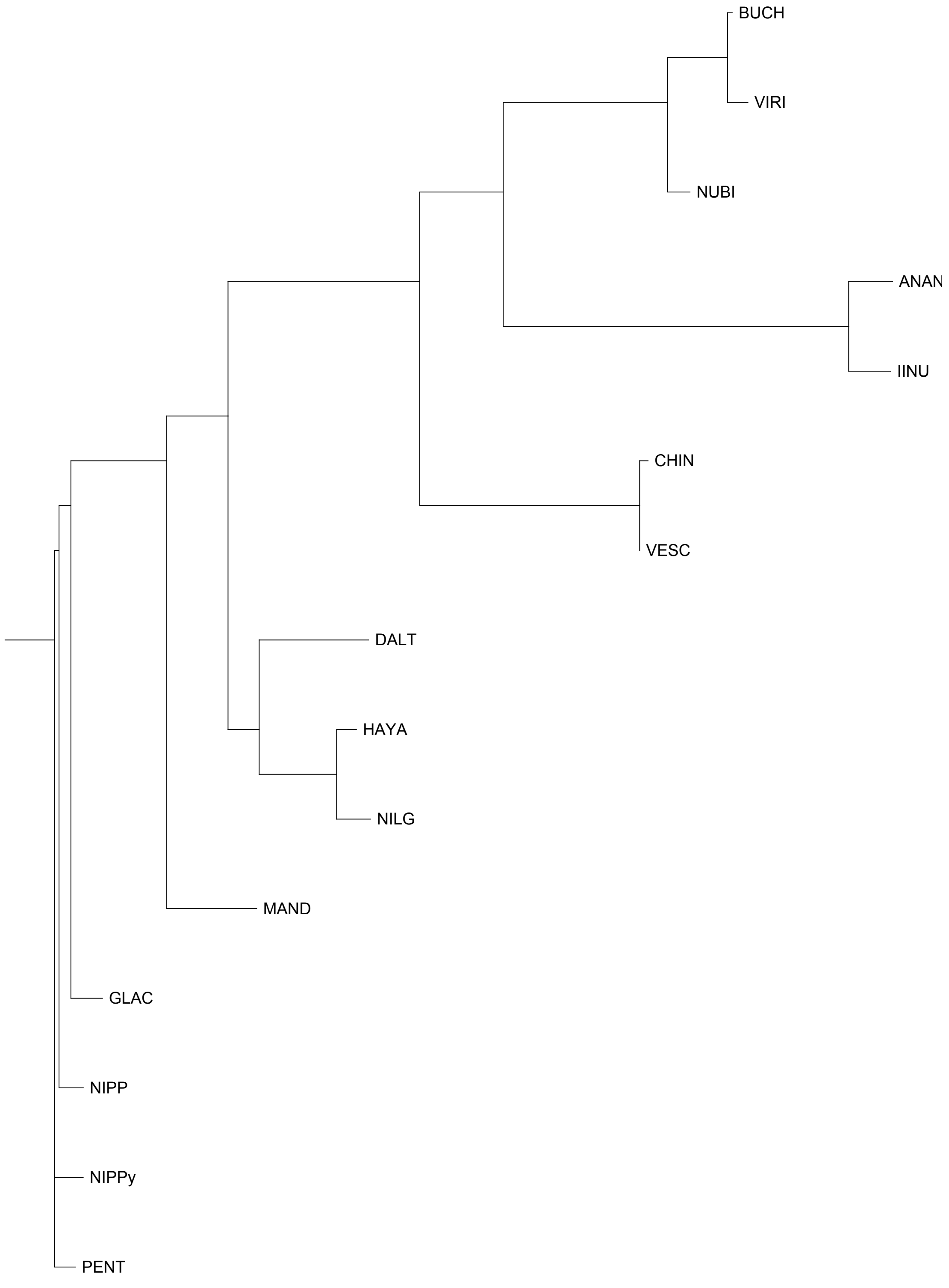


FAN_r2.3ch2X2a

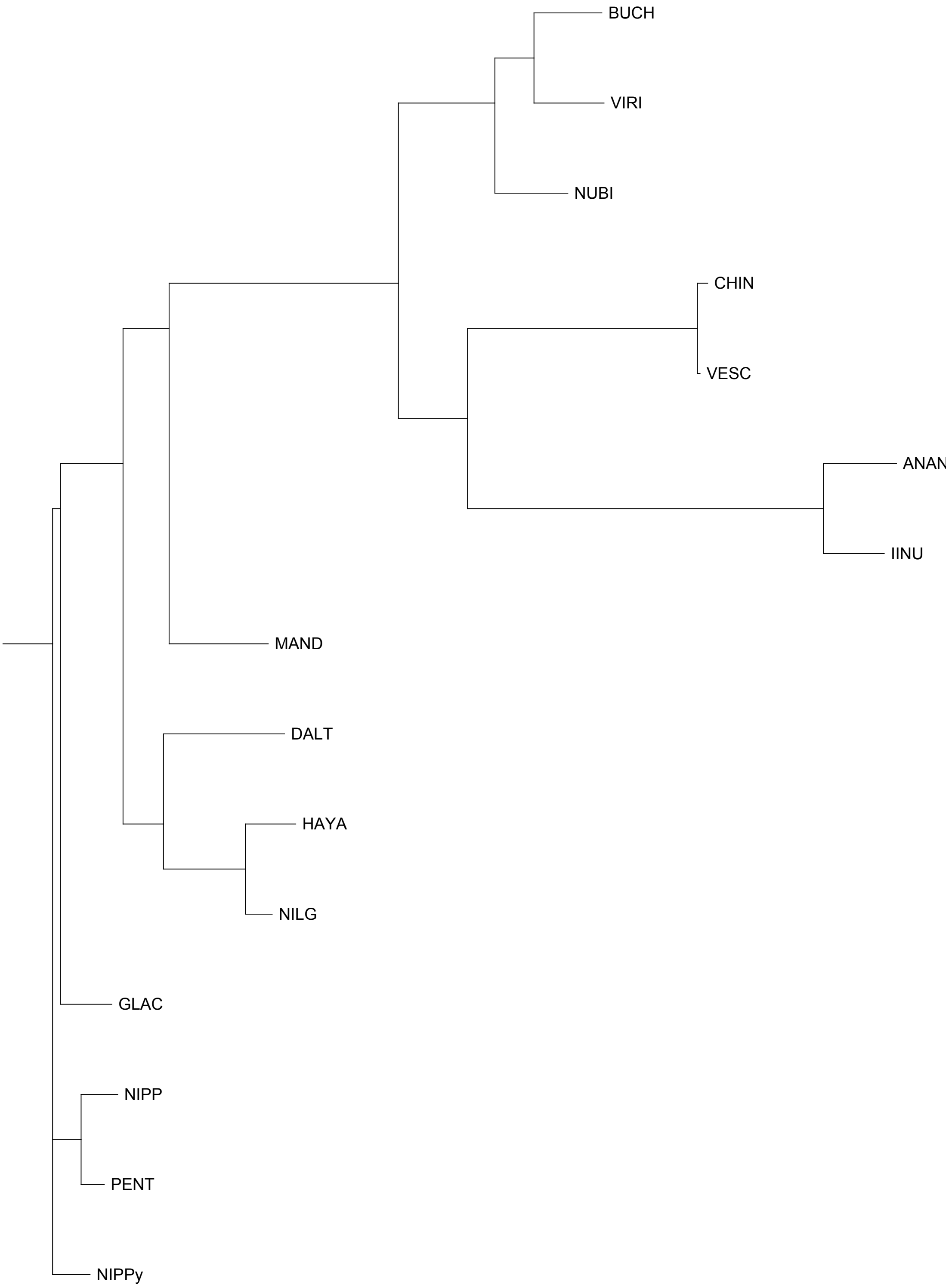


FAN_r2.3ch2X2b

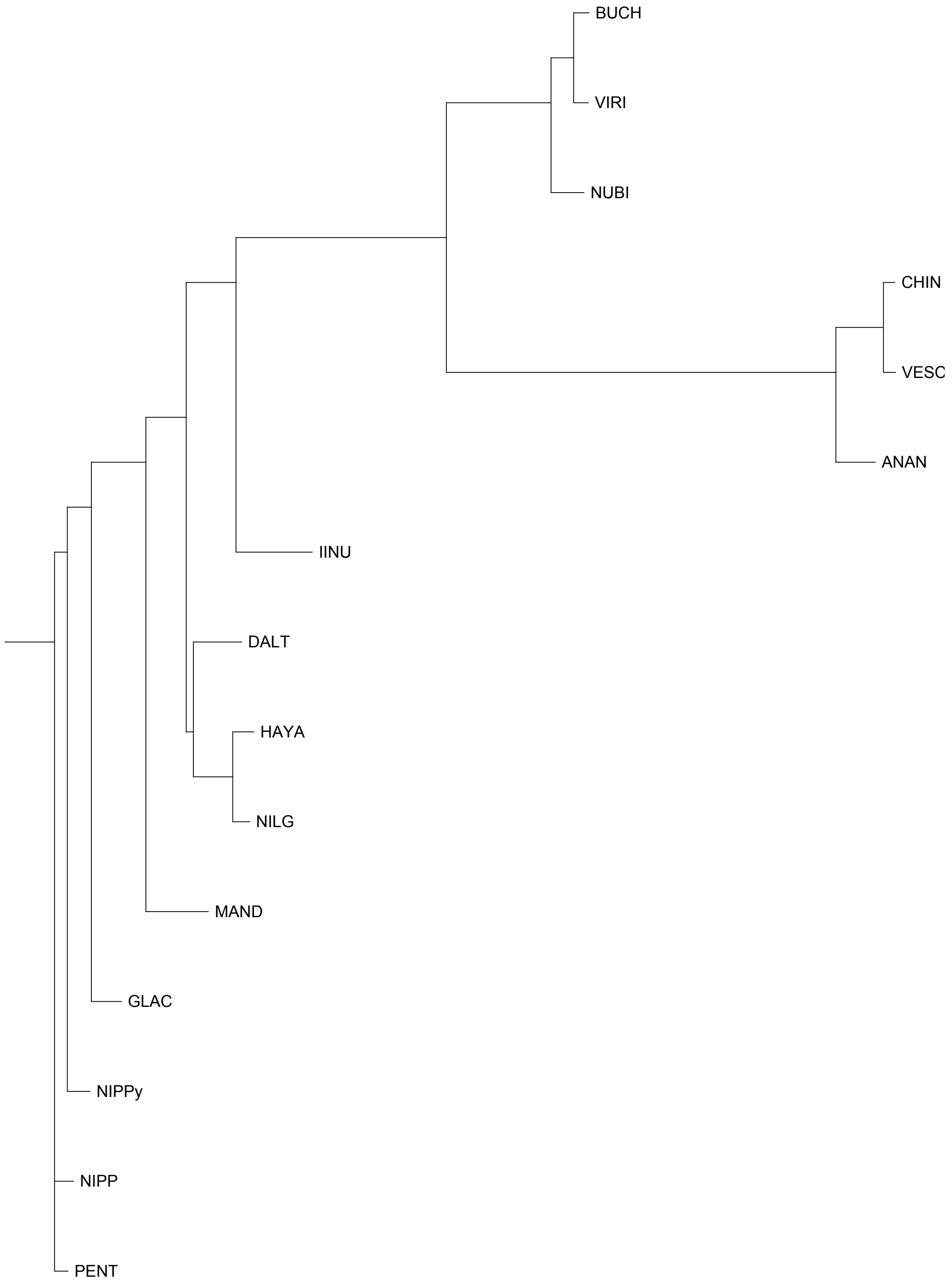


FAN_r2.3ch3Ava

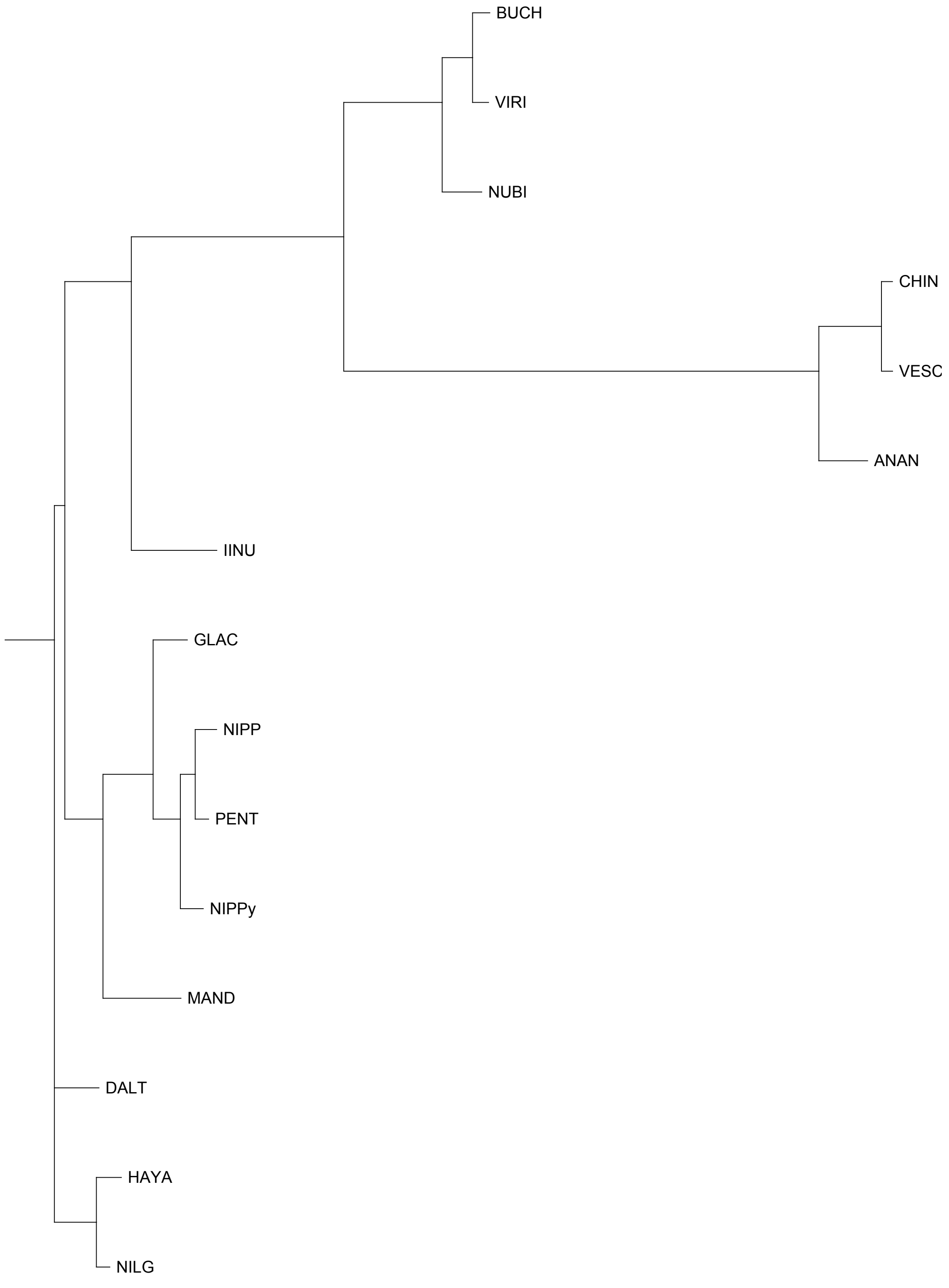


FAN_r2.3ch3Avb

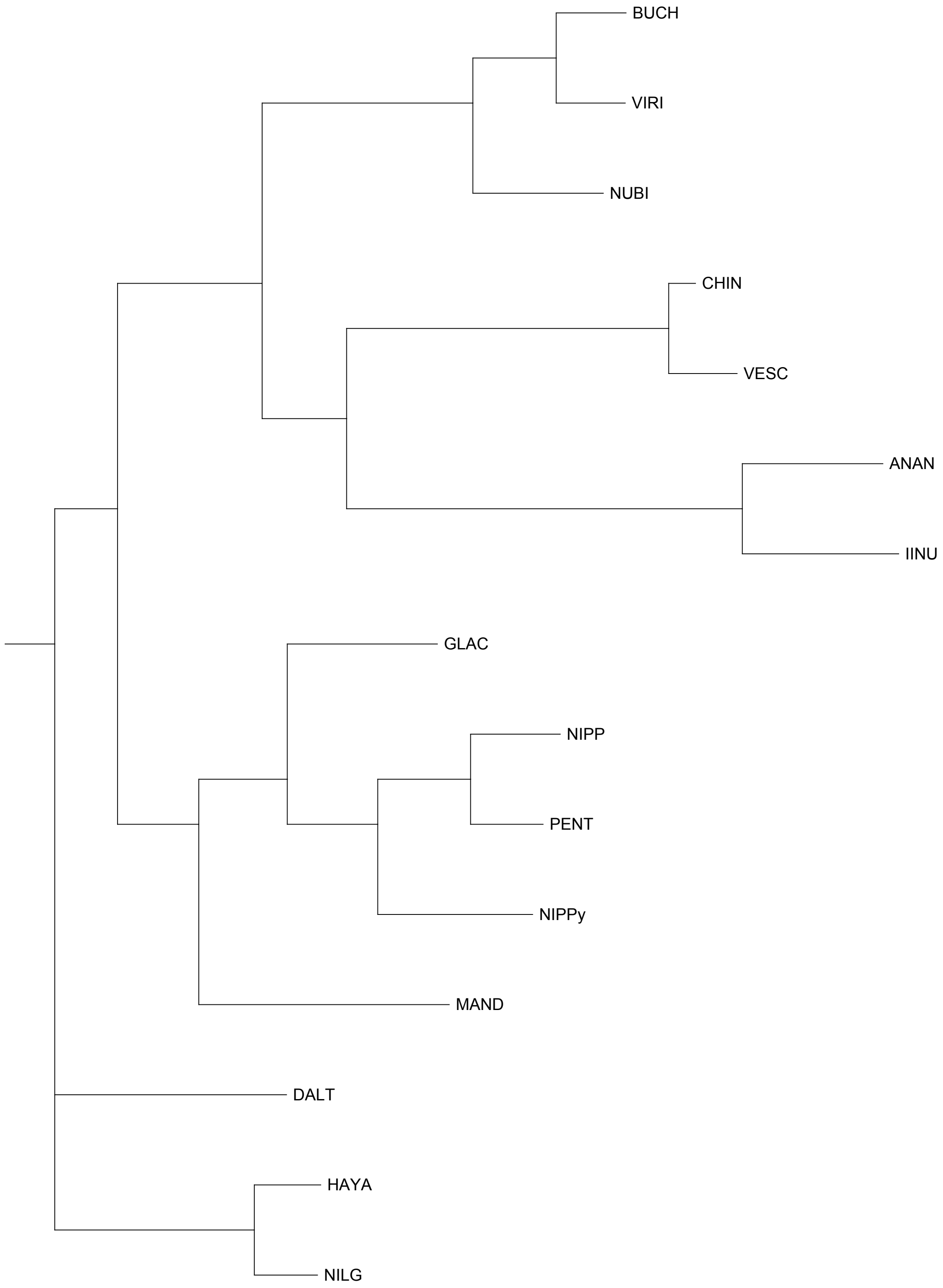


FAN_r2.3ch3Bia

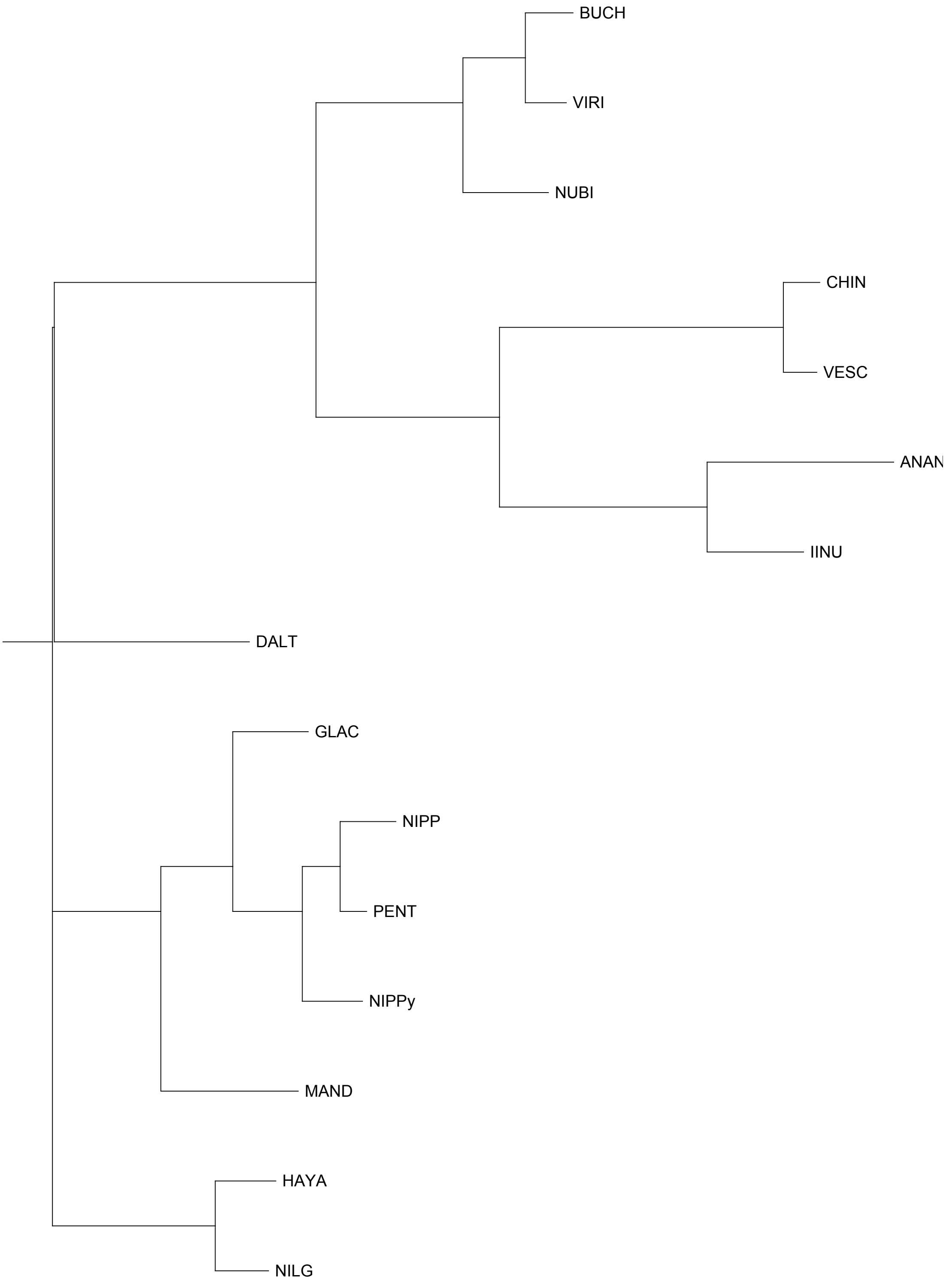


FAN_r2.3ch3Bib

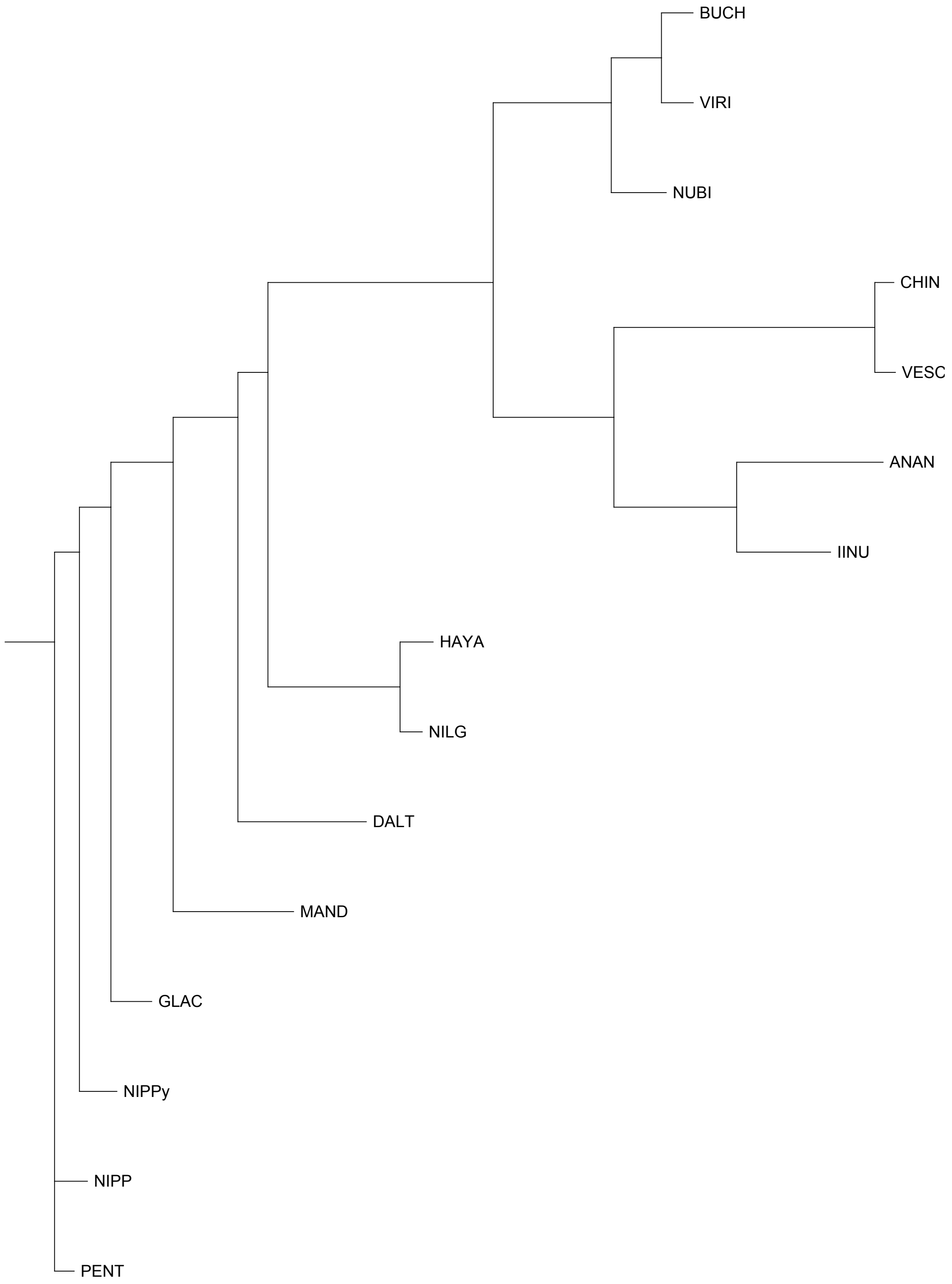


FAN_r2.3ch3X1a

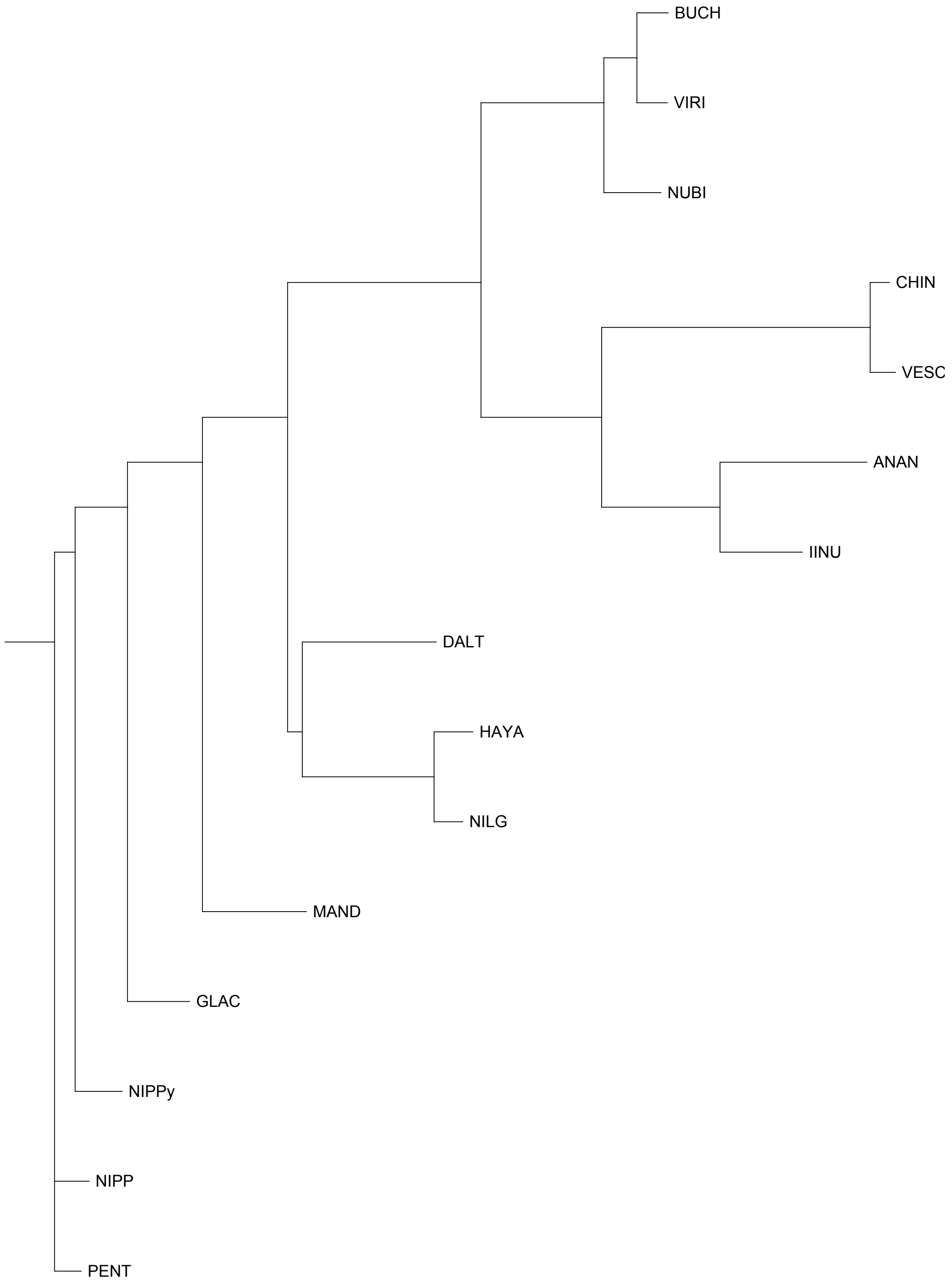


FAN_r2.3ch3X1b

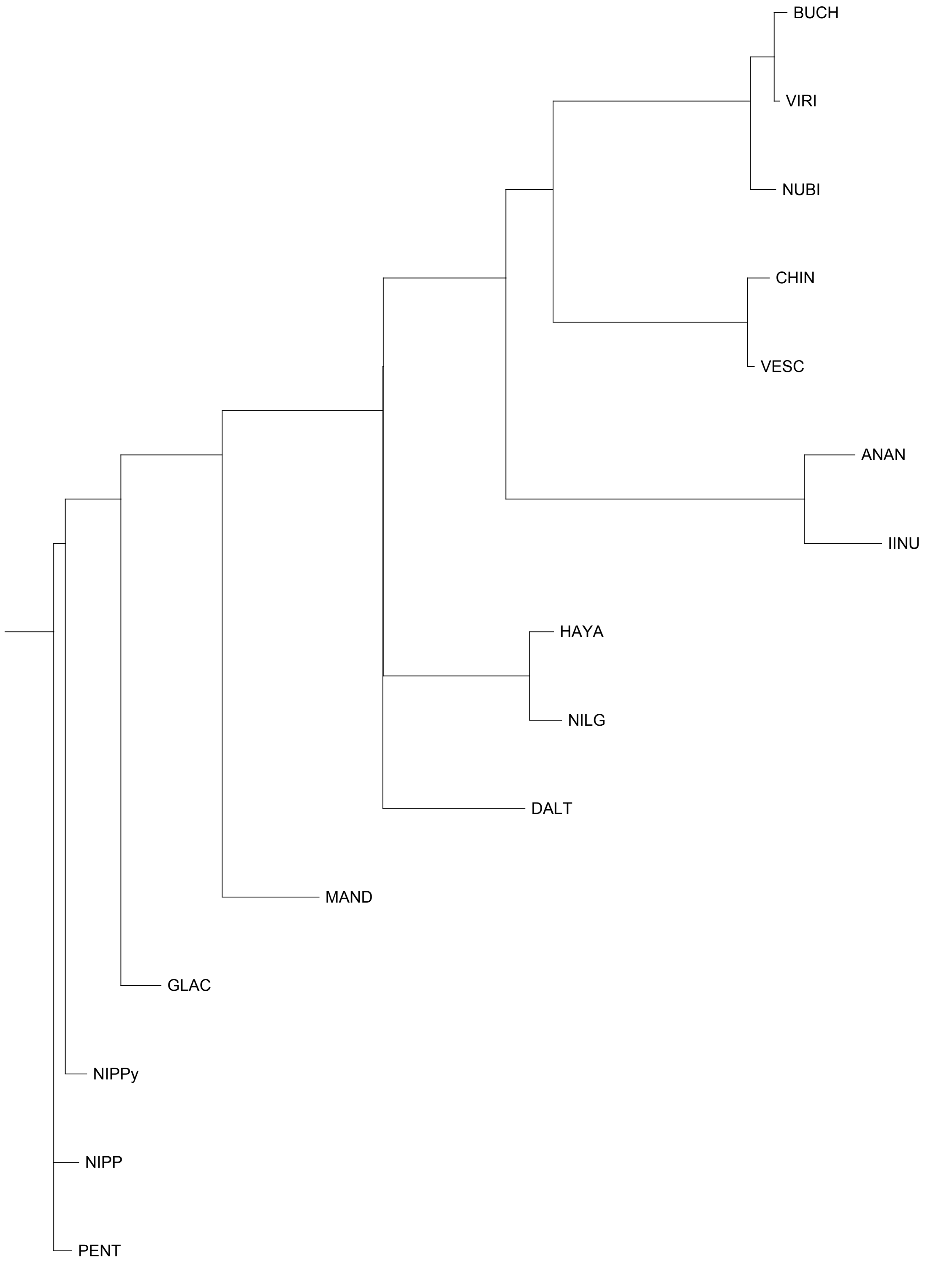


FAN_r2.3ch3X2a

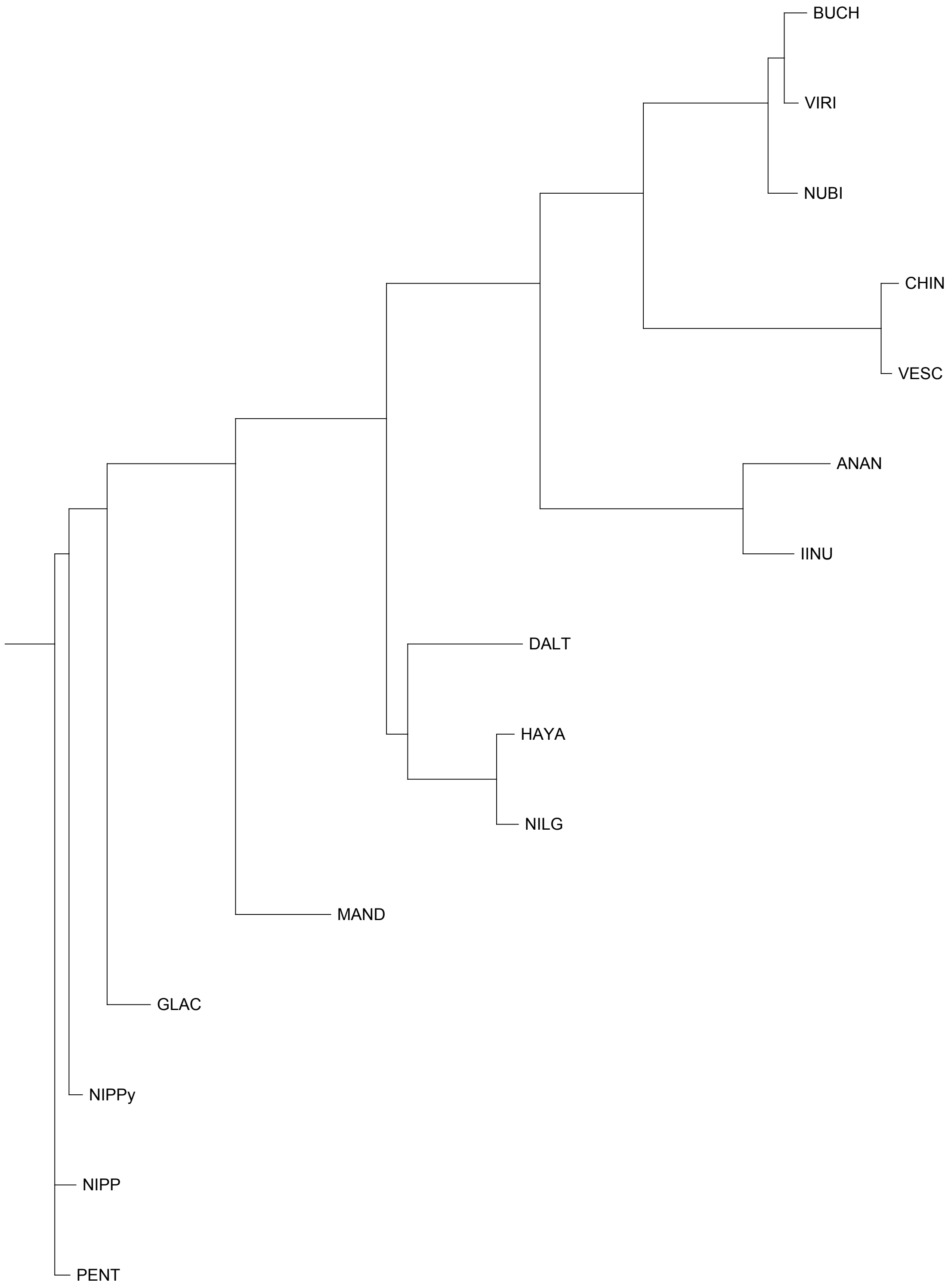


FAN_r2.3ch3X2b

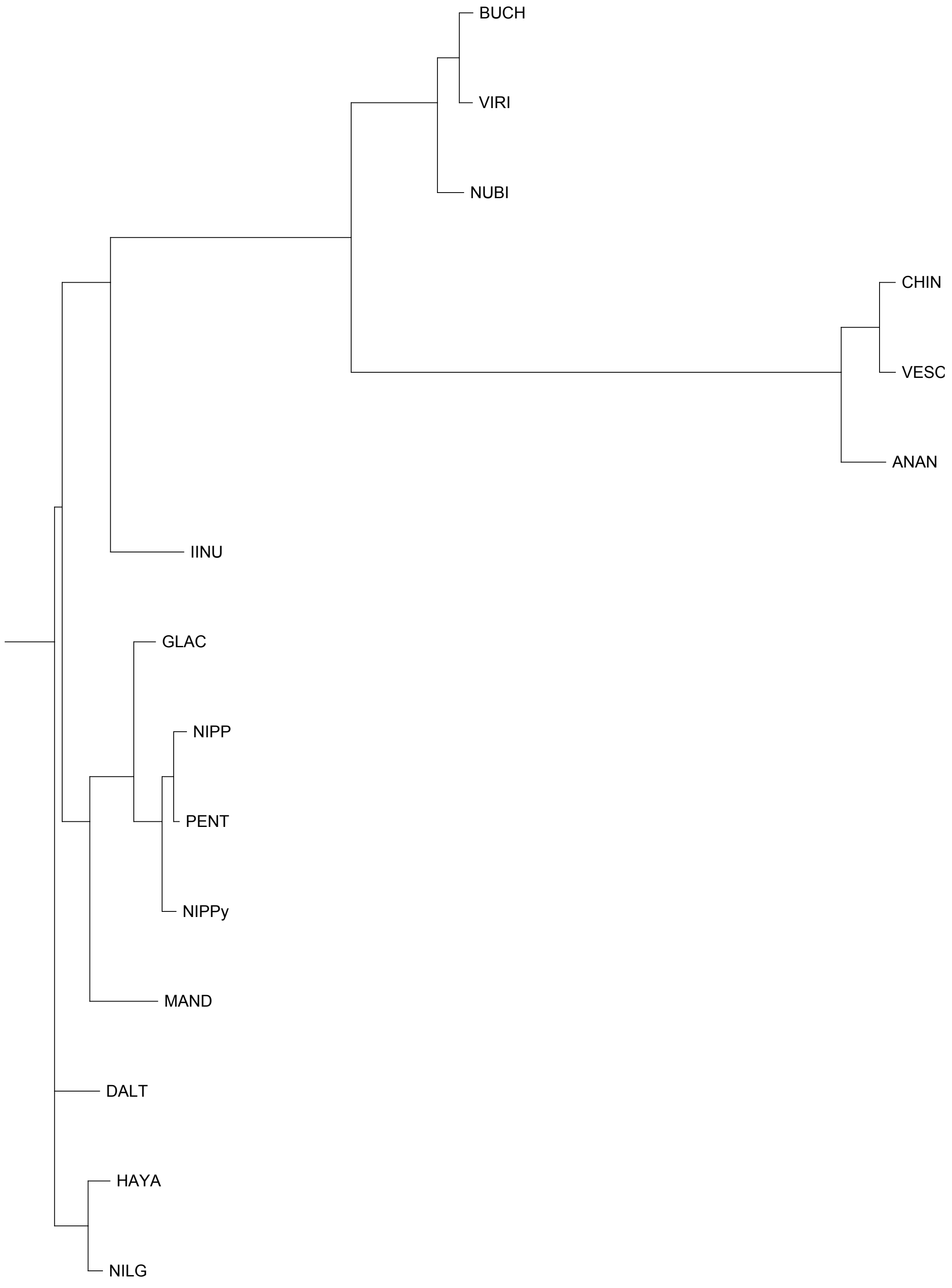


FAN_r2.3ch4Ava

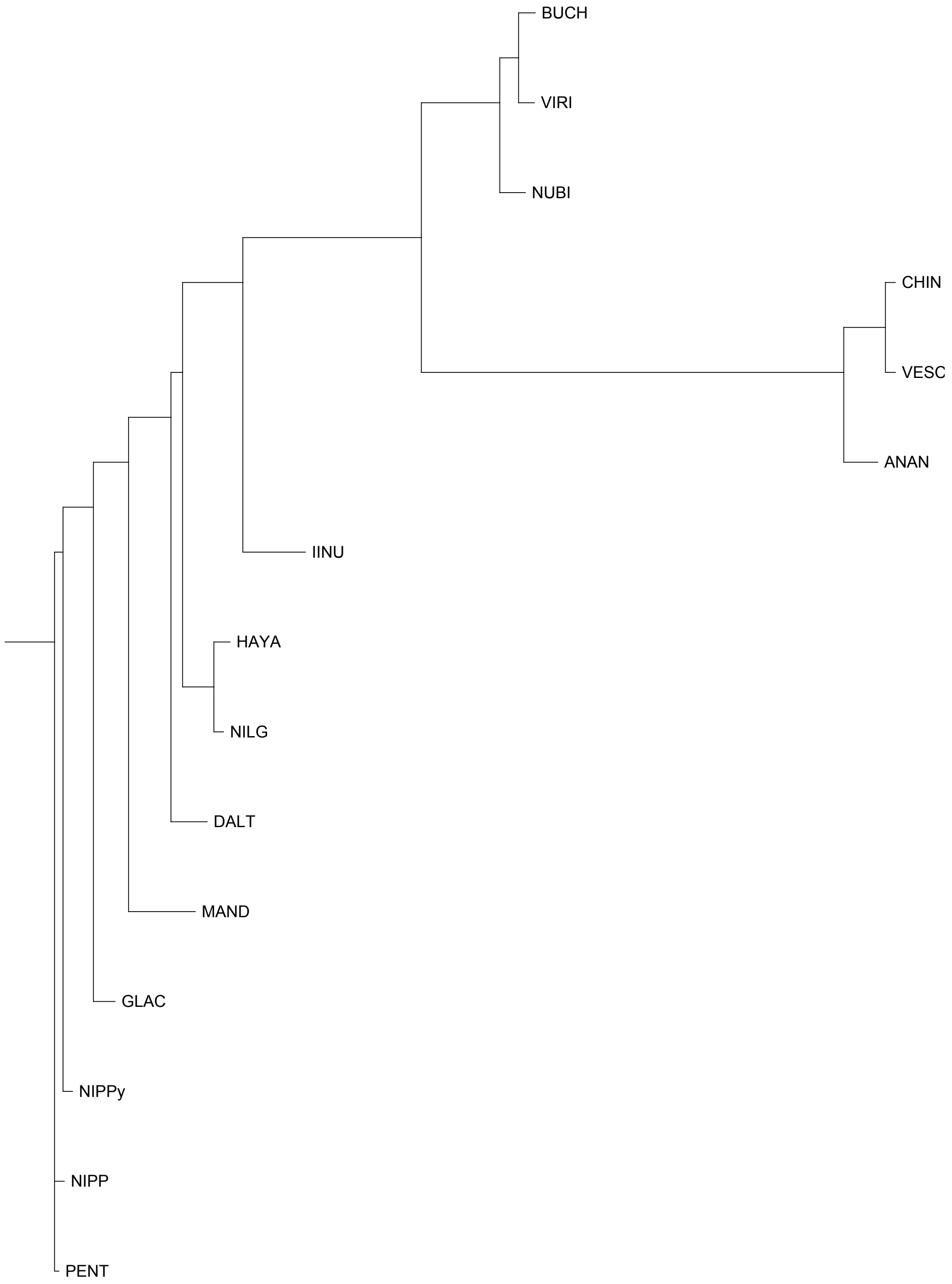


FAN_r2.3ch4Avb

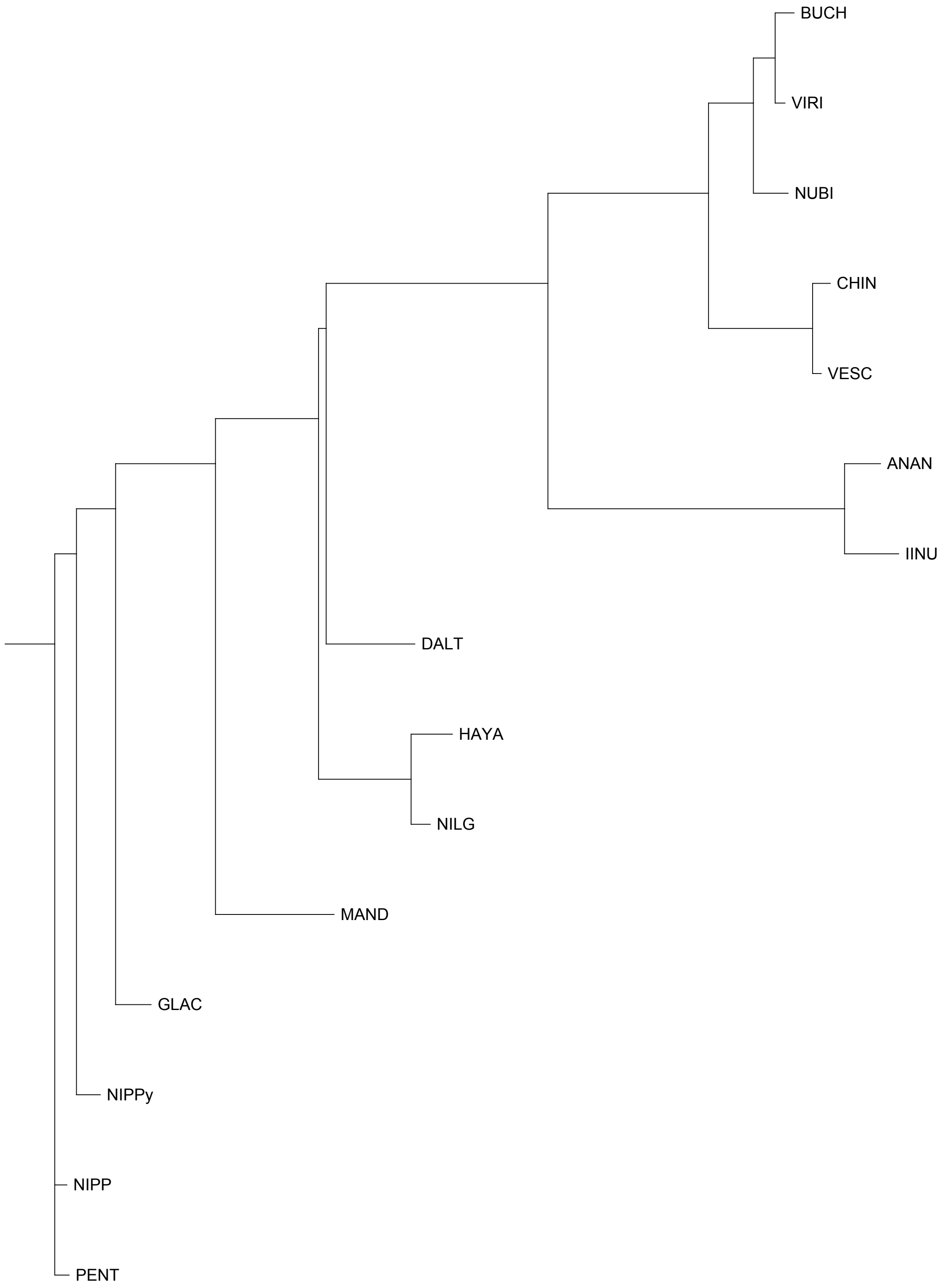


FAN_r2.3ch4Bia

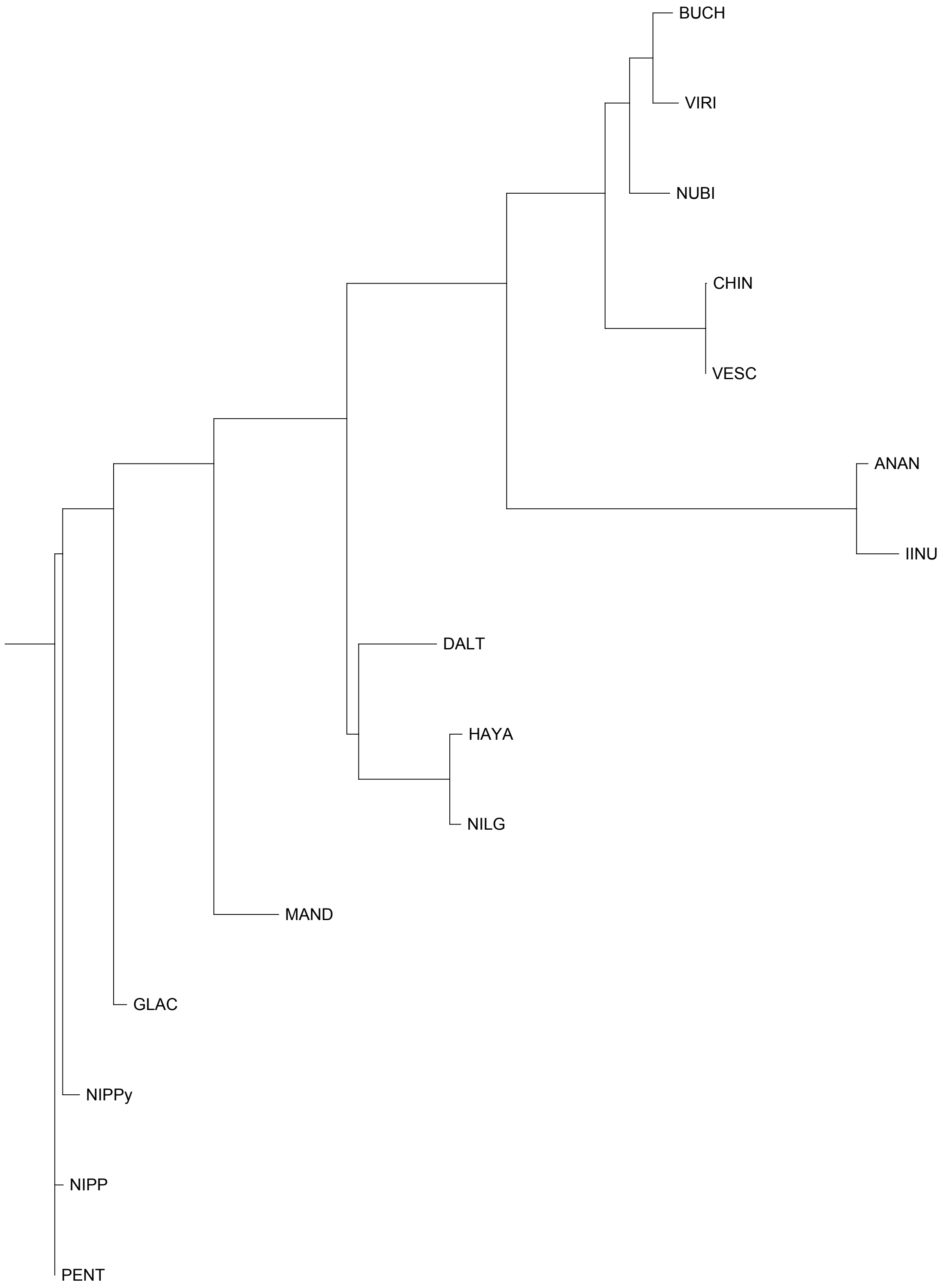


FAN_r2.3ch4Bib

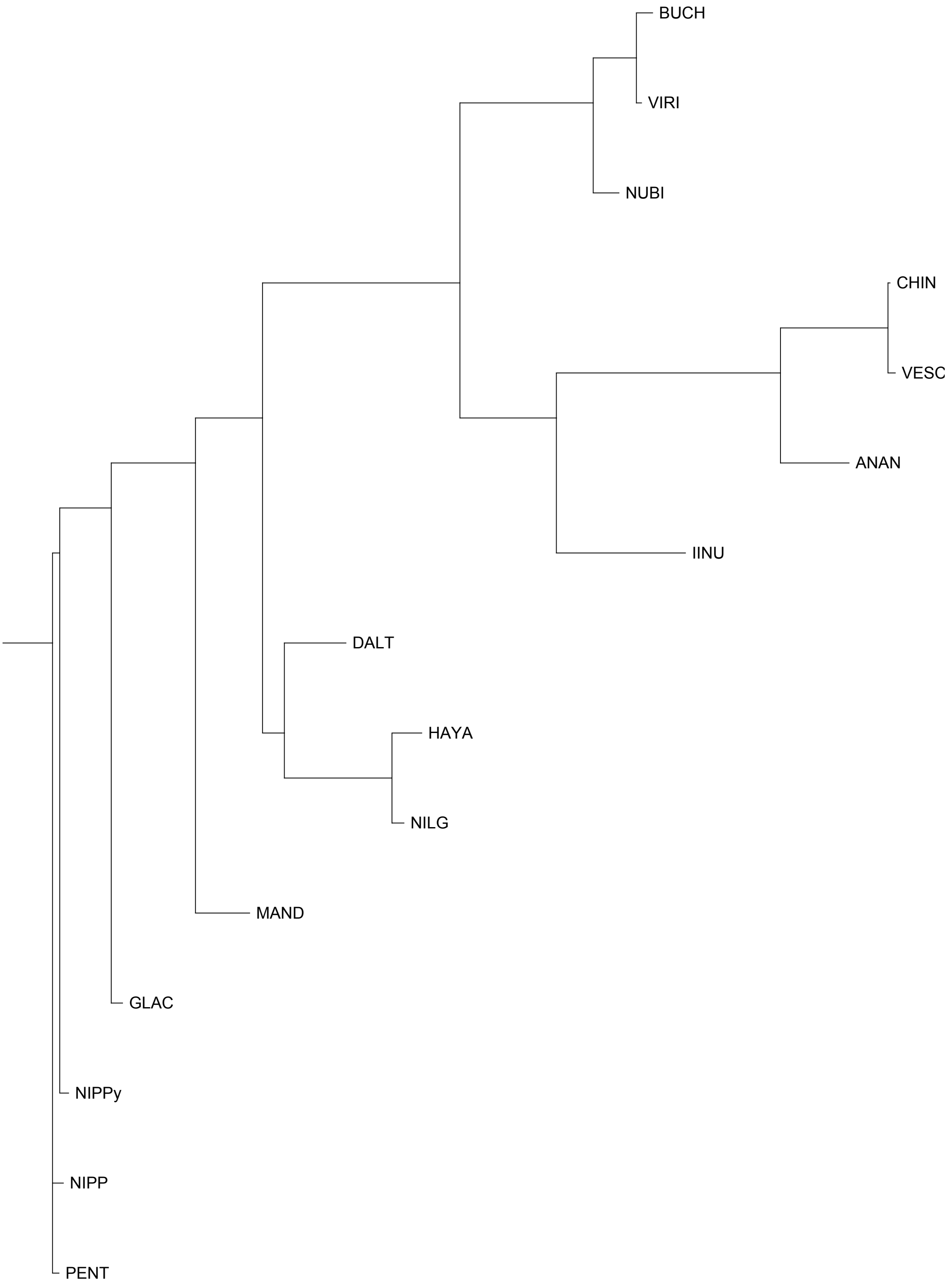


FAN_r2.3ch4X1a

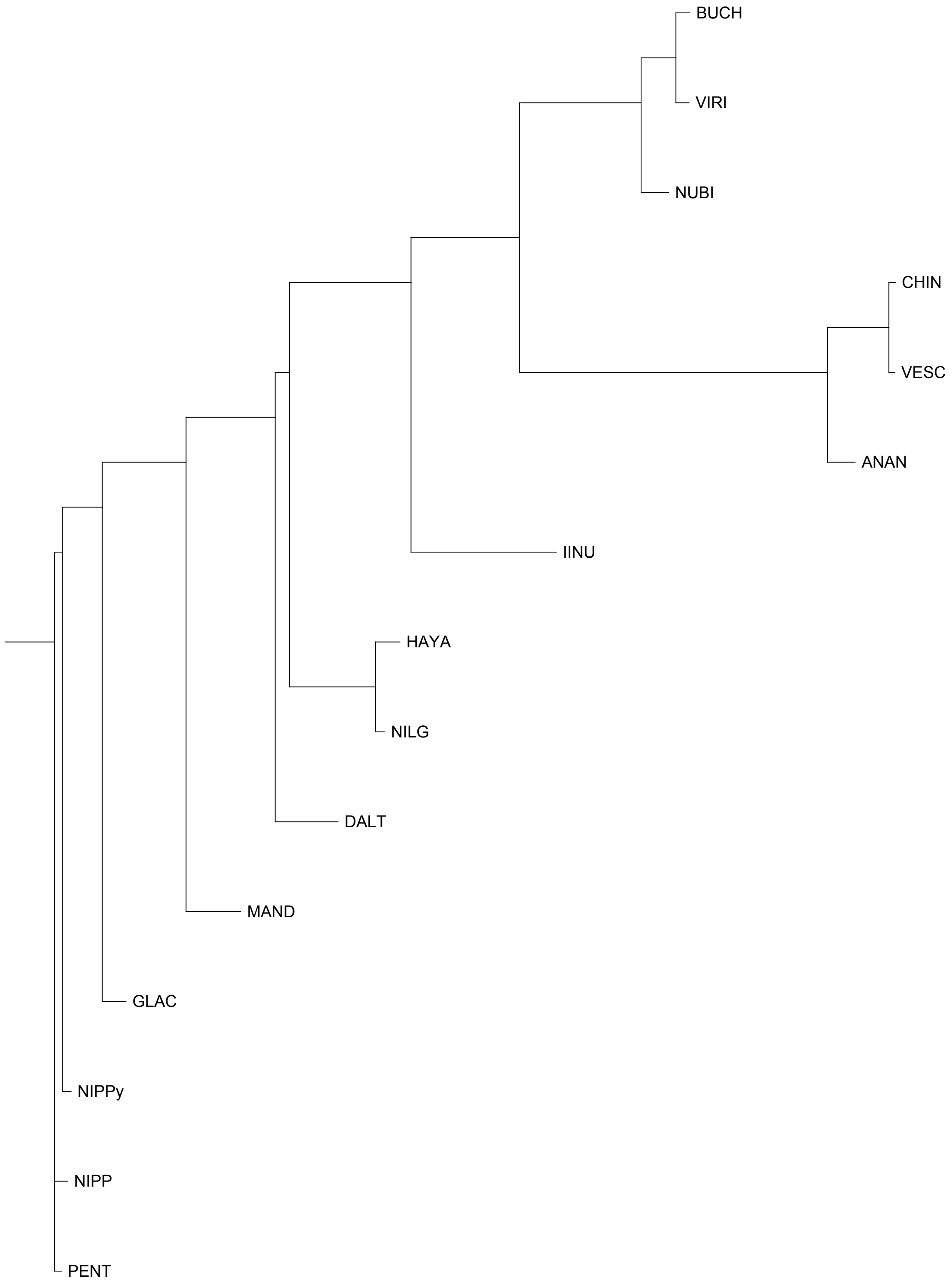


FAN_r2.3ch4X1b

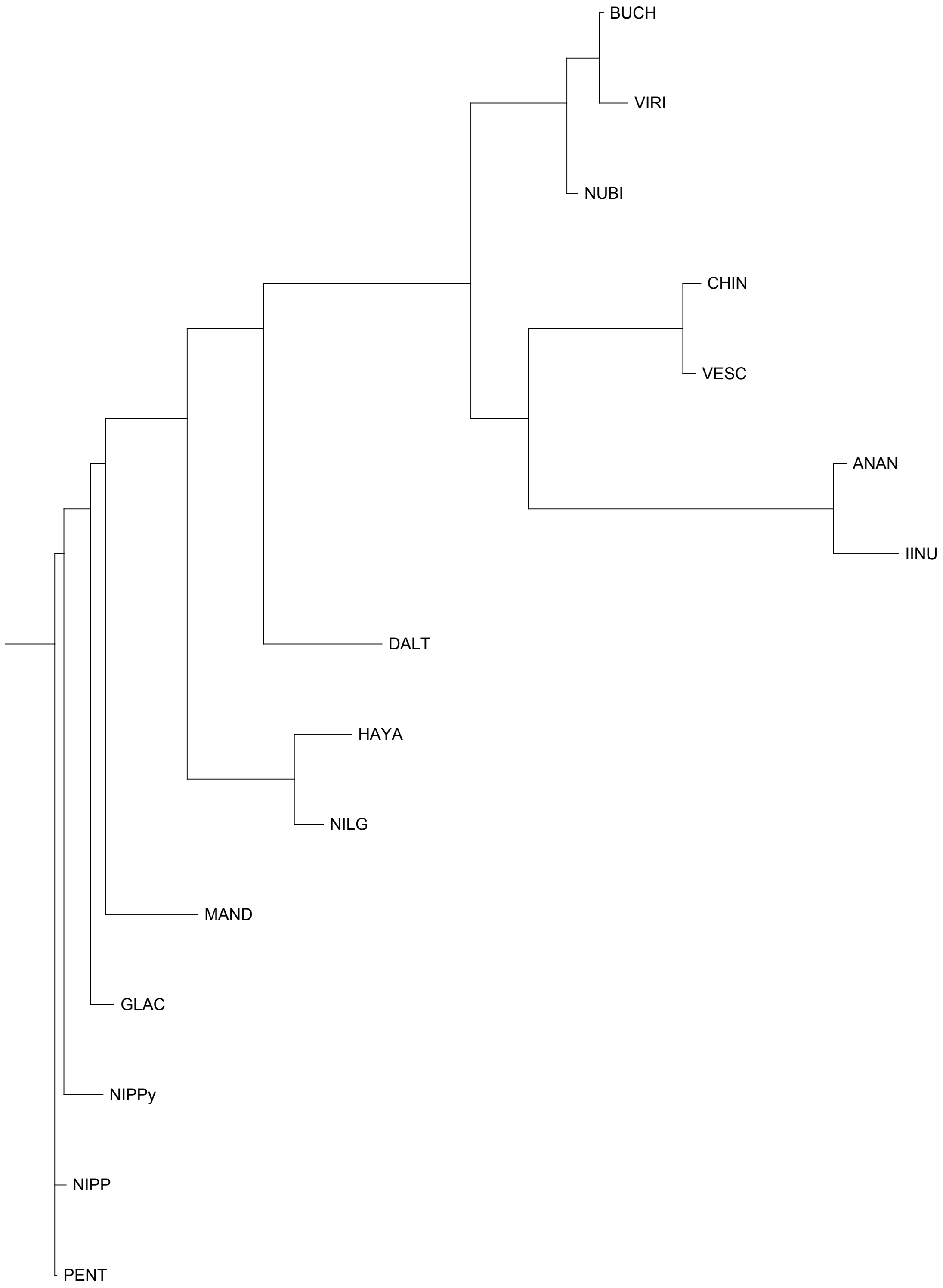


FAN_r2.3ch4X2a

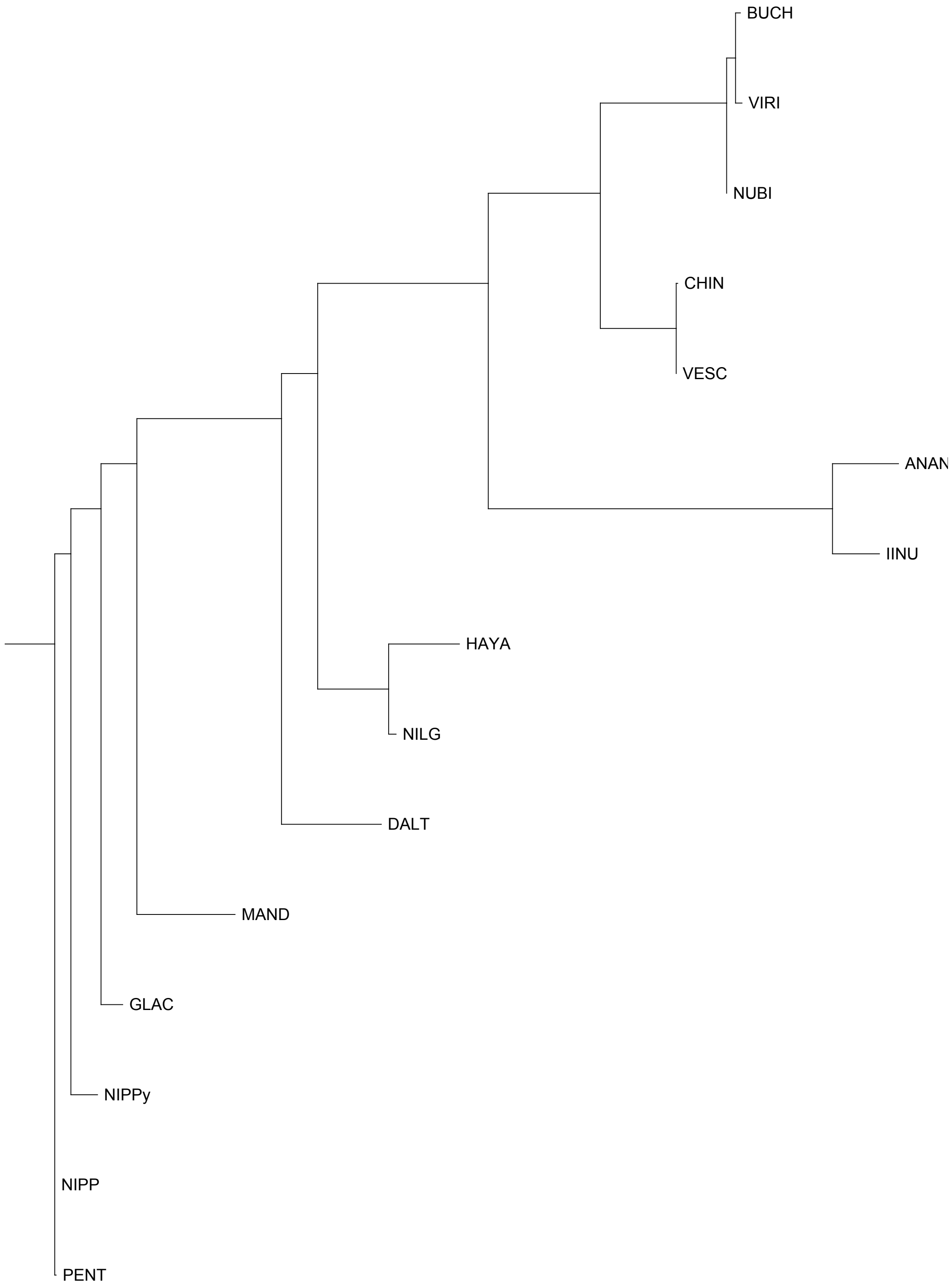


FAN_r2.3ch4X2b

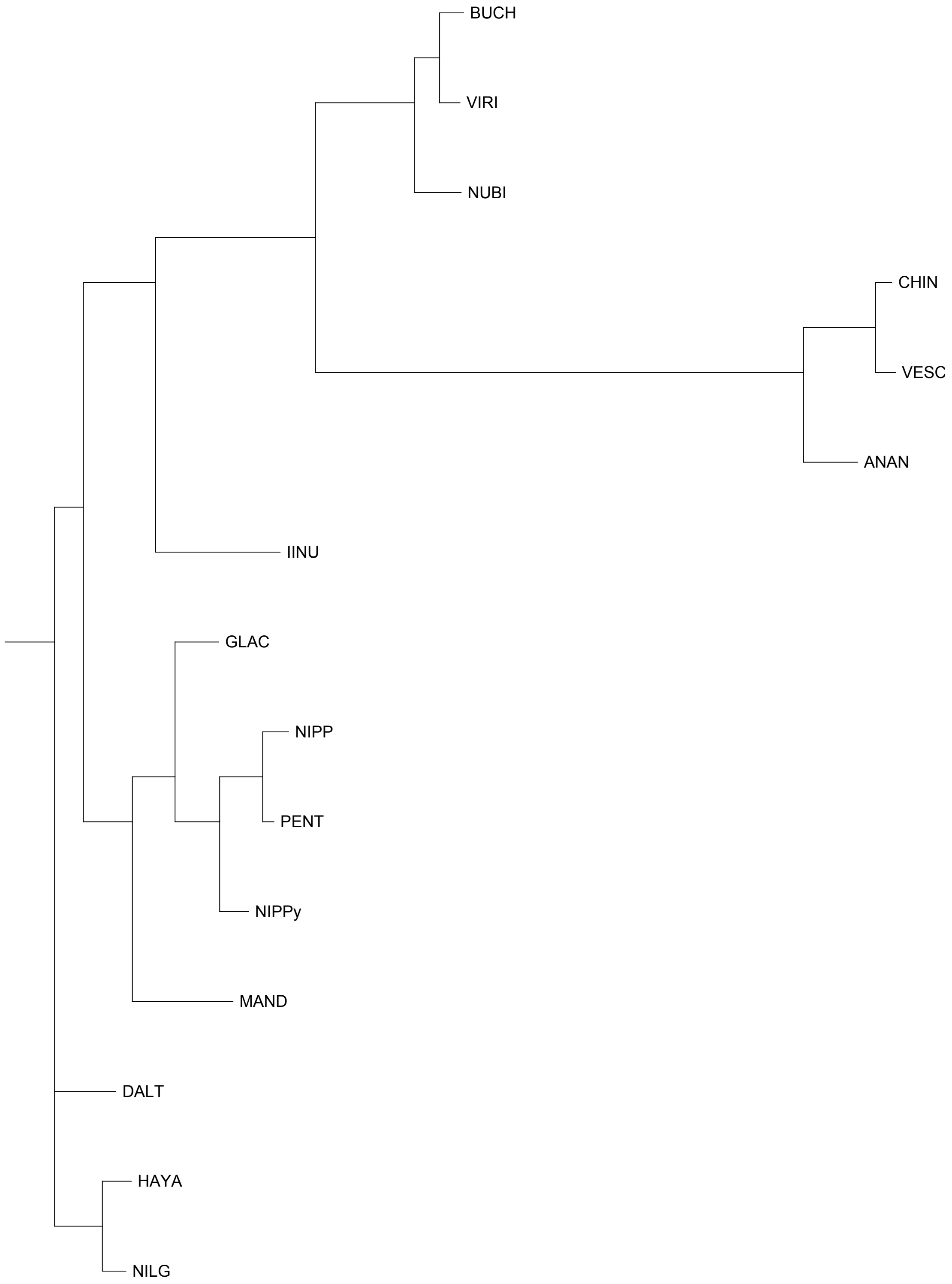


FAN_r2.3ch5Ava

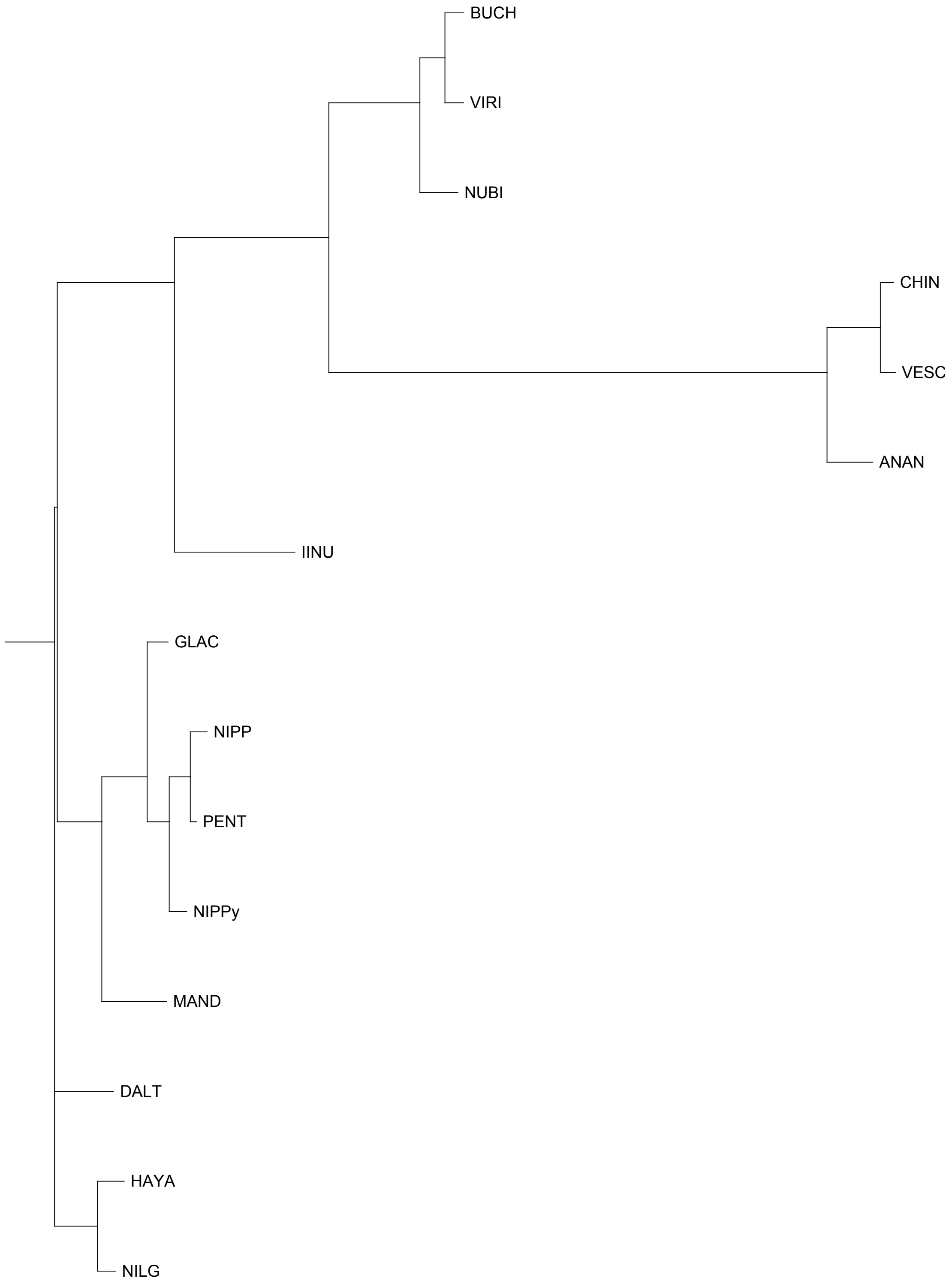


FAN_r2.3ch5Avb

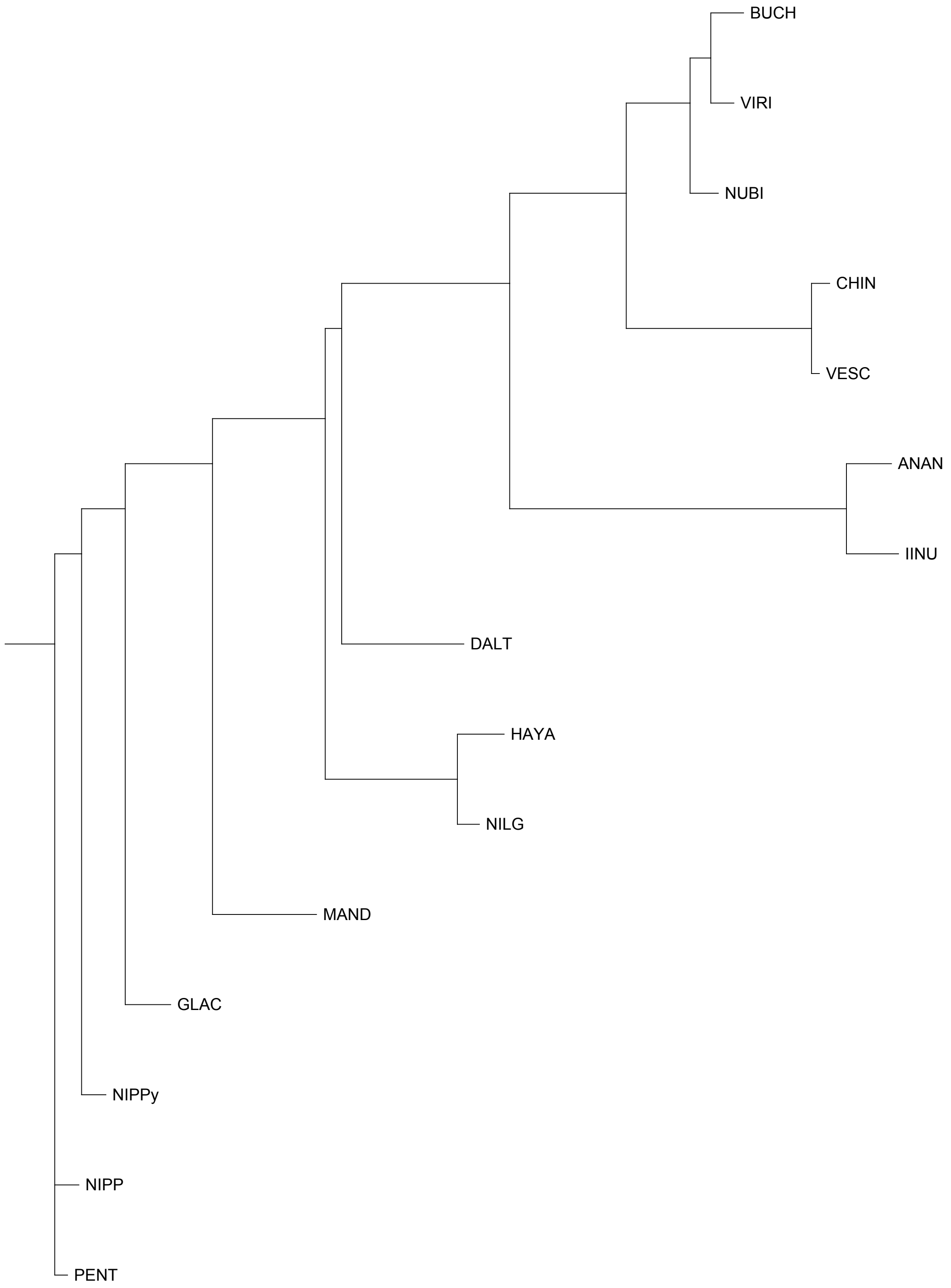


FAN_r2.3ch5Bia

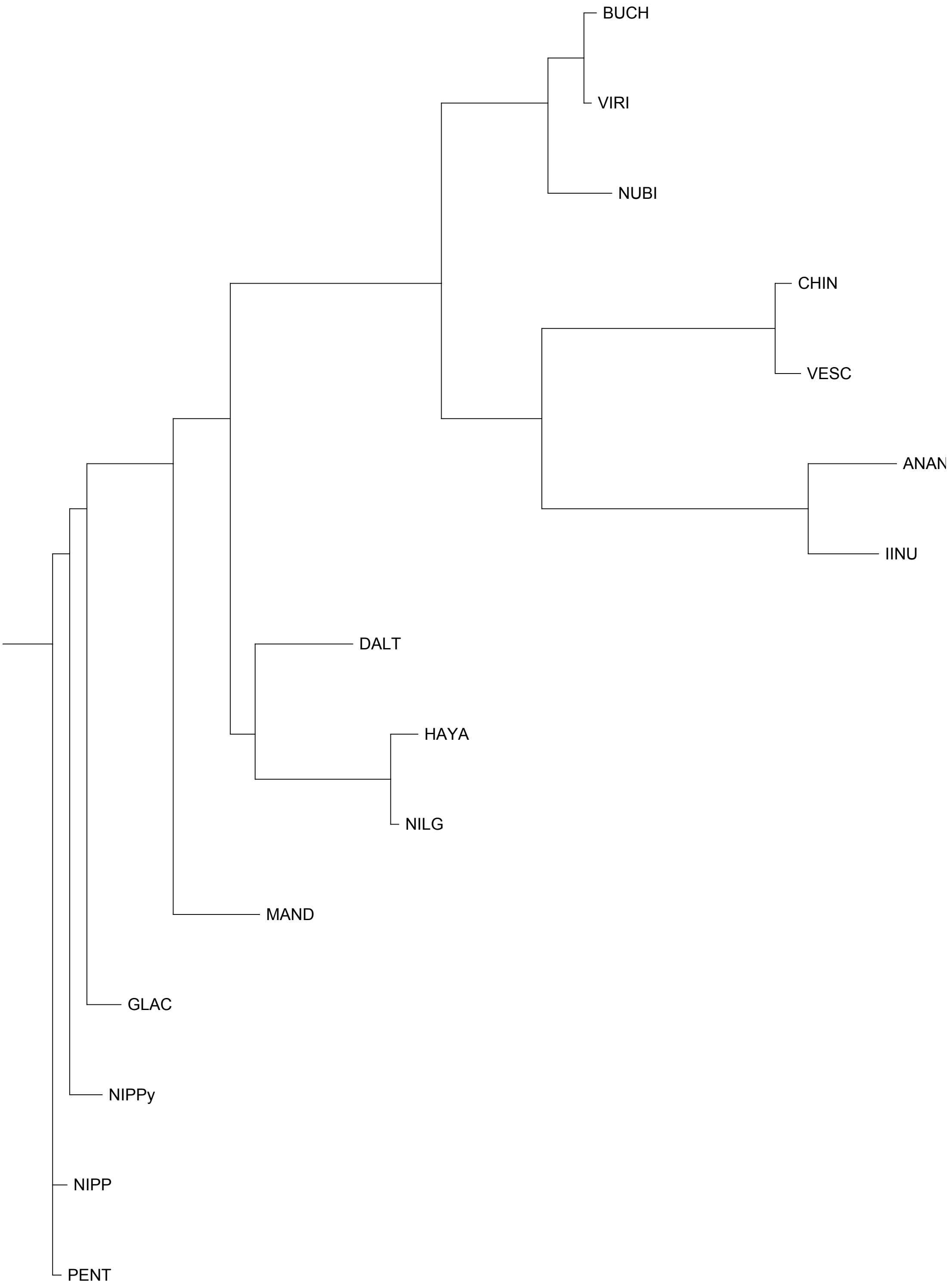


FAN_r2.3ch5Bib

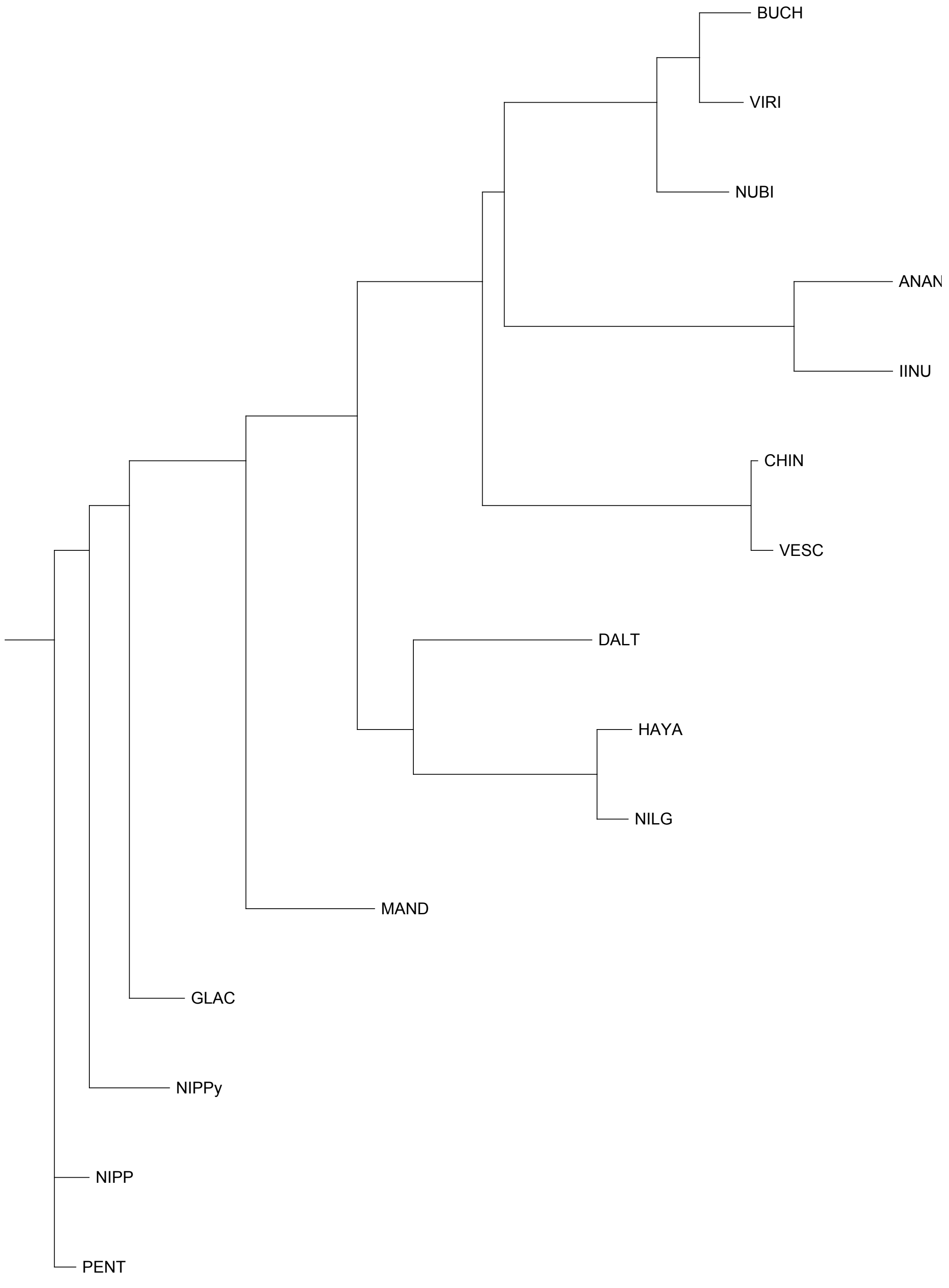


FAN_r2.3ch5X1a

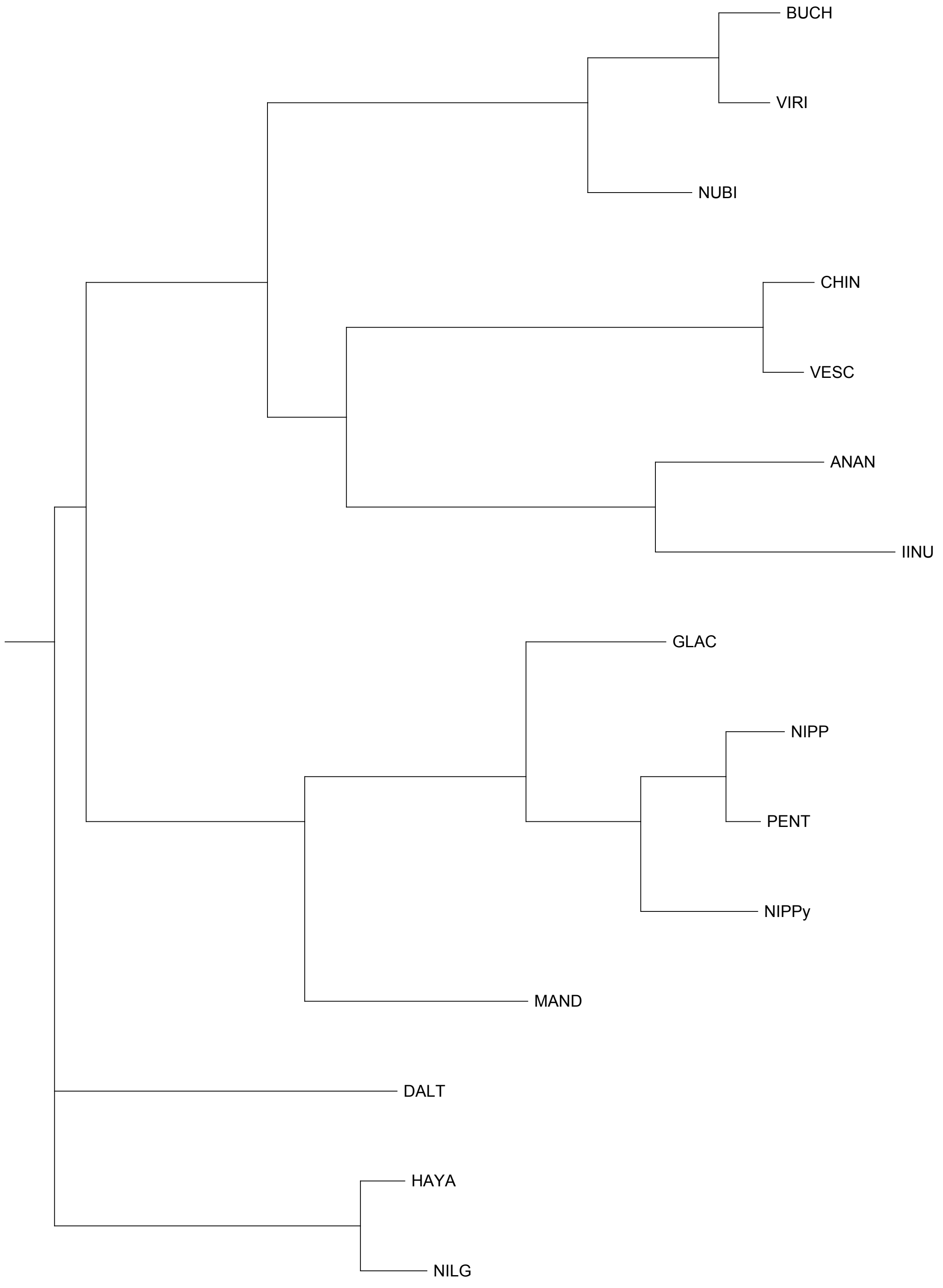


FAN_r2.3ch5X1b

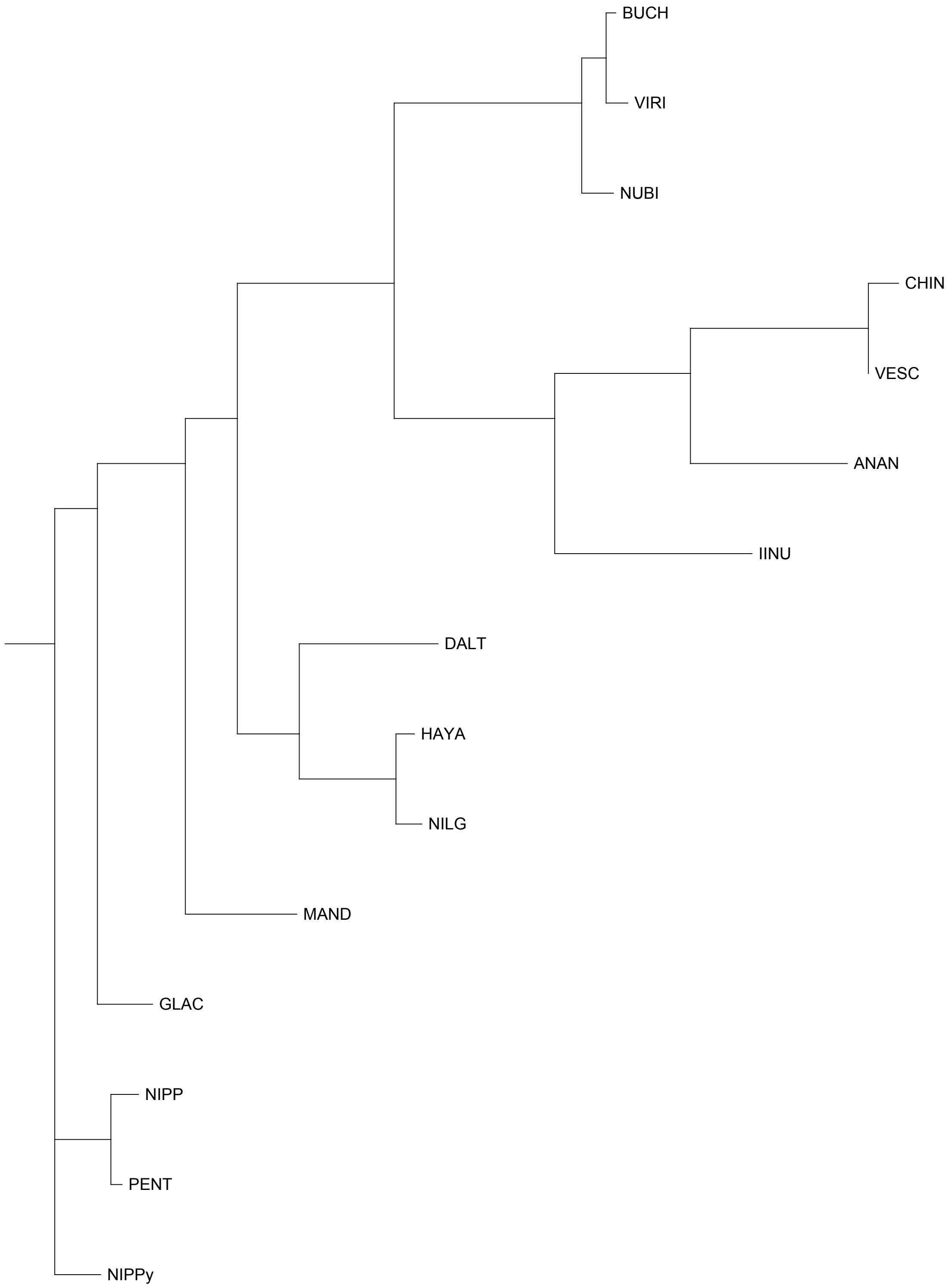


FAN_r2.3ch5X2a

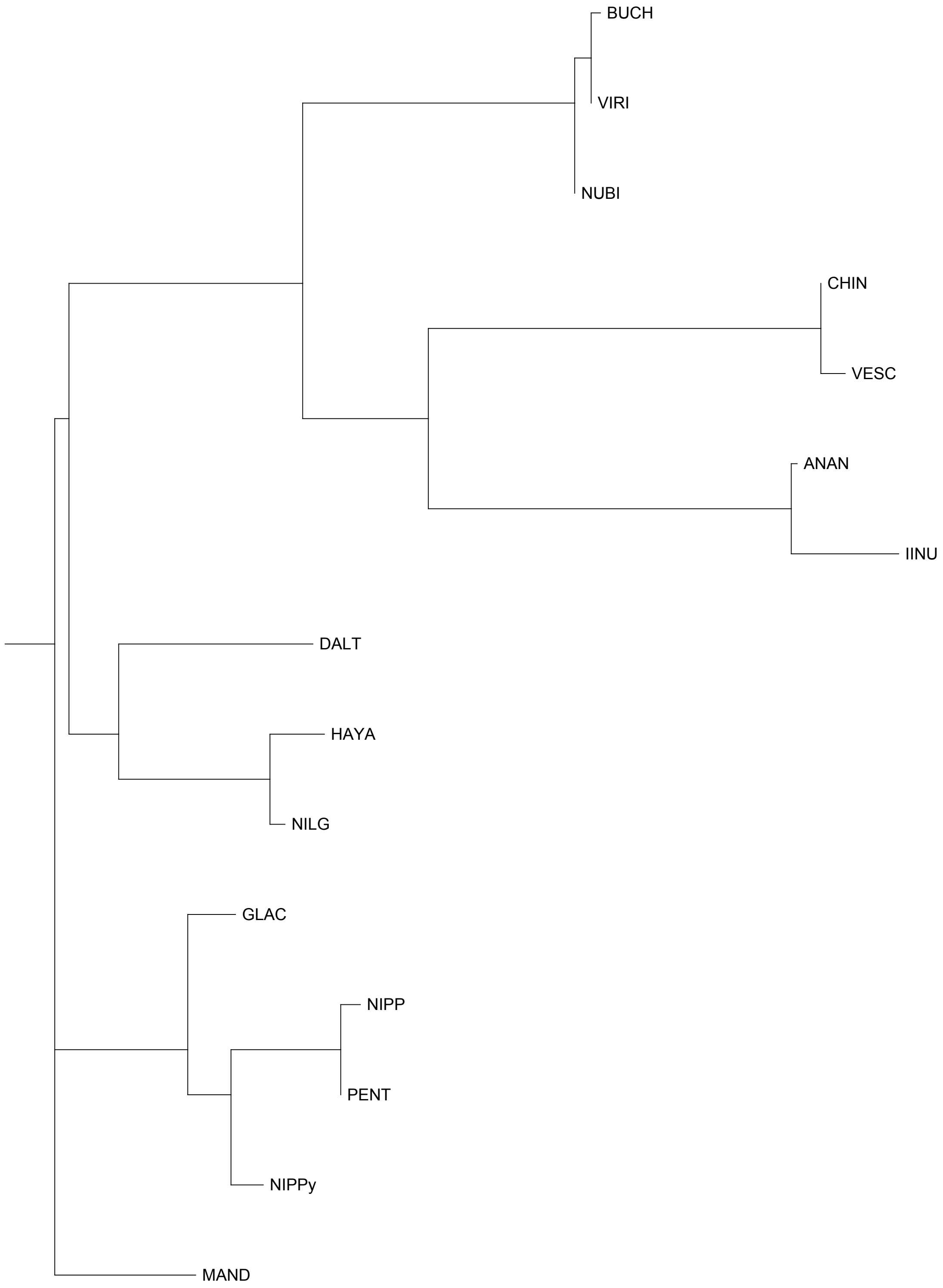


FAN_r2.3ch5X2b

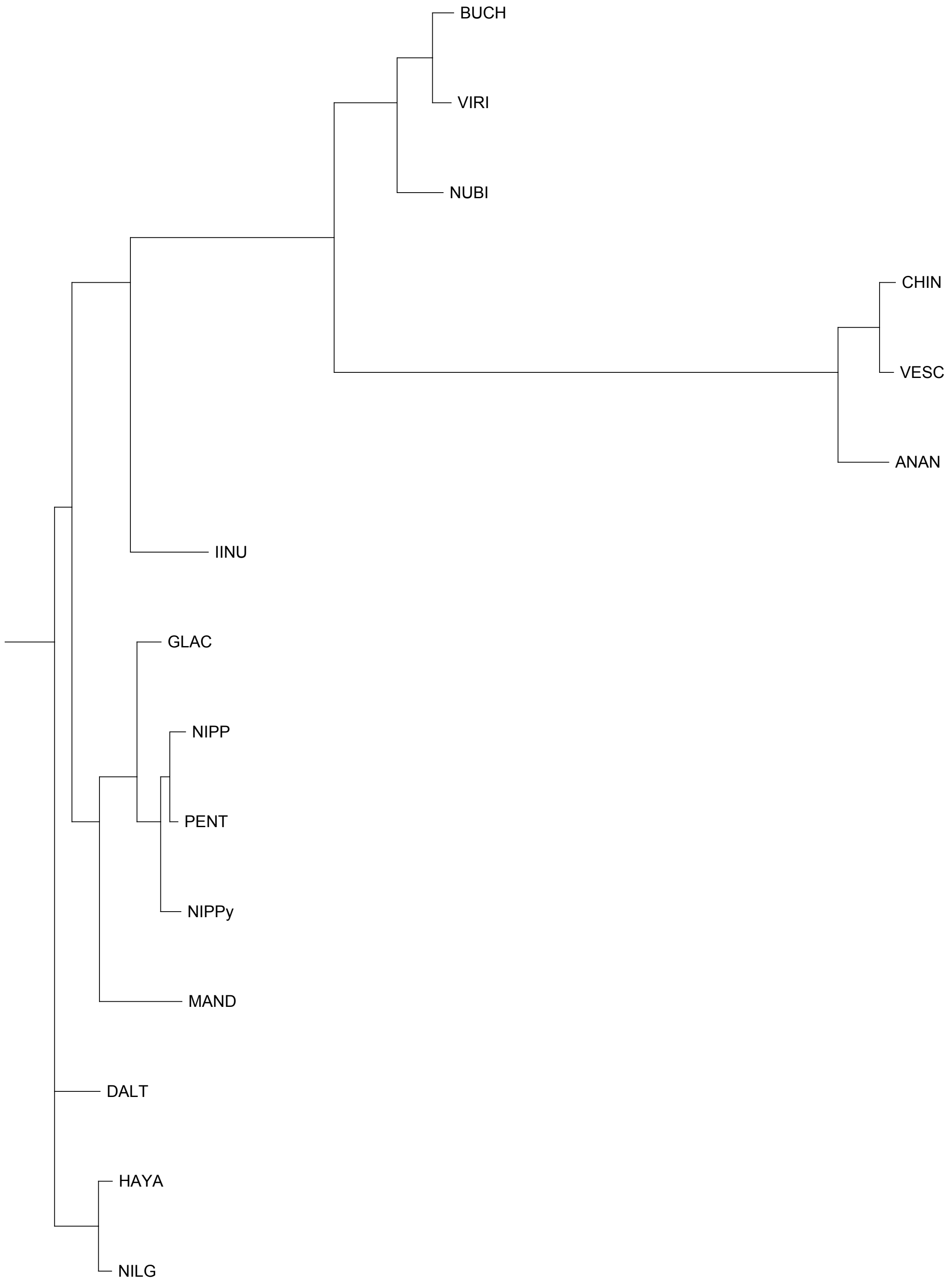


FAN_r2.3ch6Av1a

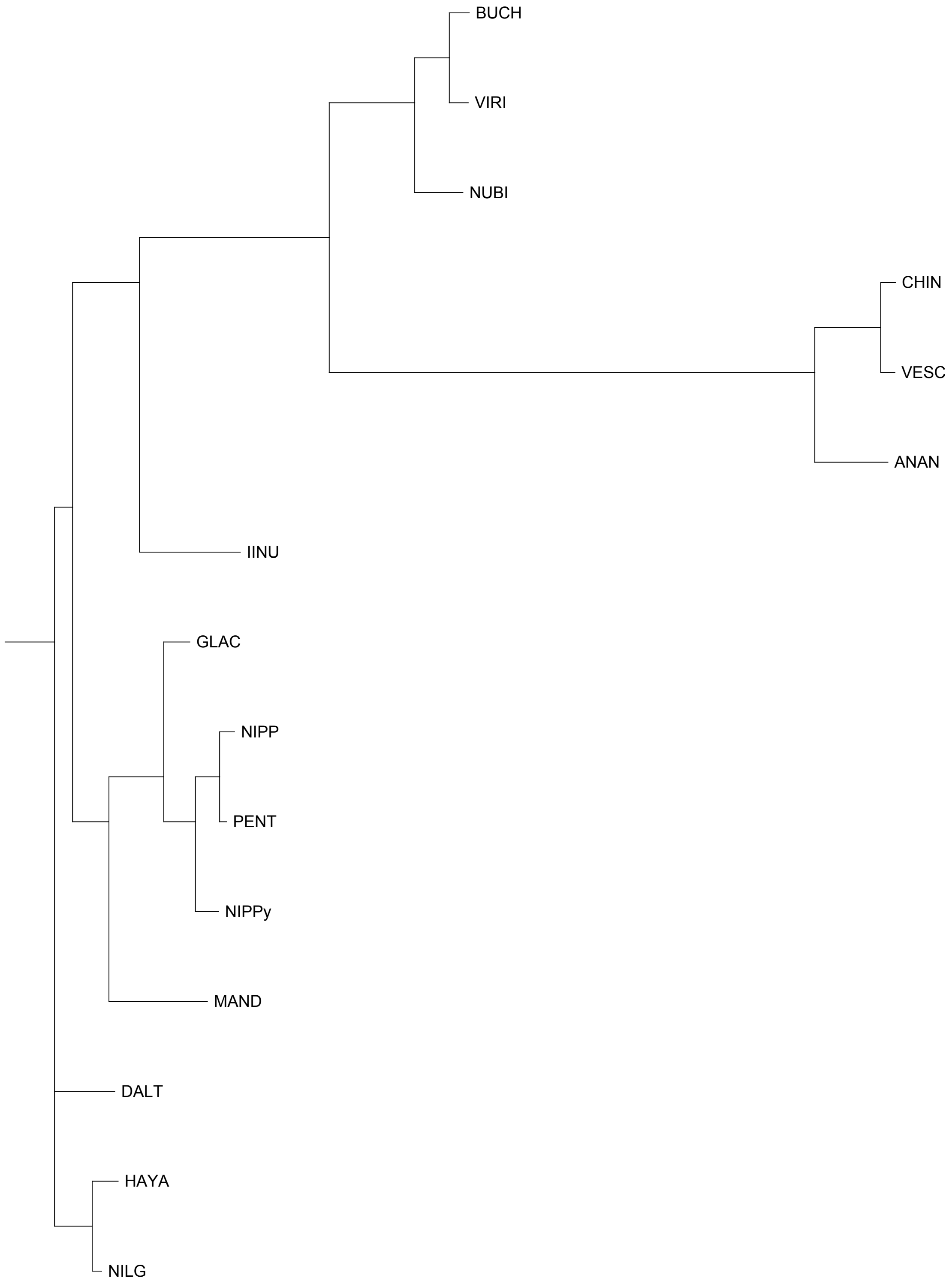


FAN_r2.3ch6Av1b

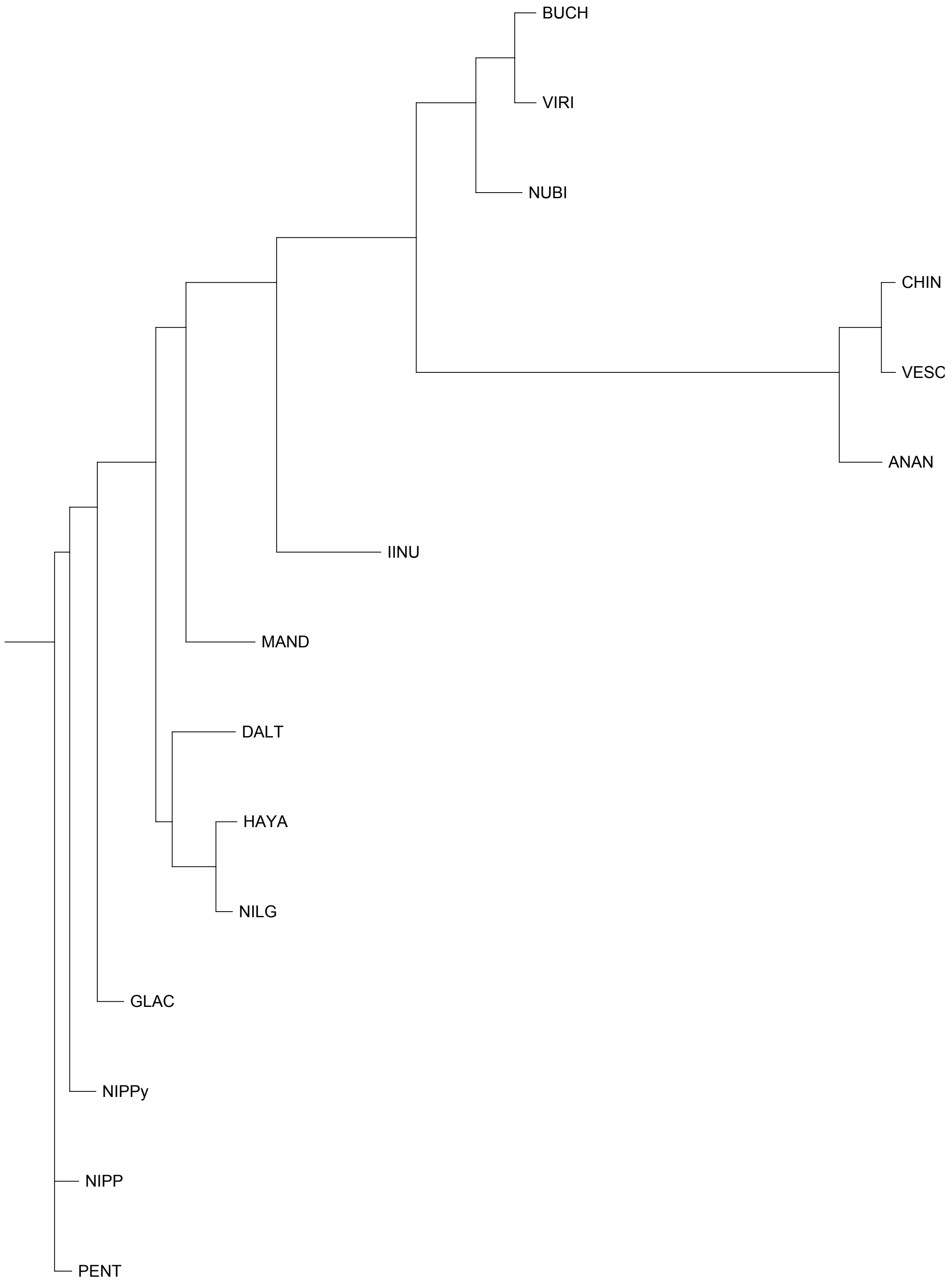


FAN_r2.3ch6Av2a

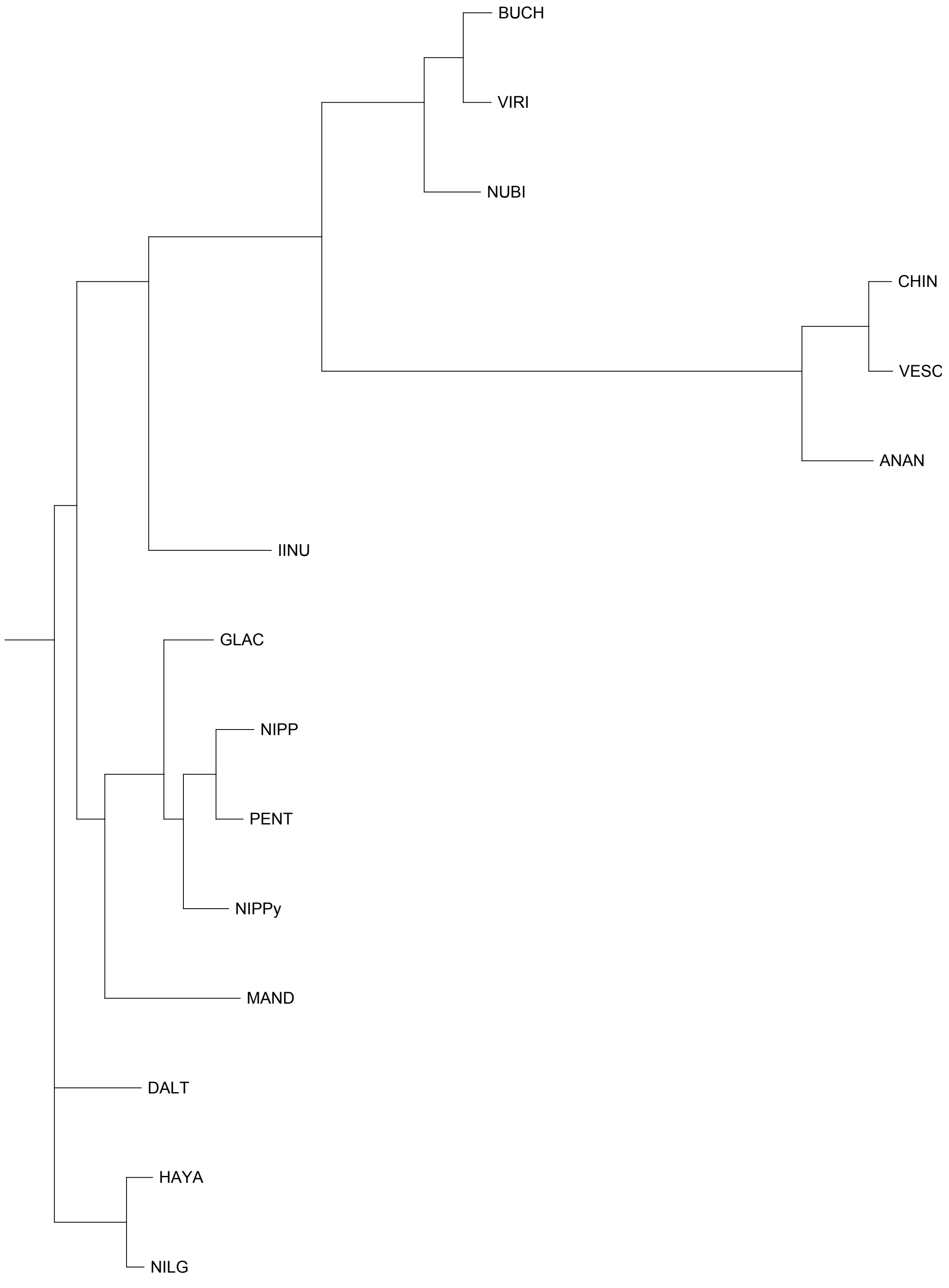


FAN_r2.3ch6Av2b

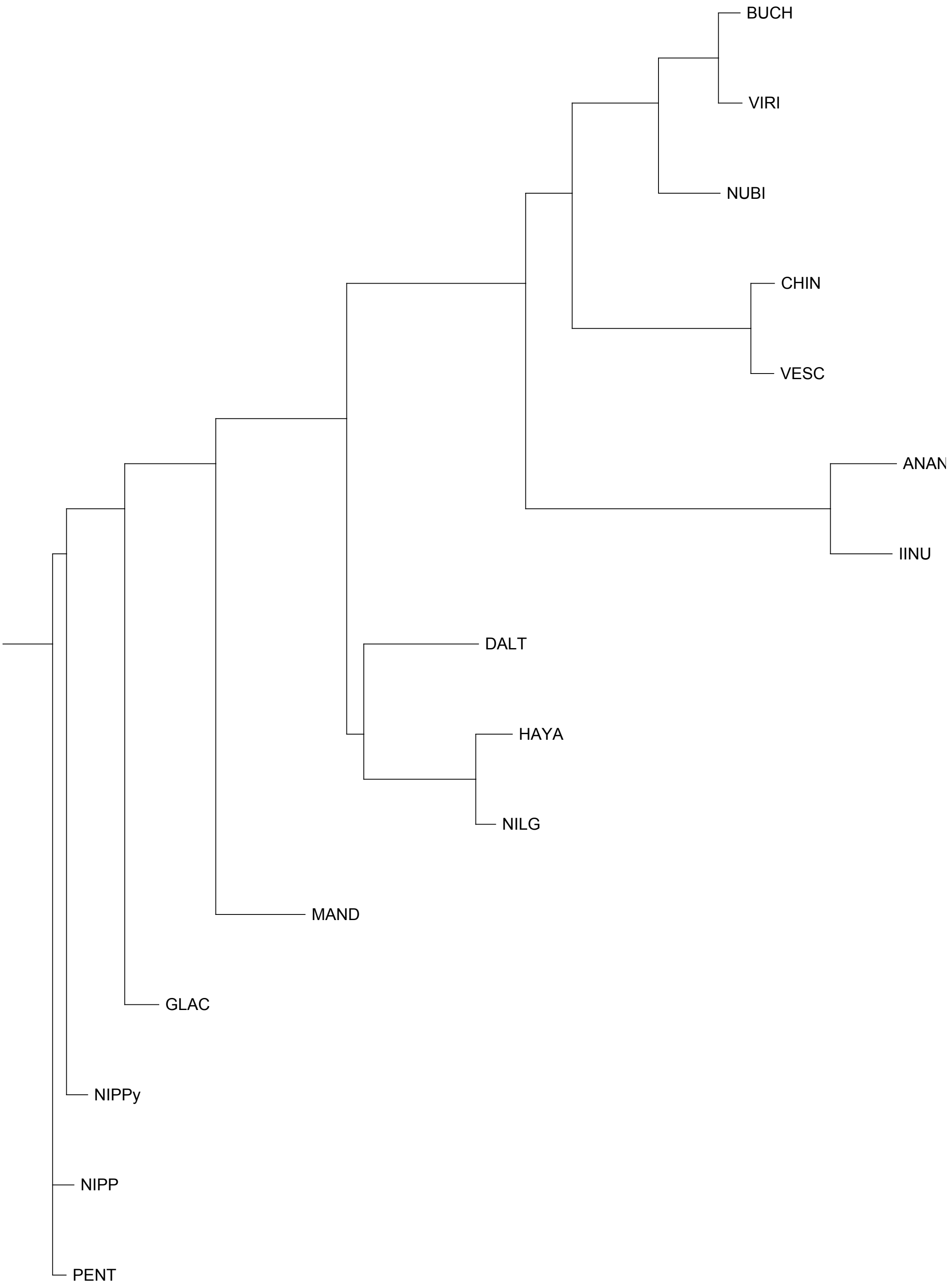


FAN_r2.3ch6Bia

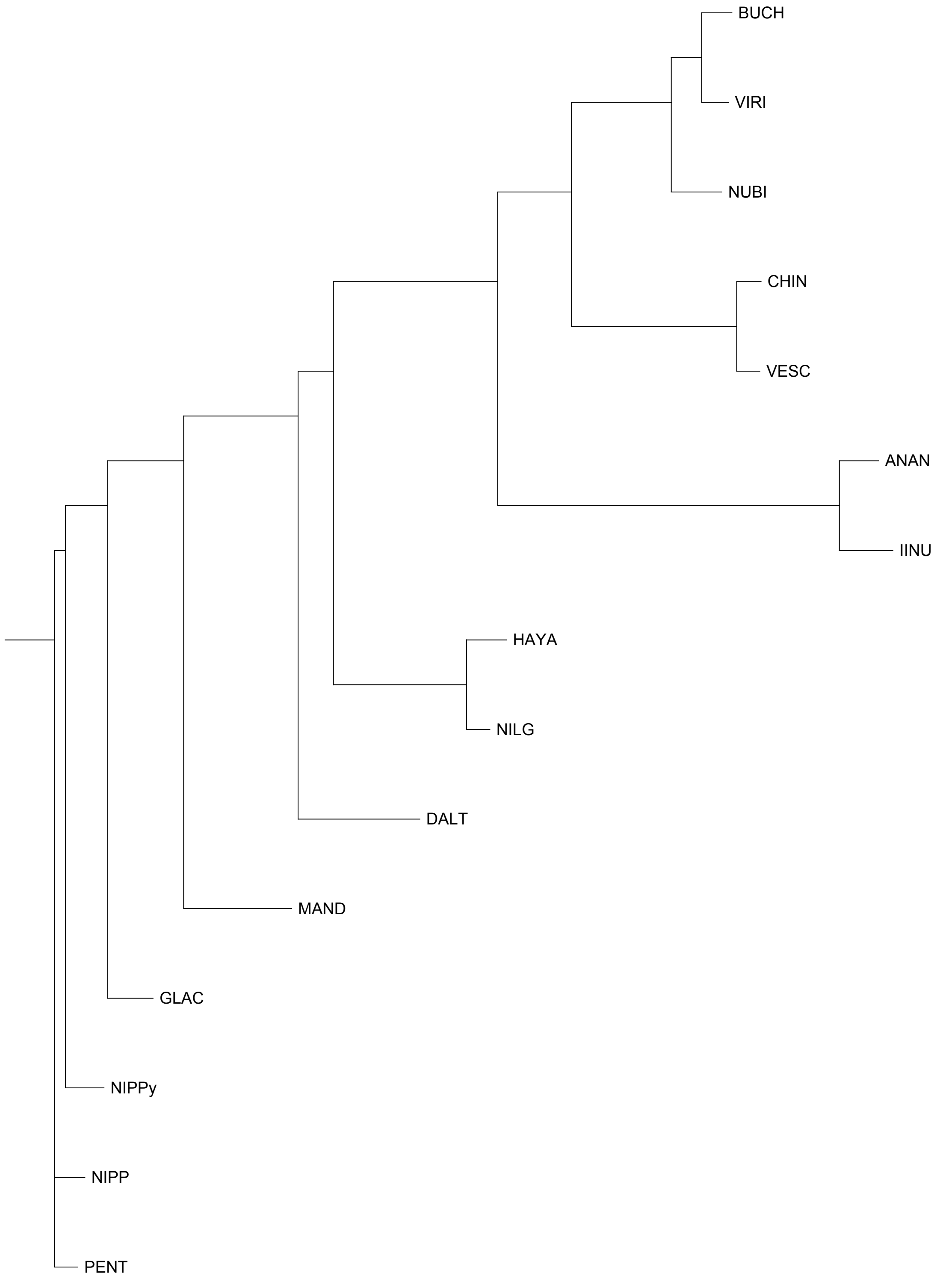


FAN_r2.3ch6Bib

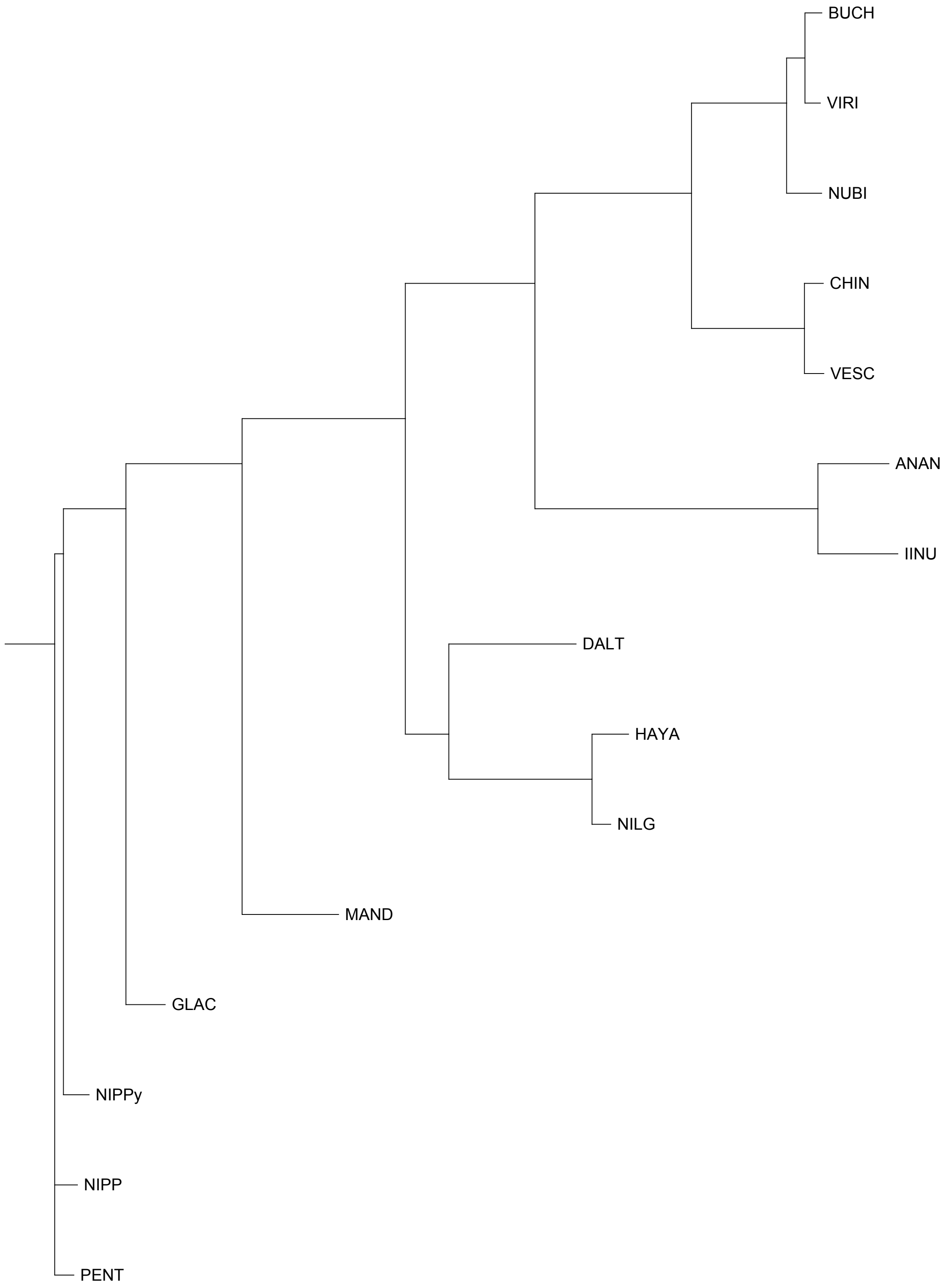


FAN_r2.3ch6X1a

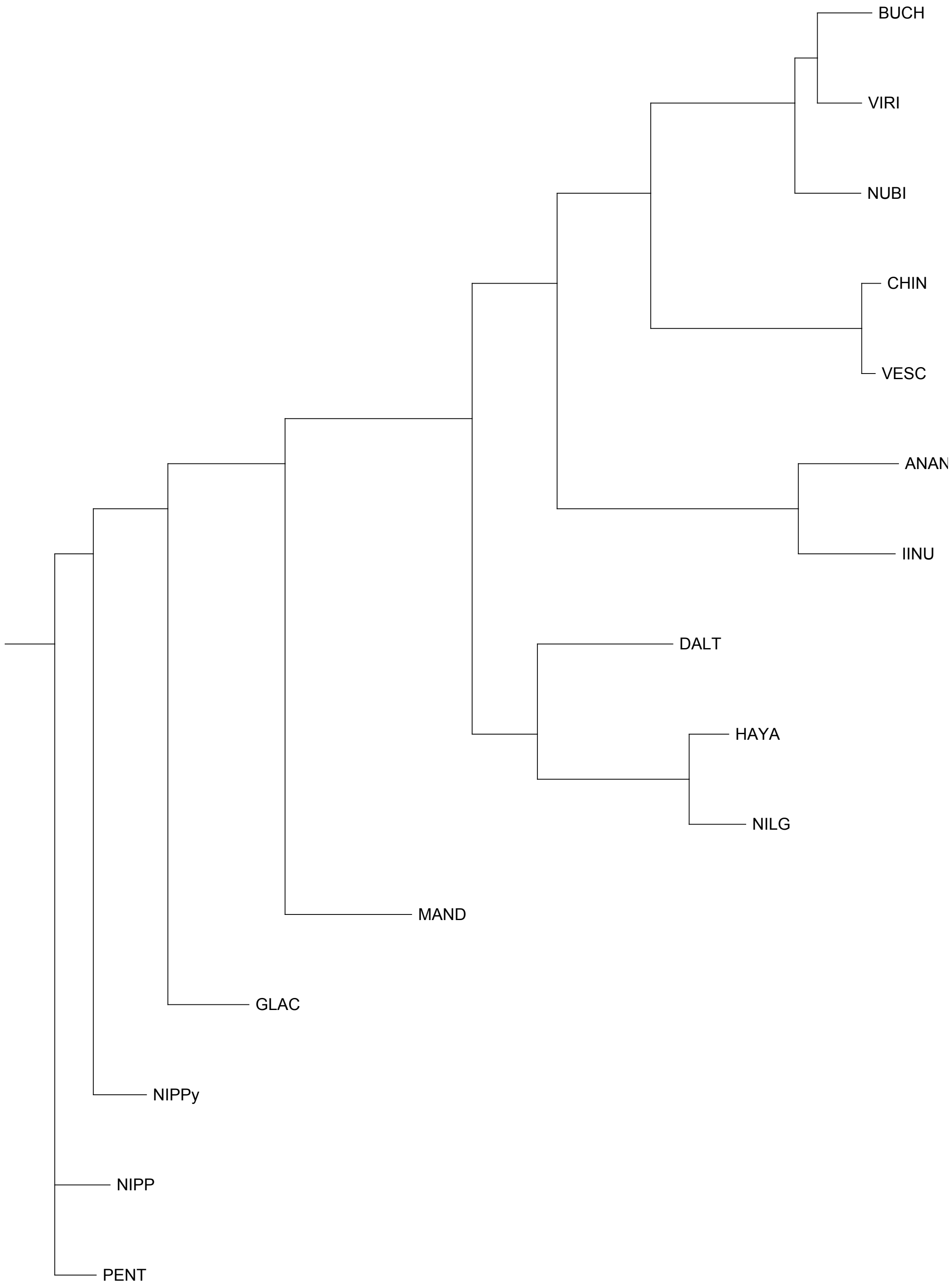


FAN_r2.3ch6X1b

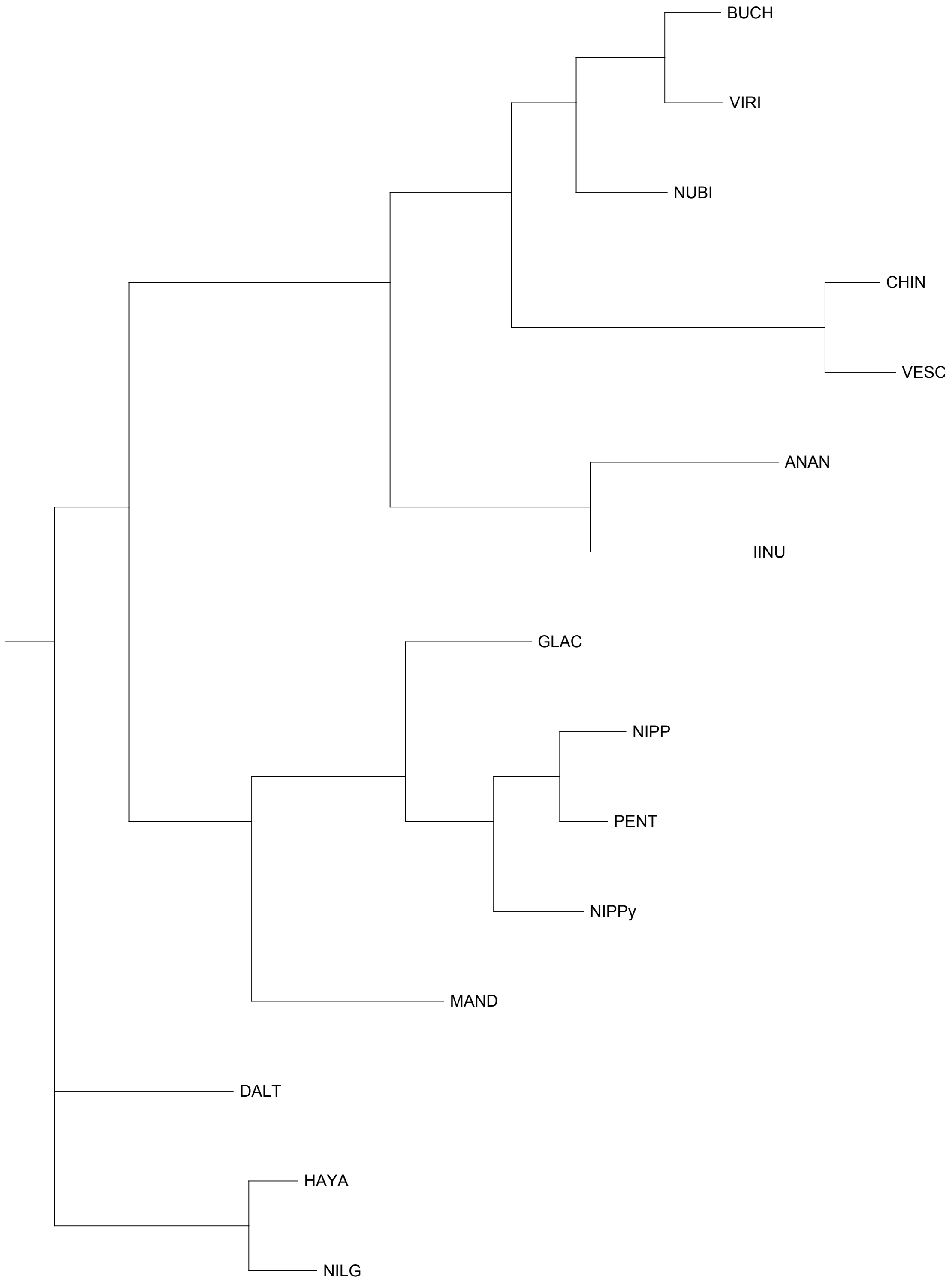


FAN_r2.3ch6X2a

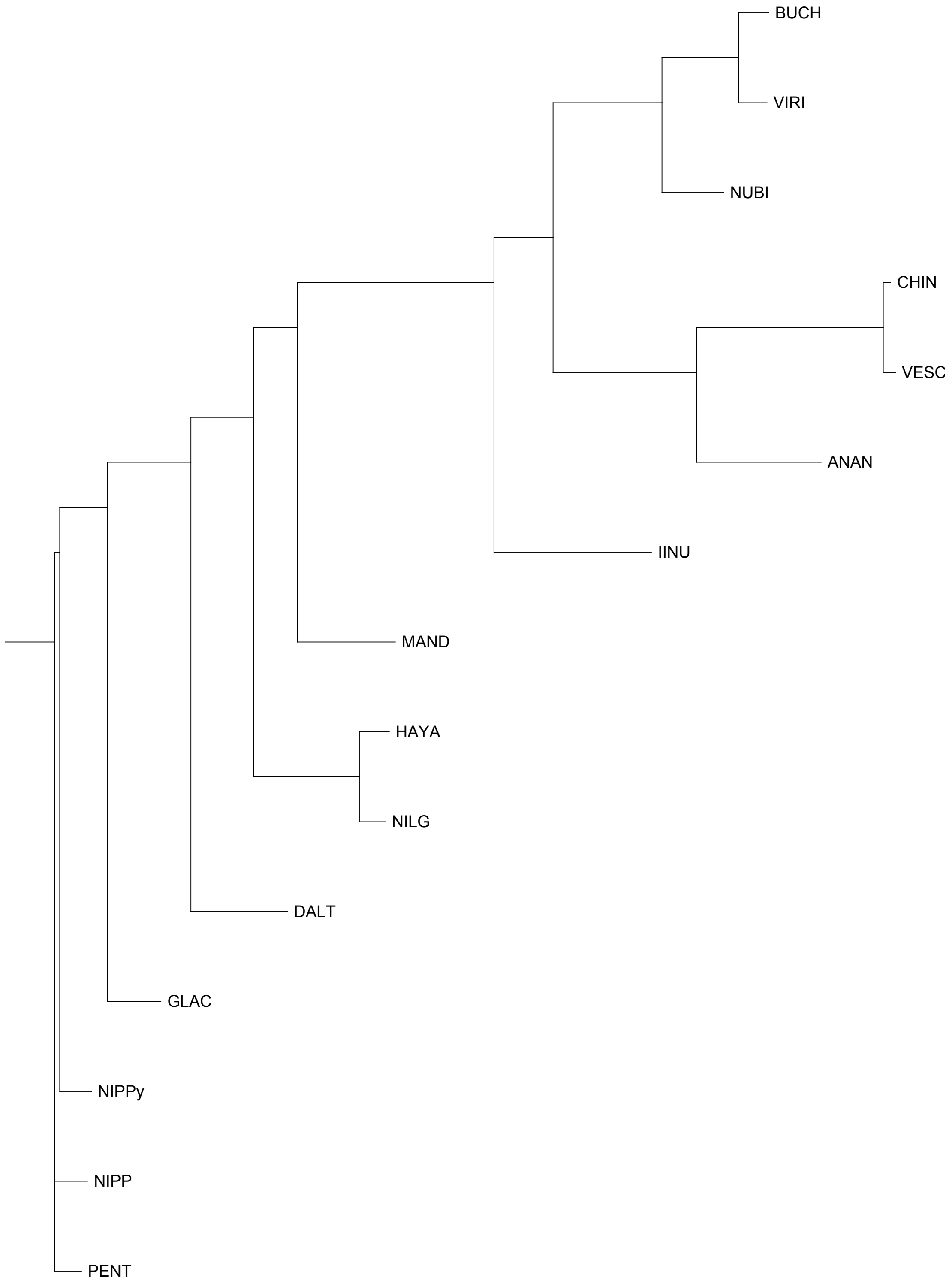


FAN_r2.3ch6X2b

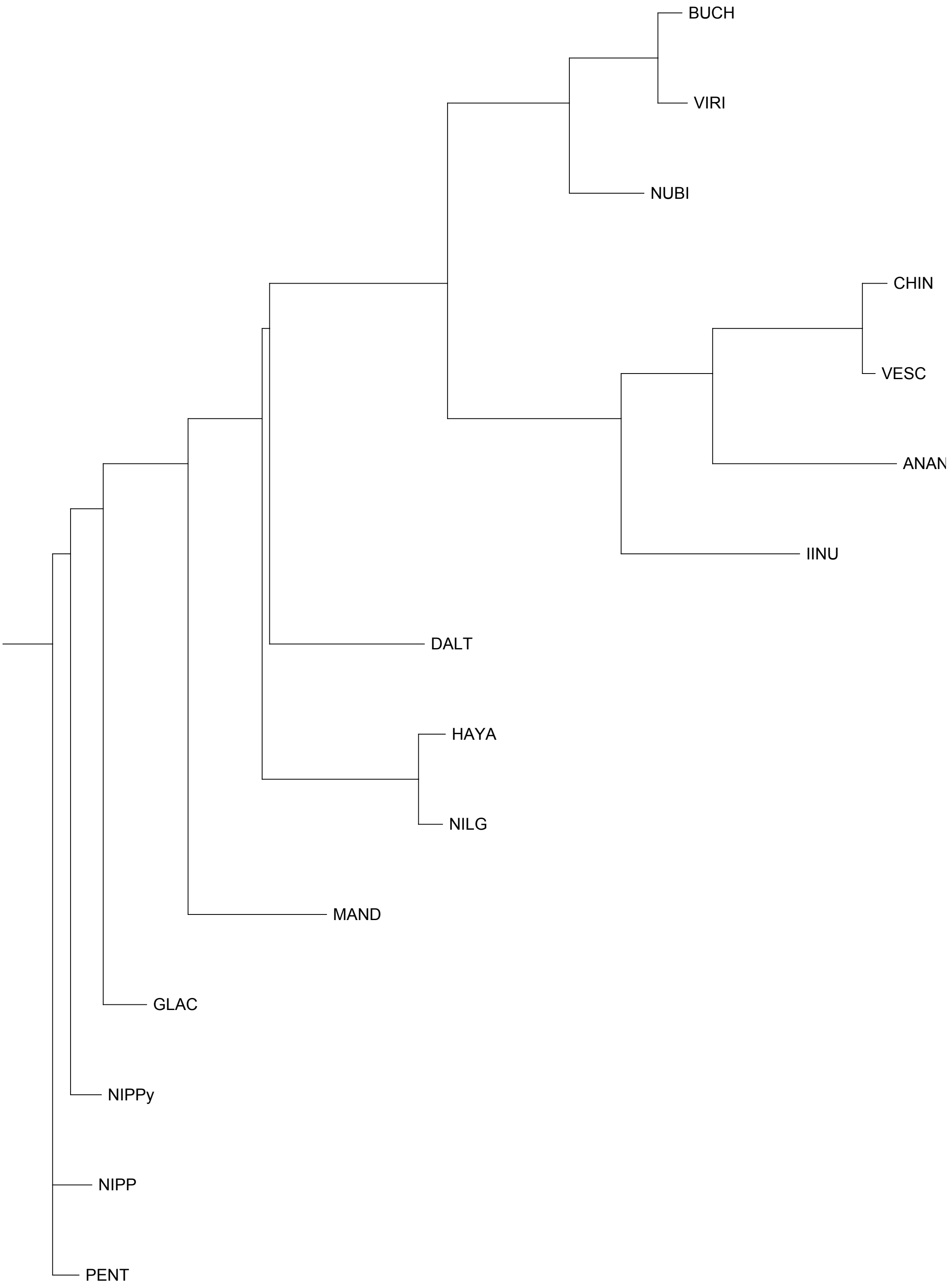


FAN_r2.3ch6X3a

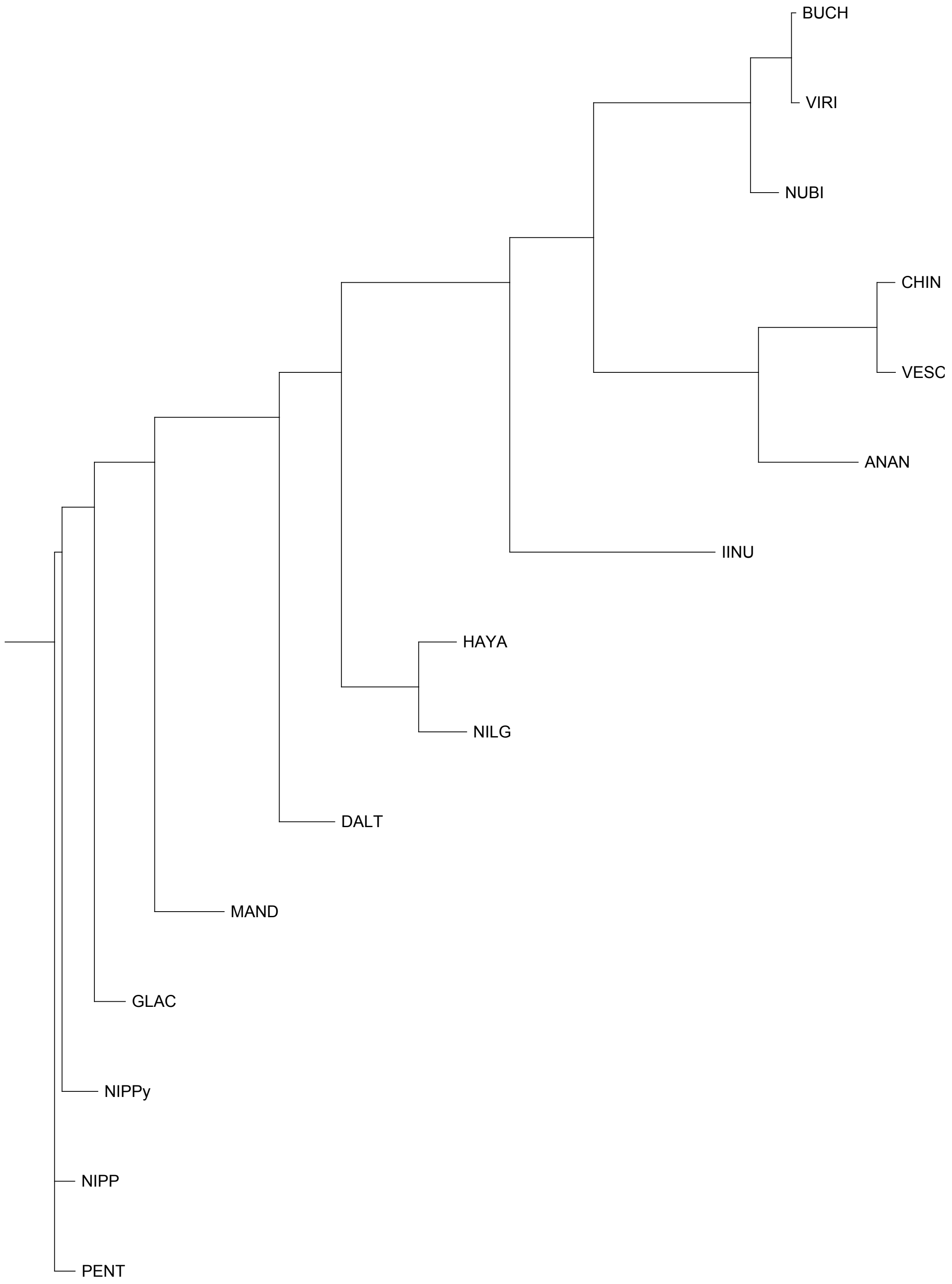


FAN_r2.3ch6X3b

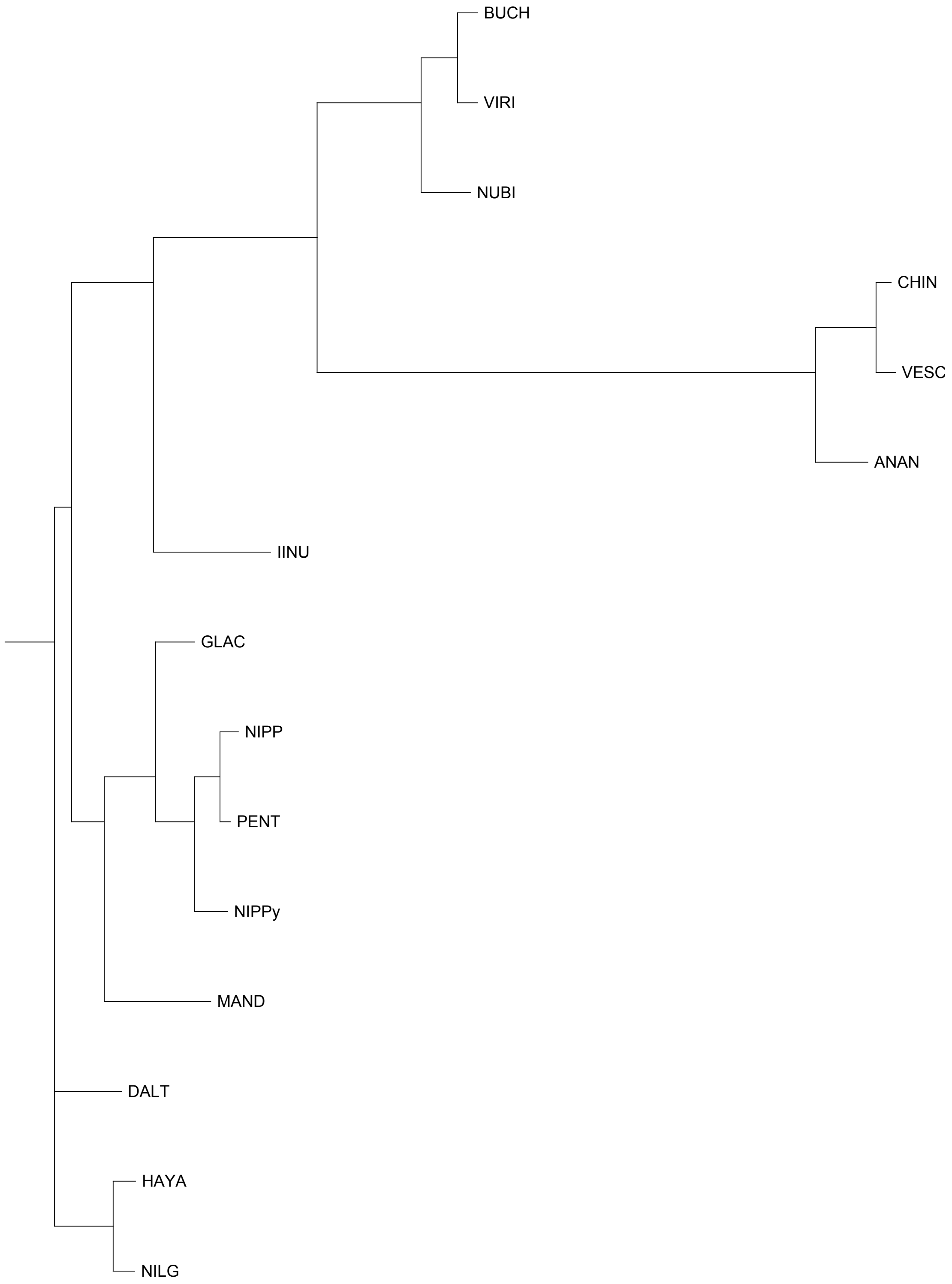


FAN_r2.3ch7Ava

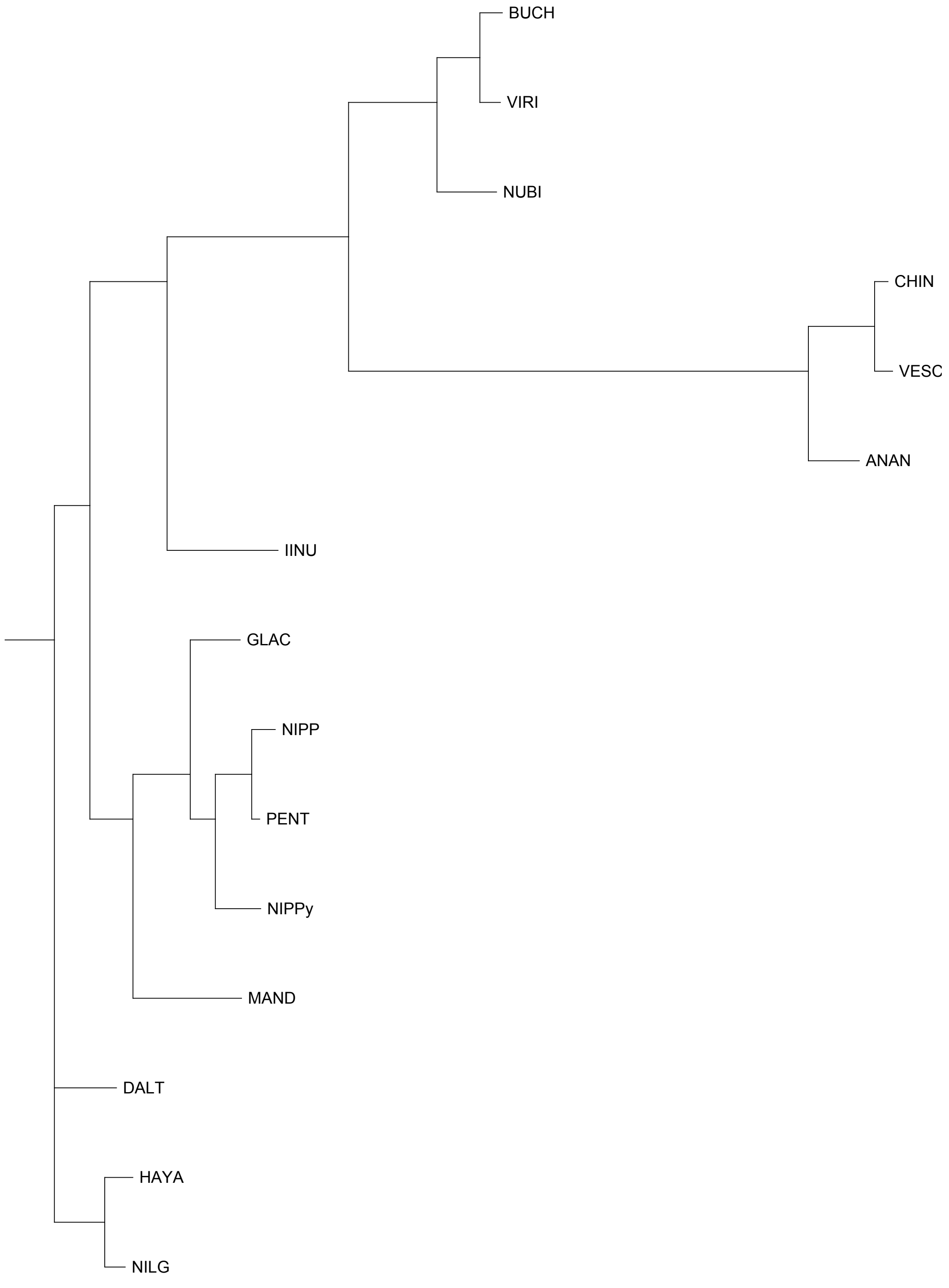


FAN_r2.3ch7Avb

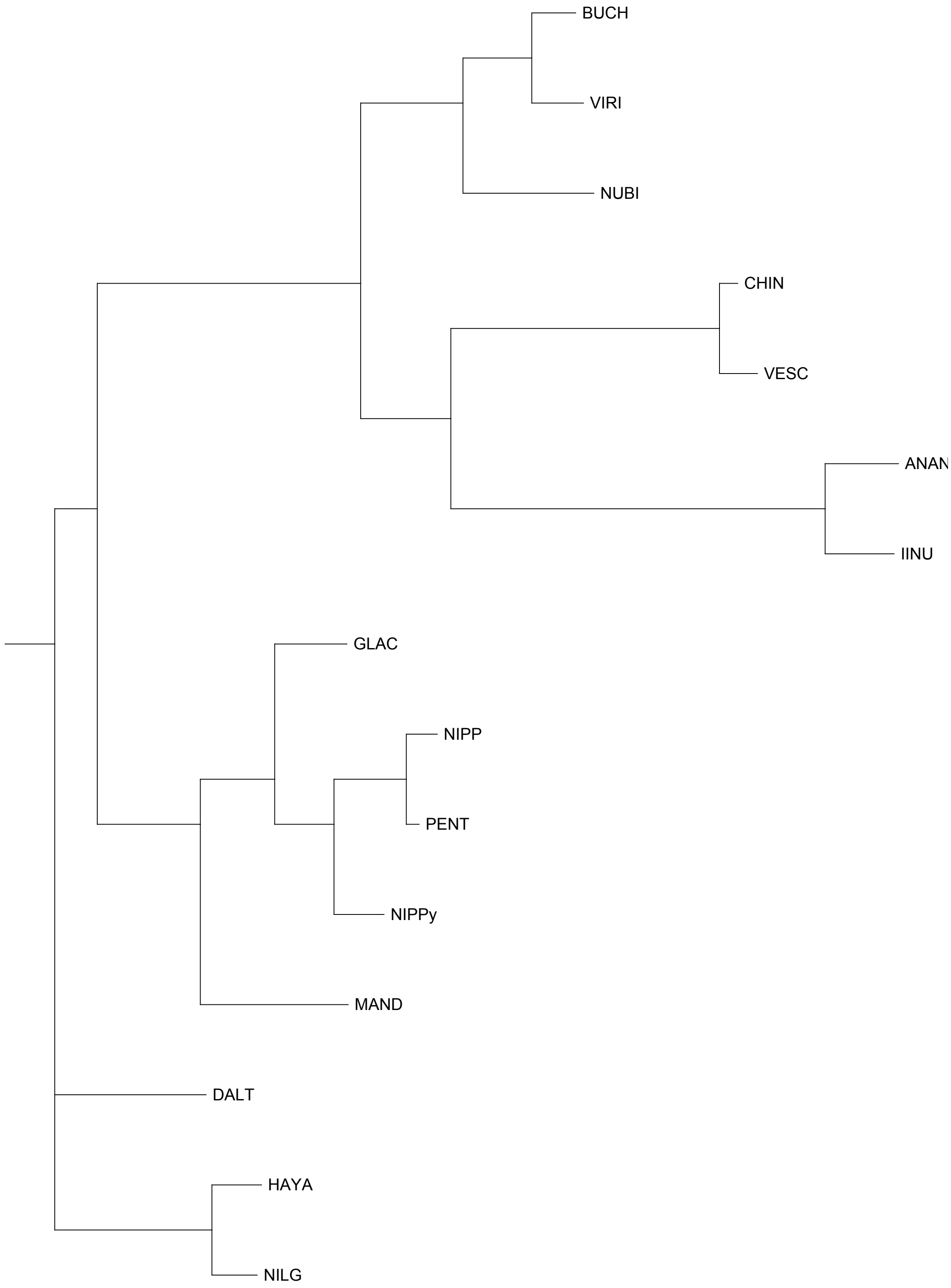


FAN_r2.3ch7Bia

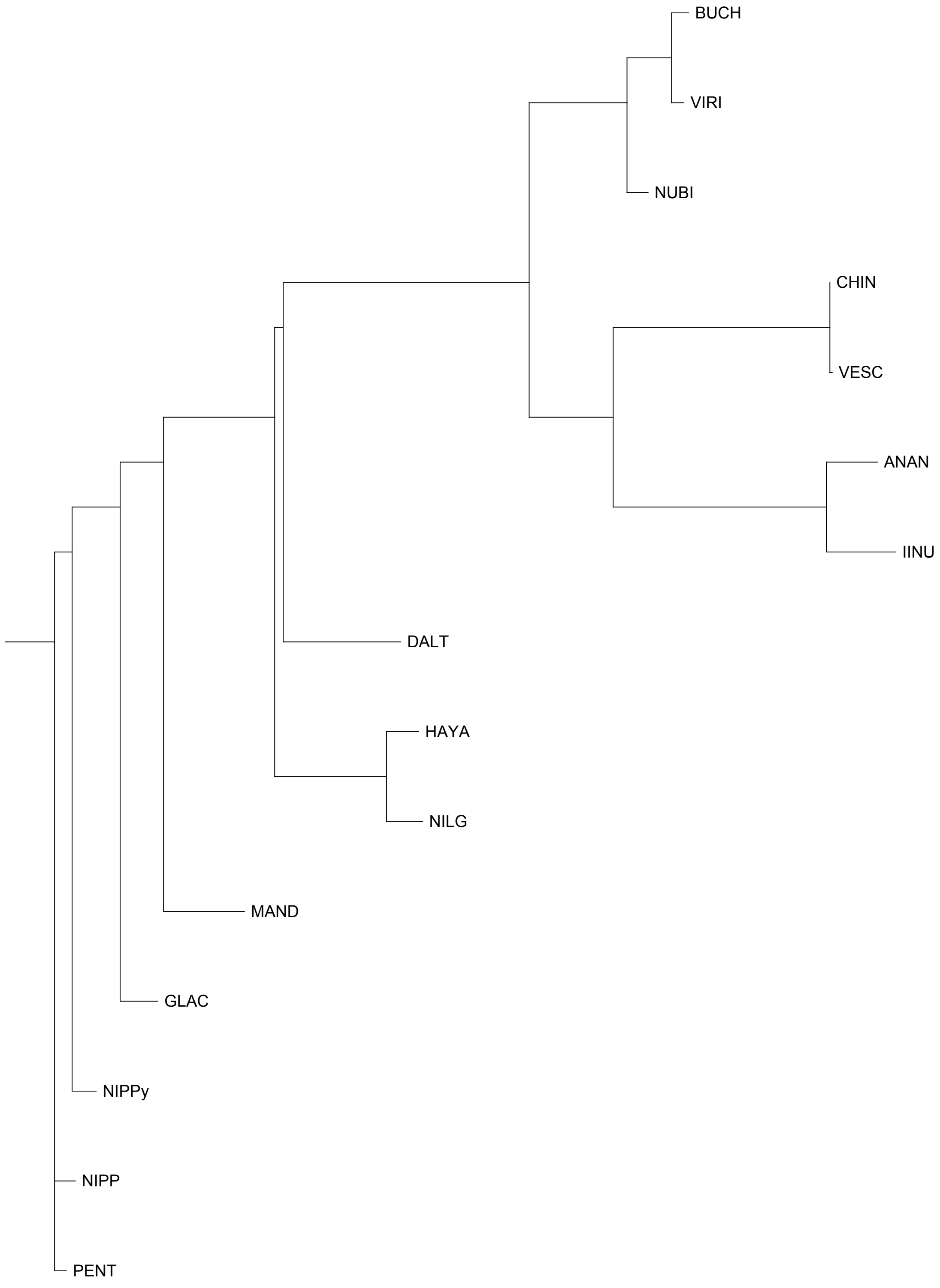


FAN_r2.3ch7Bib

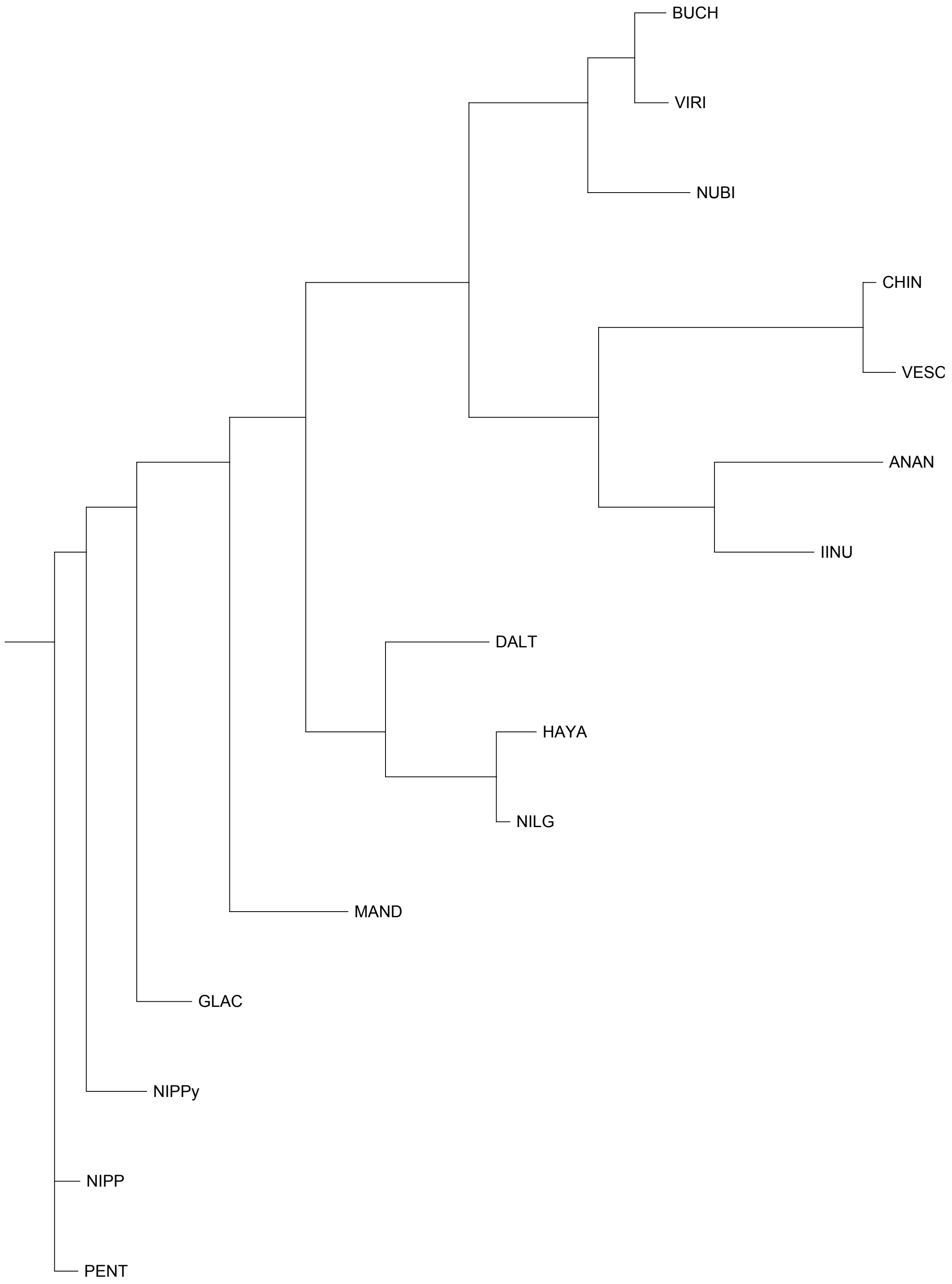


FAN_r2.3ch7X1a

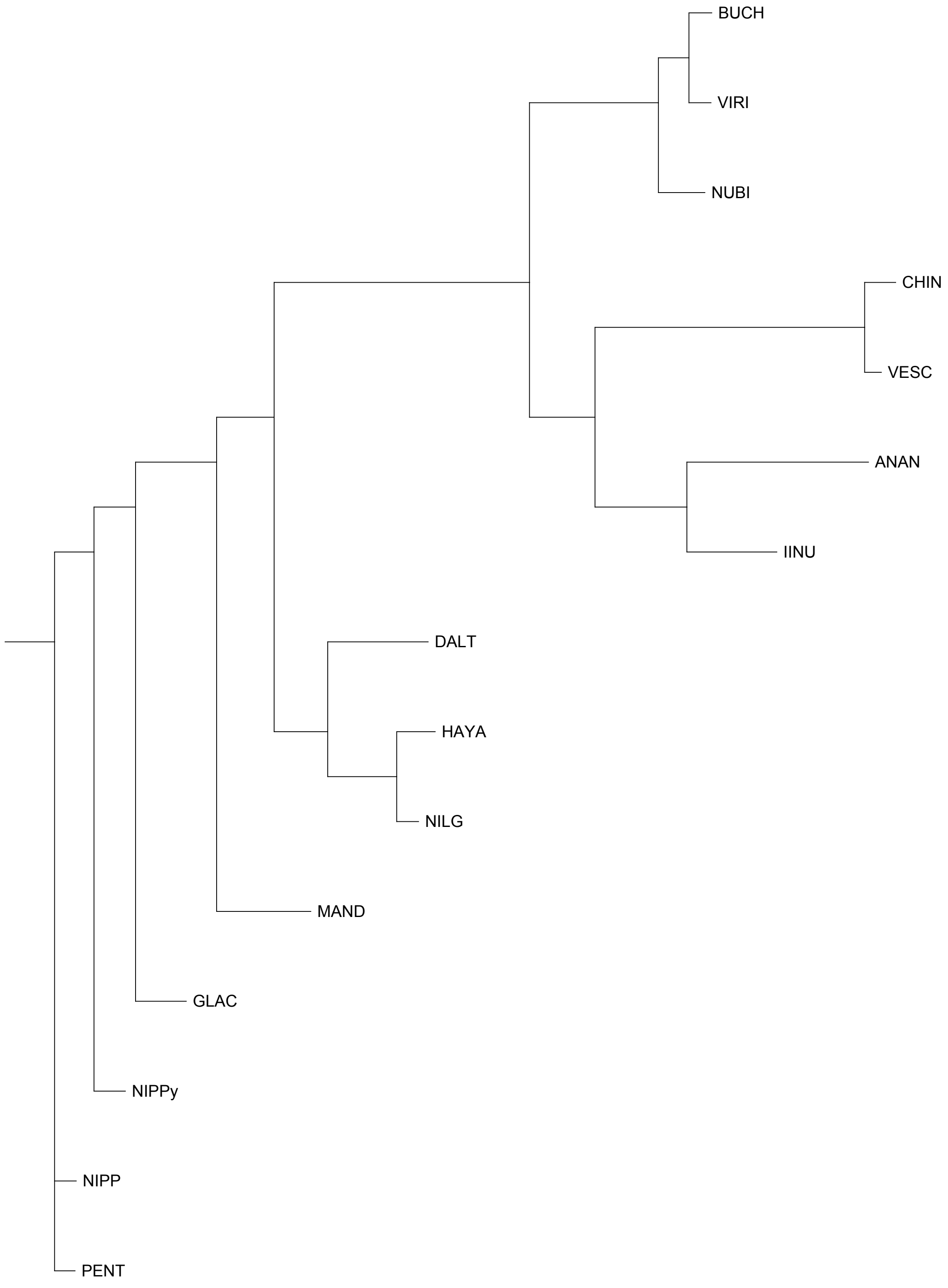


FAN_r2.3ch7X1b

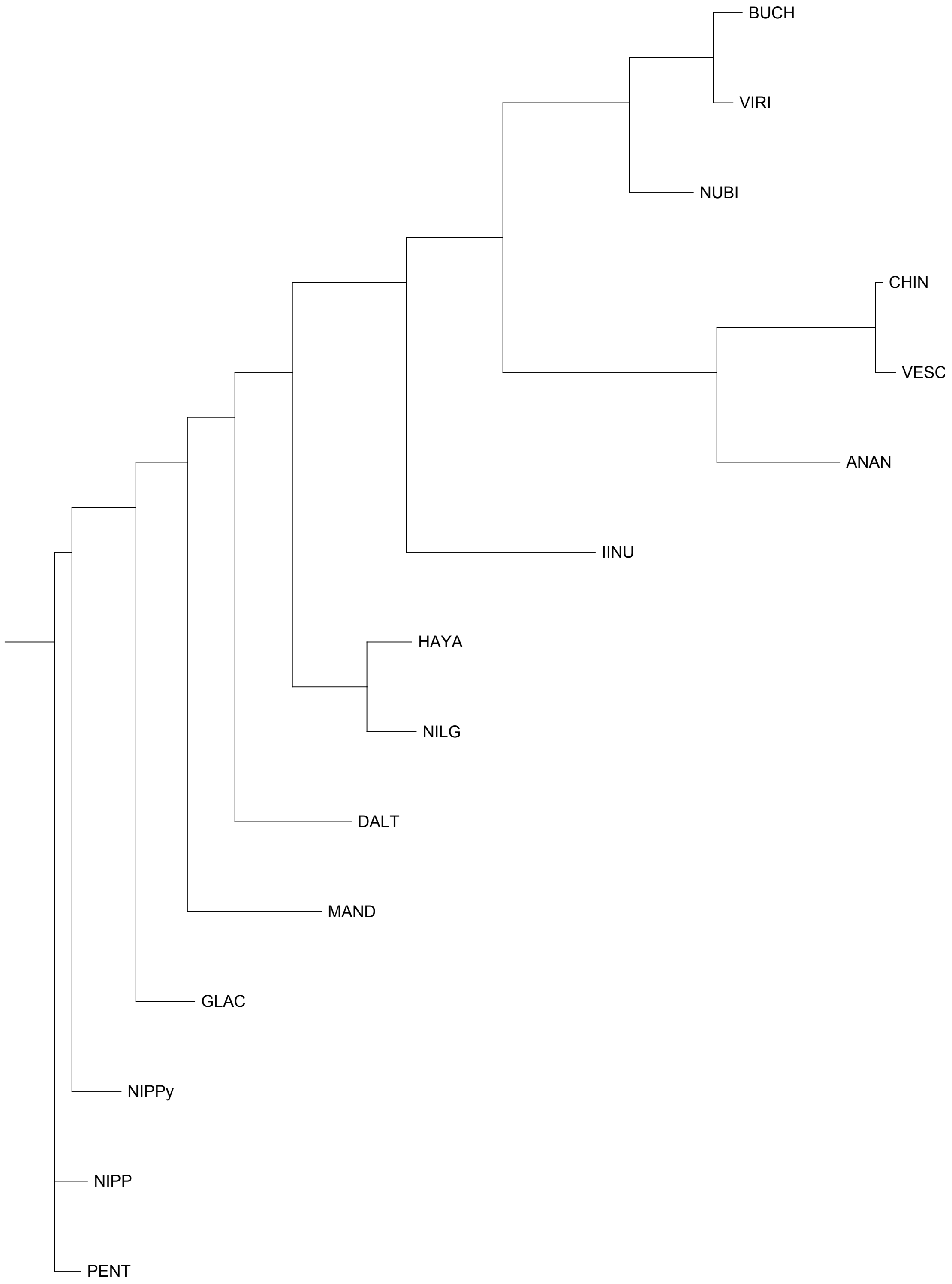


FAN_r2.3ch7X2a

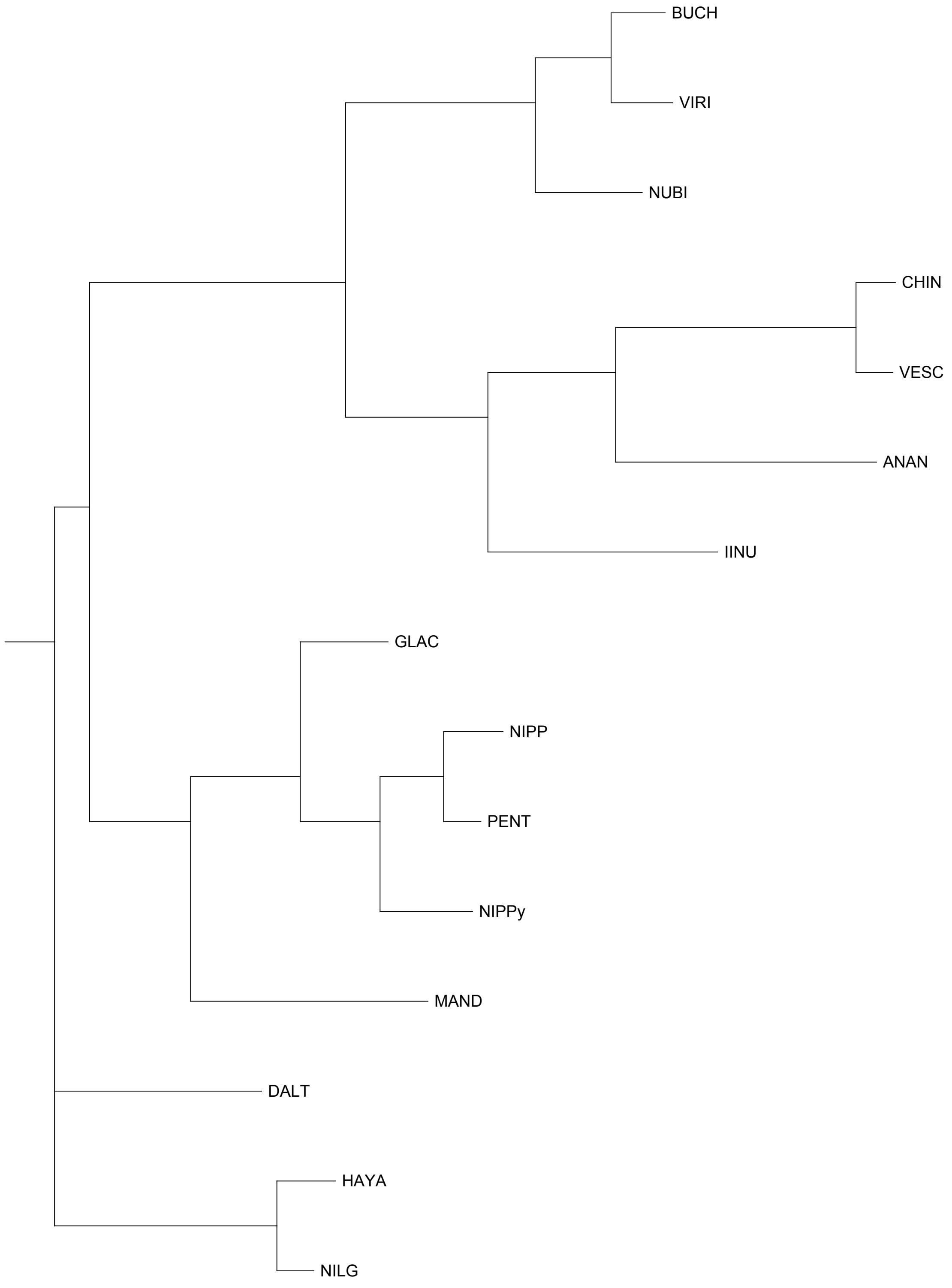


FAN_r2.3ch7X2b
