## Supplementary Figure S2 for "A chromosome-scale strawberry genome assembly of a Japanese variety, Reikou"

Chromosome-level synteny in the genomes of *F.* × *ananassa* (FAN_r2.3.pseudomolecule) and *F. vesca* (F. vesca v1.1)

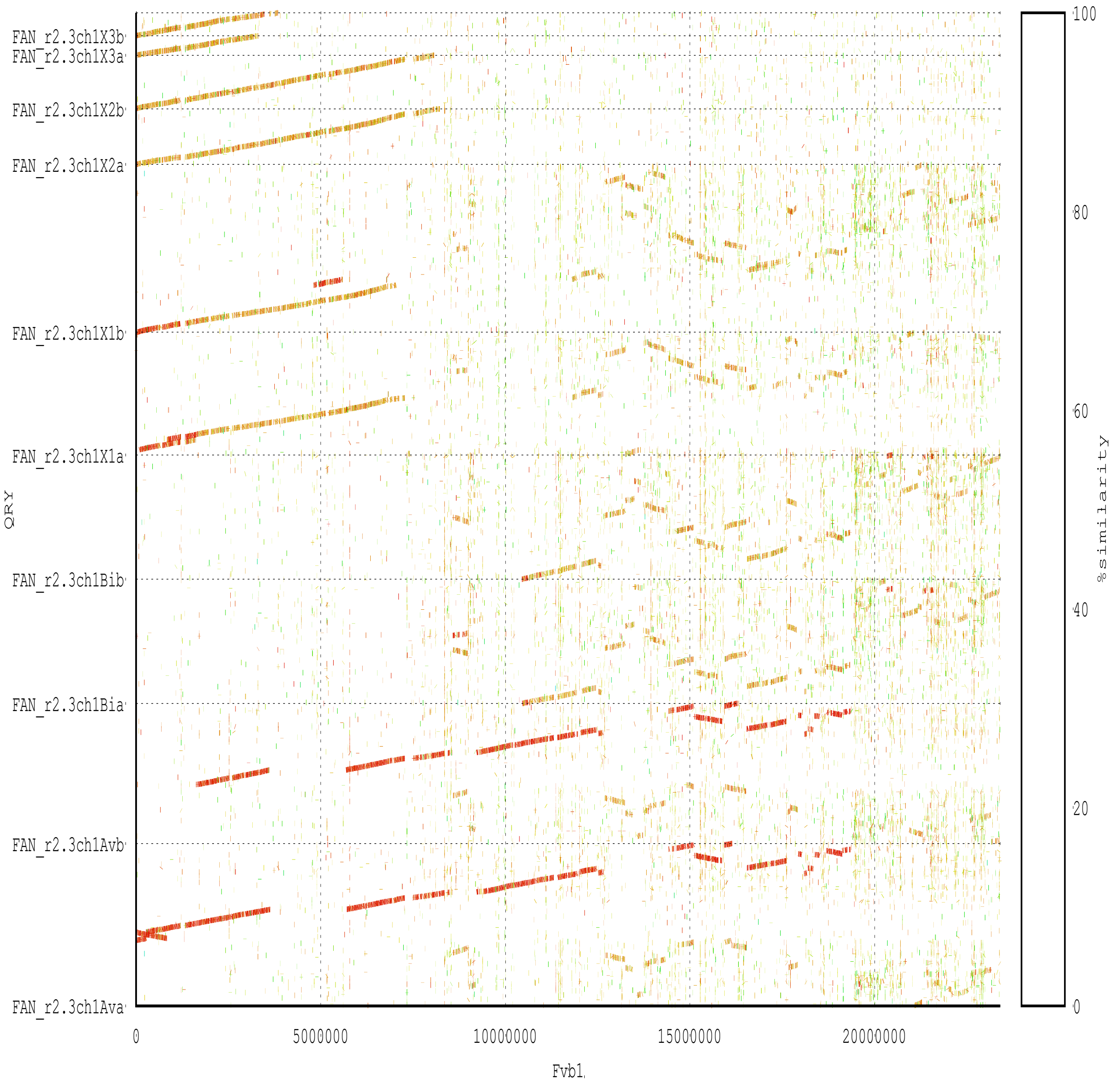

FAN_r2.3ch1X3b
FAN_r2.3ch1X3a
FAN_r2.3ch1X2b
FAN_r2.3ch1X2a
FAN_r2.3ch1X1b
FAN_r2.3ch1X1a
FAN_r2.3ch1Bib
FAN_r2.3ch1Bia
FAN_r2.3ch1Avb
FAN_r2.3ch1Ava
QRY
0
5000000
10000000
15000000
20000000
Fvb1
100
80
60
40
20
0
%similarity

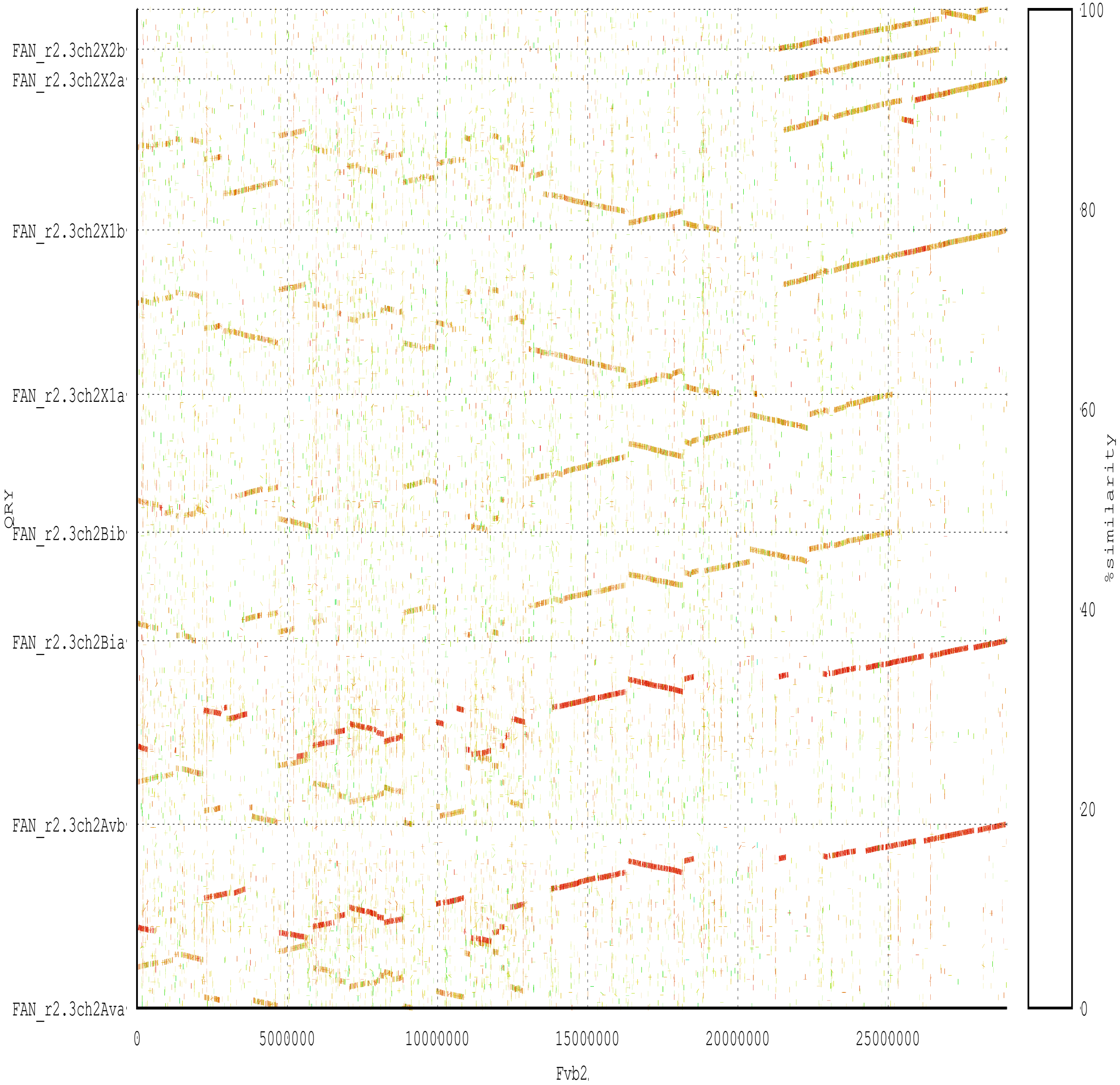
FAN_r2.3ch2X2b
FAN_r2.3ch2X2a
FAN_r2.3ch2X1b
FAN_r2.3ch2X1a
FAN_r2.3ch2Bib
FAN_r2.3ch2Bia
FAN_r2.3ch2Avb
FAN_r2.3ch2Ava
QRY
0
5000000
10000000
15000000
20000000
25000000
Fvb2
100
80
60
40
20
0
%similarity

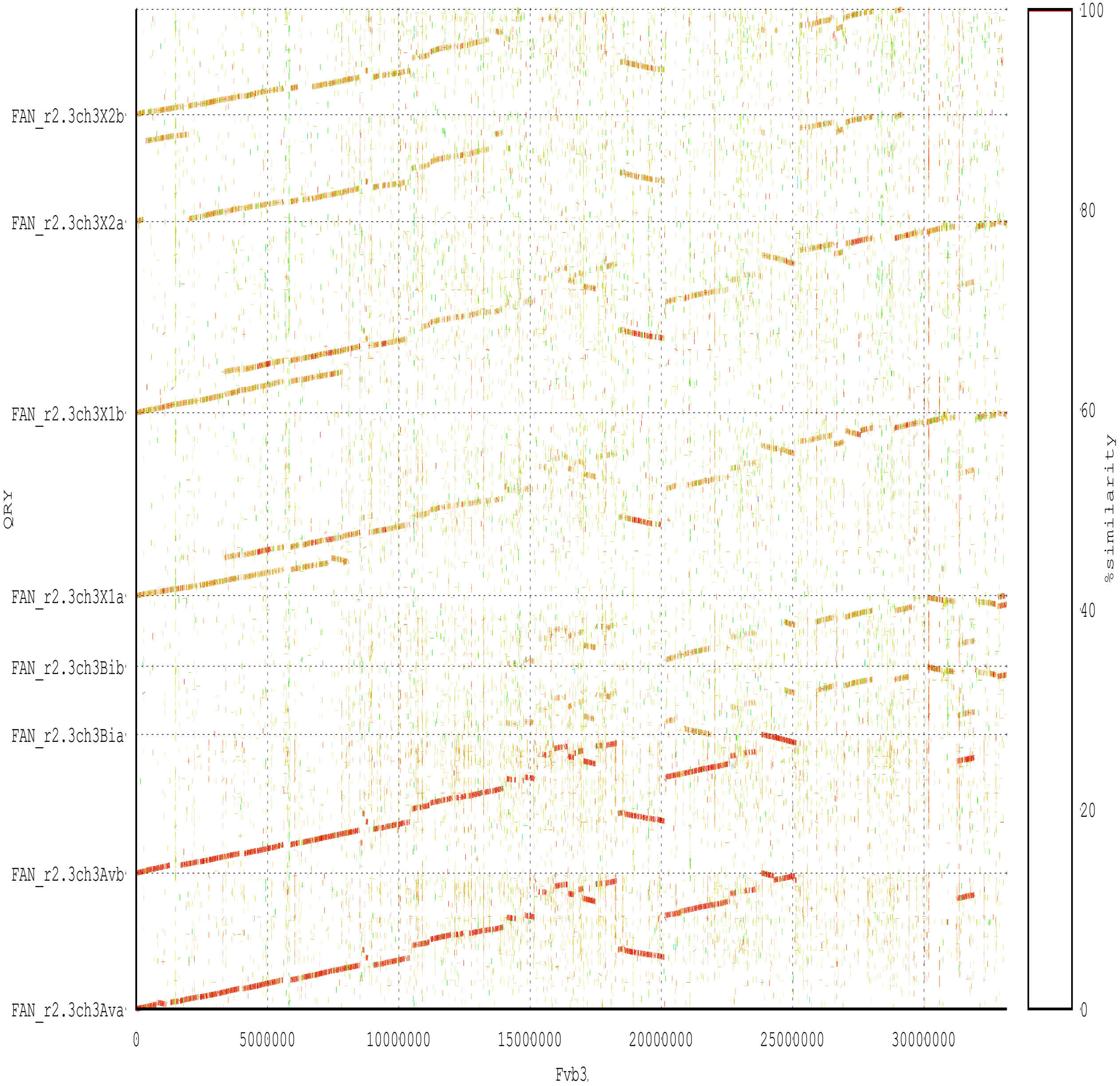

FAN_r2.3ch3X2b
FAN_r2.3ch3X2a
FAN_r2.3ch3X1b
FAN_r2.3ch3X1a
FAN_r2.3ch3Bib
FAN_r2.3ch3Bia
FAN_r2.3ch3Avb
FAN_r2.3ch3Ava
QRY
0
5000000
10000000
15000000
20000000
25000000
30000000
Fvb3
100
80
60
40
20
0
%similarity

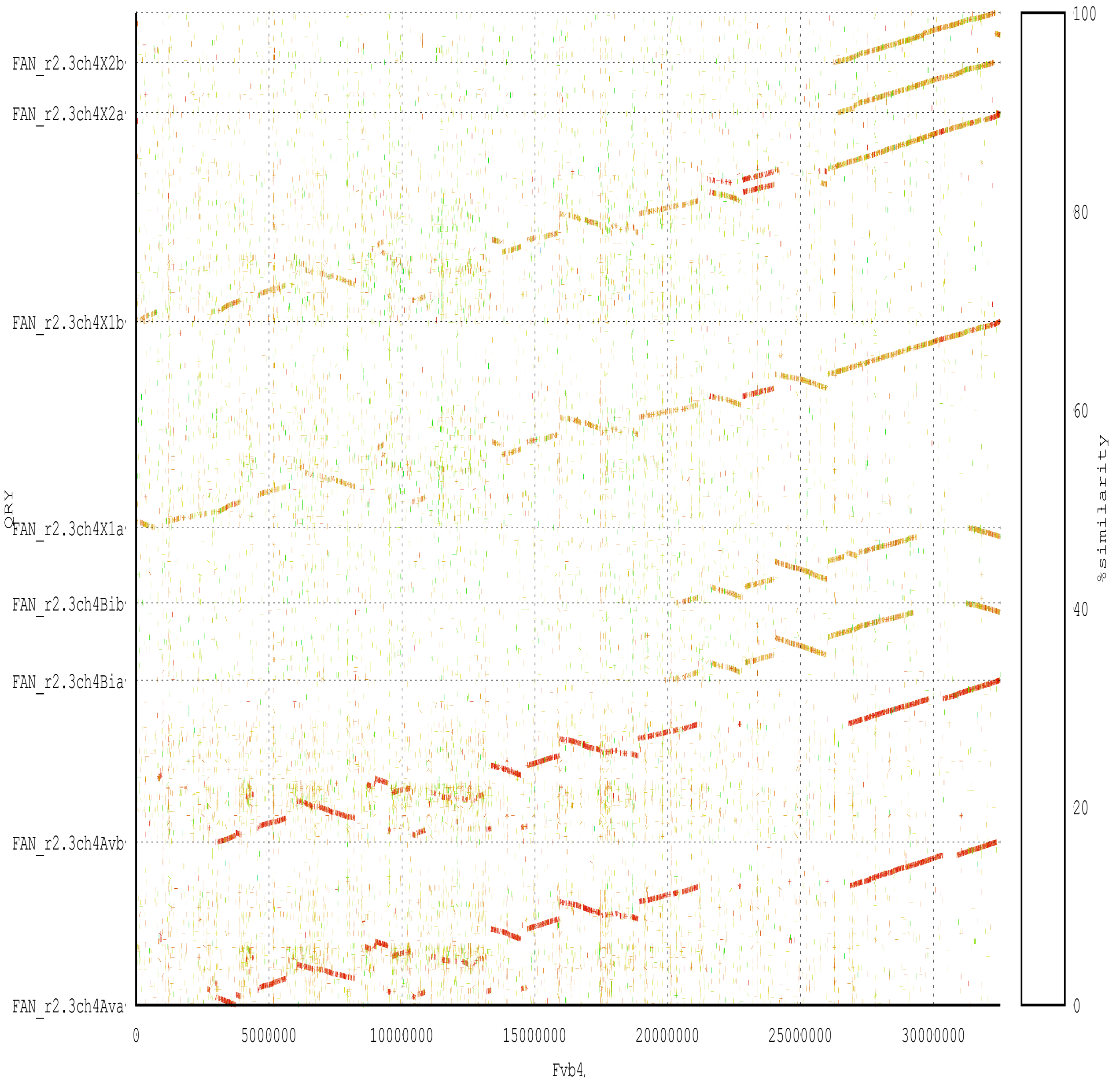
FAN_r2.3ch4X2b
FAN_r2.3ch4X2a
FAN_r2.3ch4X1b
FAN_r2.3ch4X1a
FAN_r2.3ch4Bib
FAN_r2.3ch4Bia
FAN_r2.3ch4Avb
FAN_r2.3ch4Ava
QRY
0
5000000
10000000
15000000
20000000
25000000
30000000
Fvb4
100
80
60
40
20
0
%similarity

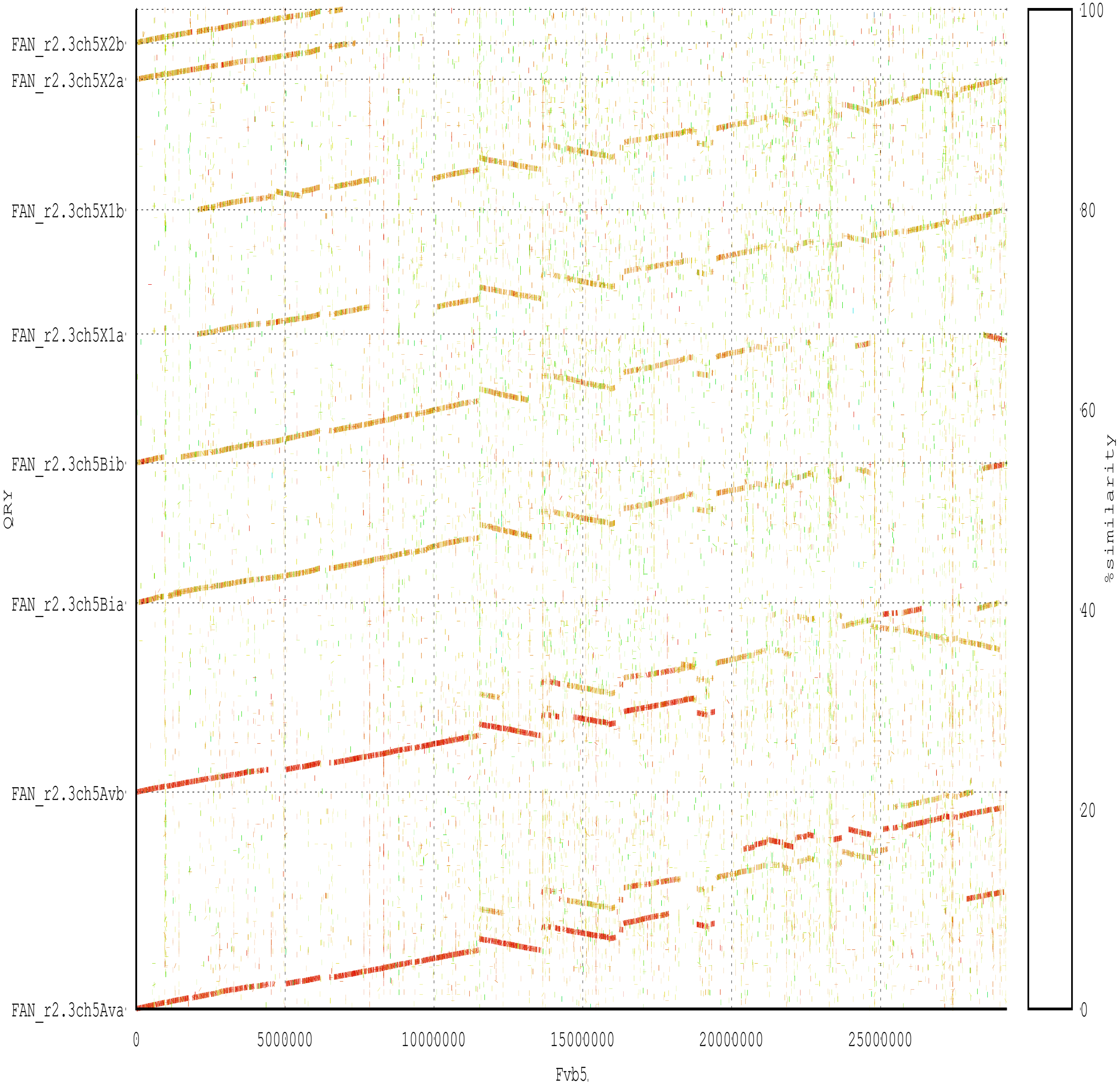

FAN_r2.3ch5X2b
FAN_r2.3ch5X2a
FAN_r2.3ch5X1b
FAN_r2.3ch5X1a
FAN_r2.3ch5Bib
FAN_r2.3ch5Bia
FAN_r2.3ch5Avb
FAN_r2.3ch5Ava
QRY
0
5000000
10000000
15000000
20000000
25000000
Fvb5
100
80
60
40
20
0
%similarity

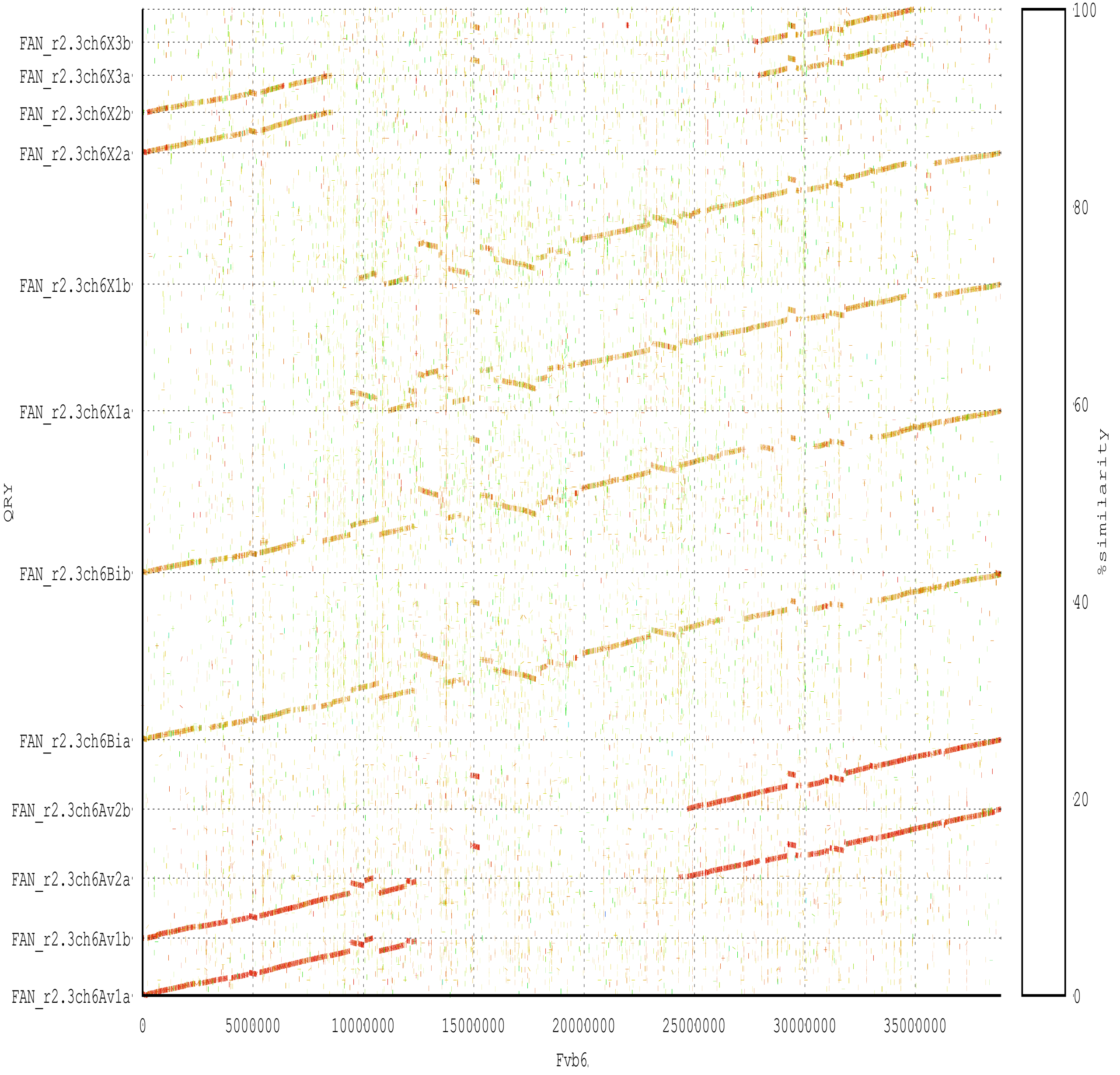
FAN_r2.3ch6X3b
FAN_r2.3ch6X3a
FAN_r2.3ch6X2b
FAN_r2.3ch6X2a
FAN_r2.3ch6X1b
FAN_r2.3ch6X1a
FAN_r2.3ch6Bib
FAN_r2.3ch6Bia
FAN_r2.3ch6Av2b
FAN_r2.3ch6Av2a
FAN_r2.3ch6Av1b
FAN_r2.3ch6Av1a
QRY
0
5000000
10000000
15000000
20000000
25000000
30000000
35000000
Fvb6
100
80
60
40
20
0
%similarity

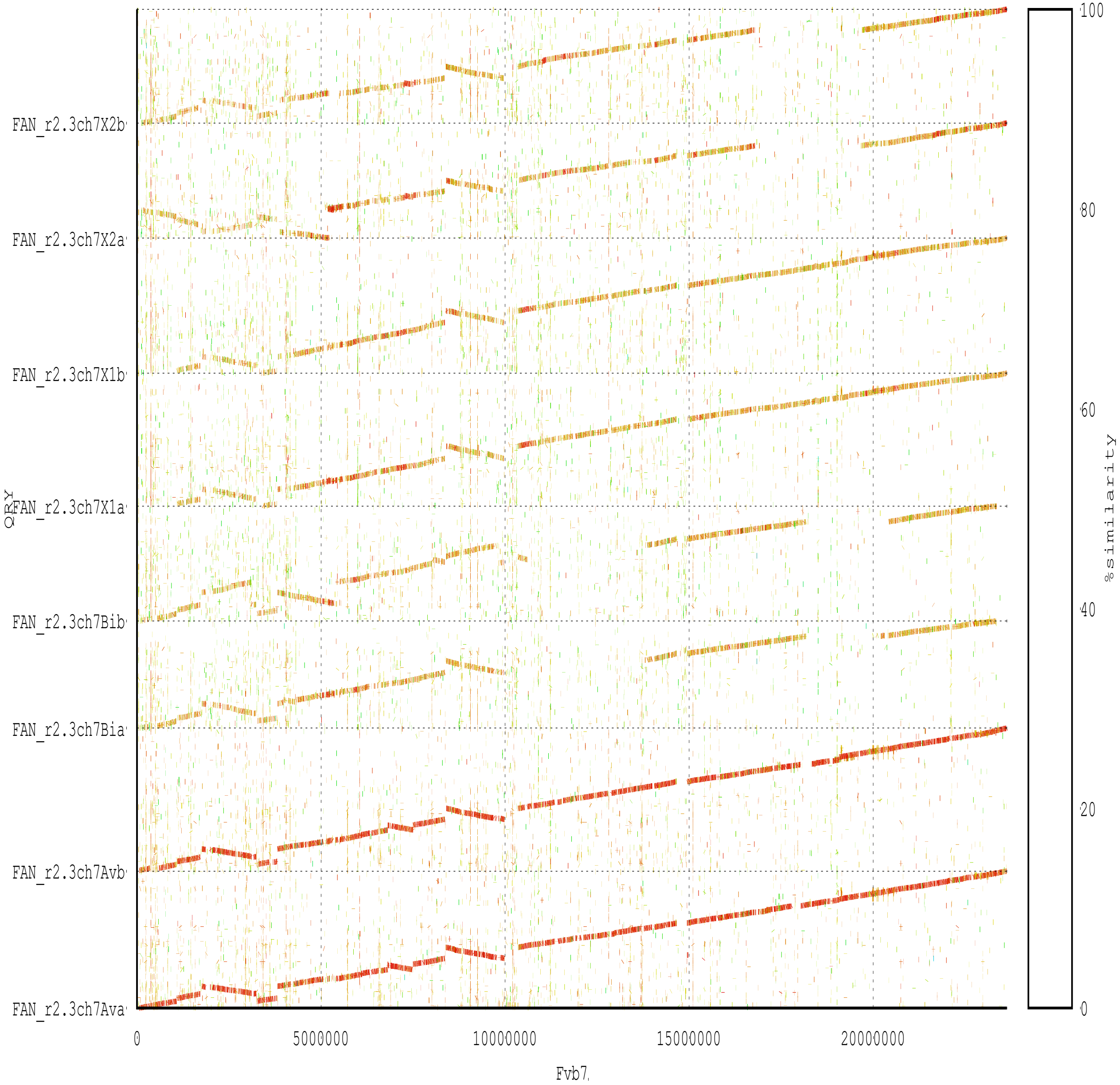
FAN_r2.3ch7X2b
FAN_r2.3ch7X2a
FAN_r2.3ch7X1b
FAN_r2.3ch7X1a
FAN_r2.3ch7Bib
FAN_r2.3ch7Bia
FAN_r2.3ch7Avb
FAN_r2.3ch7Ava
QRY
0
5000000
10000000
15000000
20000000
Fvb7
100
80
60
40
20
0
%similarity
