## Supplementary Figure S3 for "A chromosome-scale strawberry genome assembly of a Japanese variety, Reikou"

Chromosome-level synteny between two phased sequences of the *F.* × *ananassa* genome (FAN_r2.3.pseudomolecule)

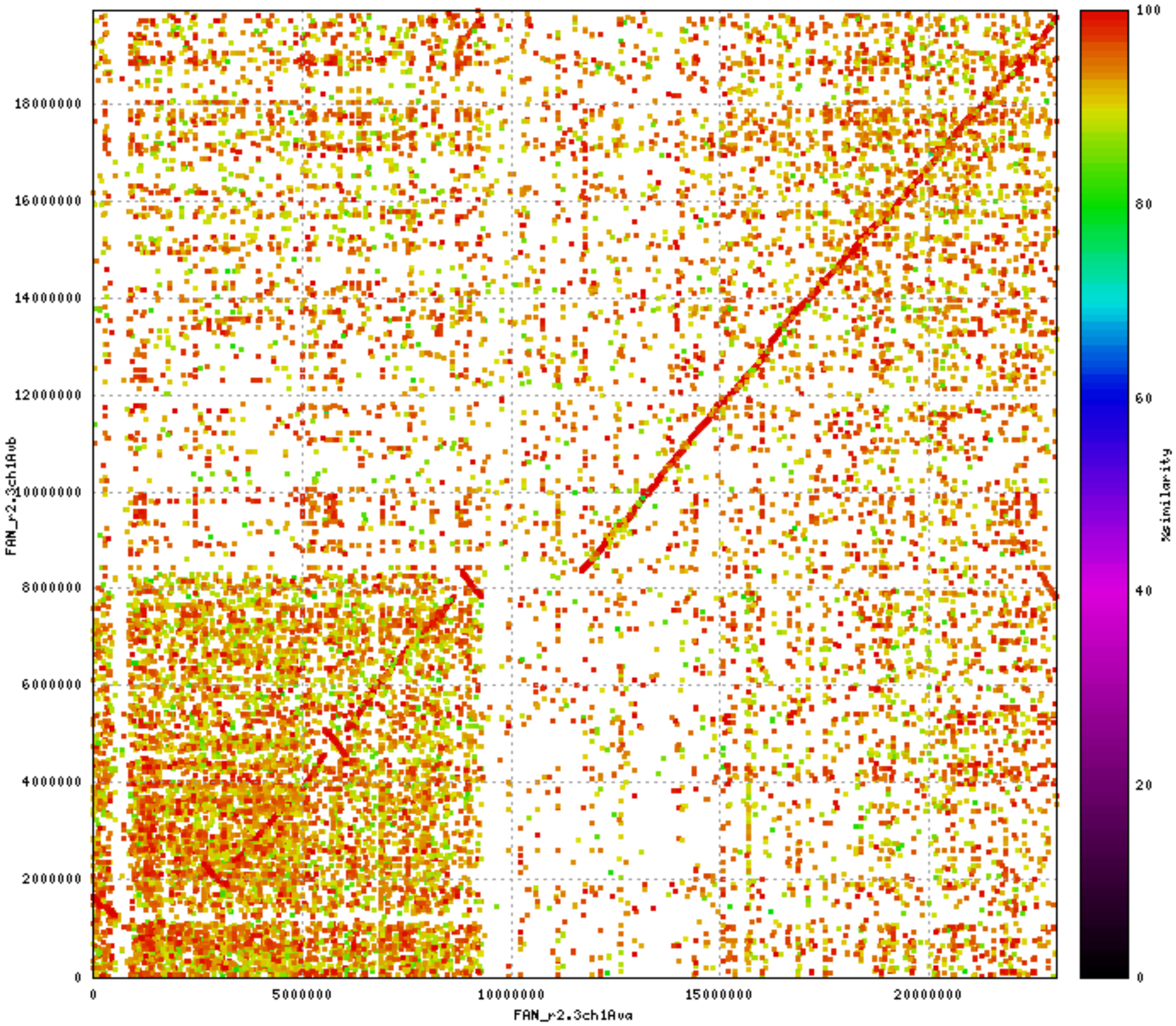

18000000
16000000
14000000
12000000
10000000
8000000
6000000
4000000
2000000
0
FAN_r2.3ch1Avb
0
5000000
10000000
15000000
20000000
FAN_r2.3ch1Ava
100
80
60
40
20
0
%similarity

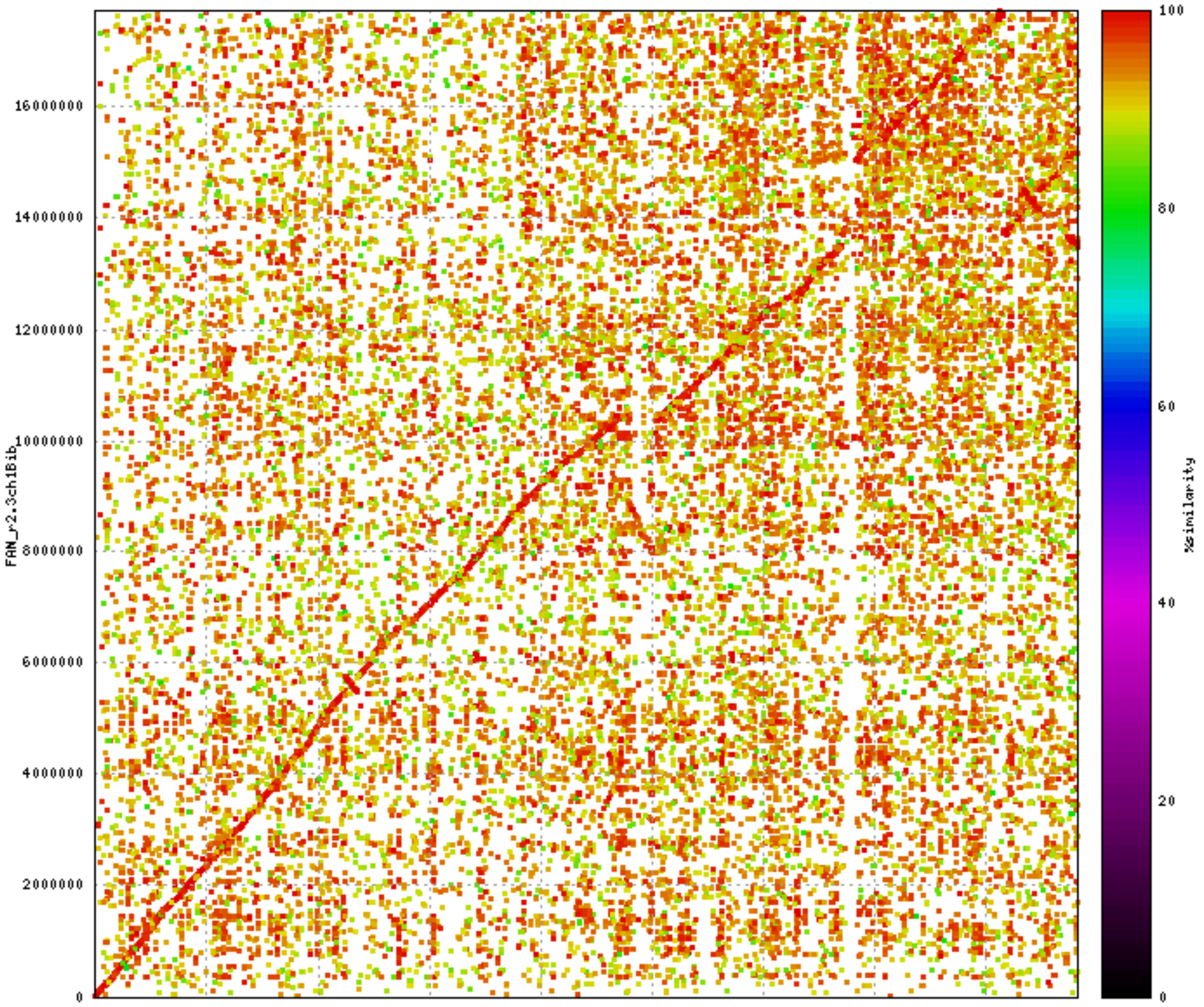

FAN_r2.3ch1Bib
FAN_r2.3ch1Bia
%similarity
0
2000000
4000000
6000000
8000000
10000000
12000000
14000000
16000000
20
40
60
80
100

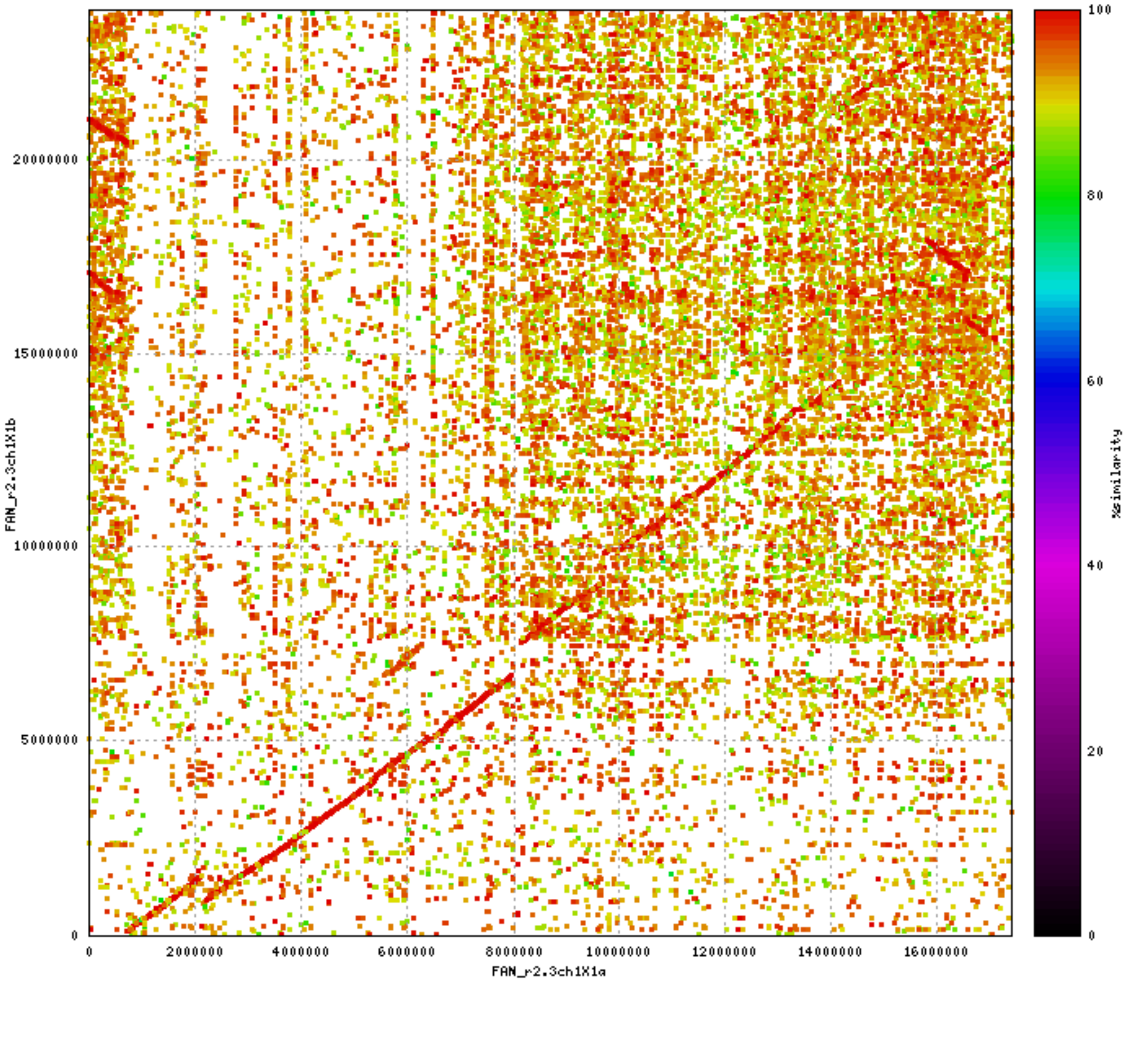

FAN_r2.3ch1X1b
FAN_r2.3ch1X1a
%similarity
0
5000000
10000000
15000000
20000000
2000000
4000000
6000000
8000000
10000000
12000000
14000000
16000000
20
40
60
80
100

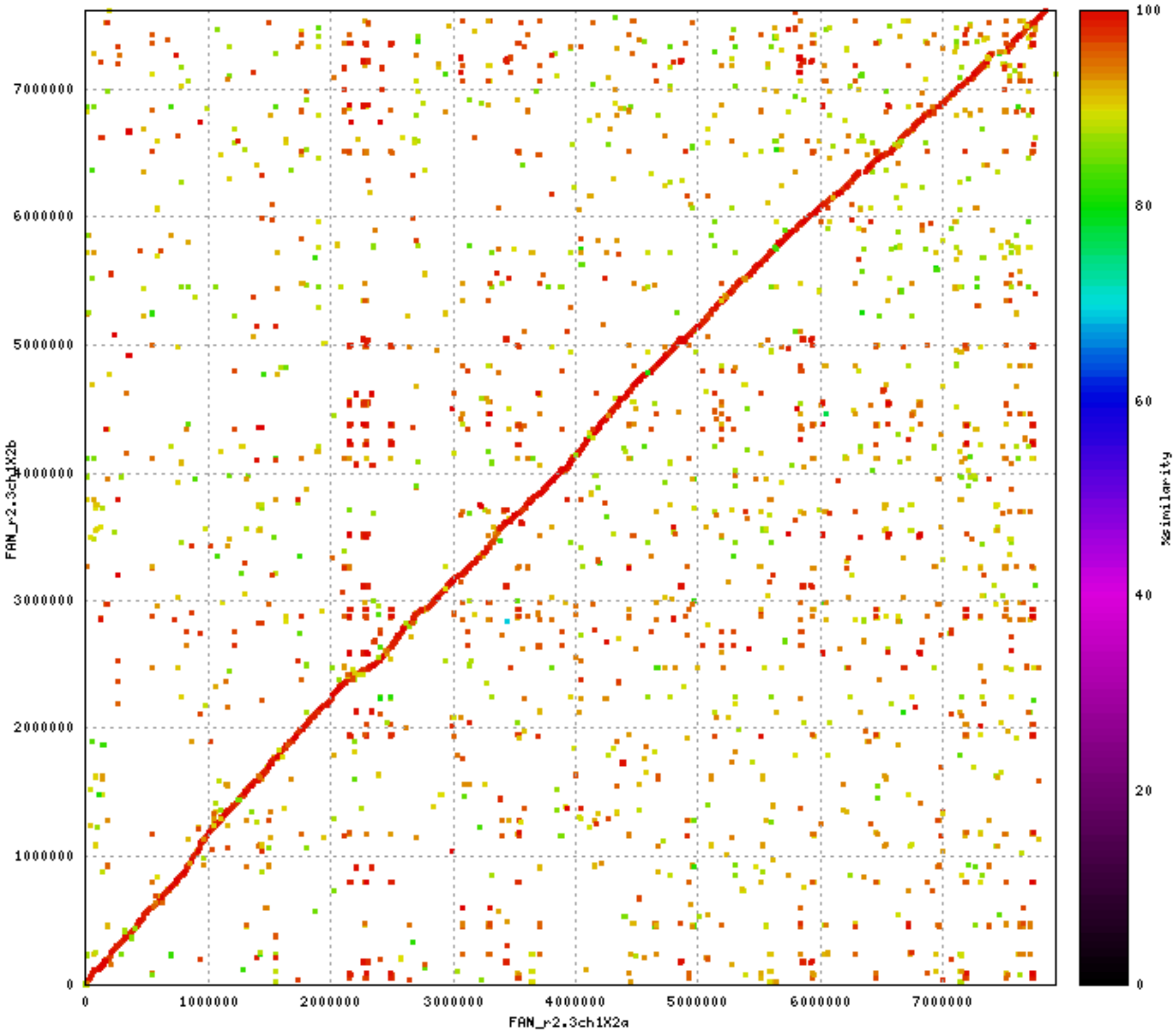

FAN_r2.3ch1X2b
FAN_r2.3ch1X2a
0
1000000
2000000
3000000
4000000
5000000
6000000
7000000
%similarity
100
80
60
40
20
0

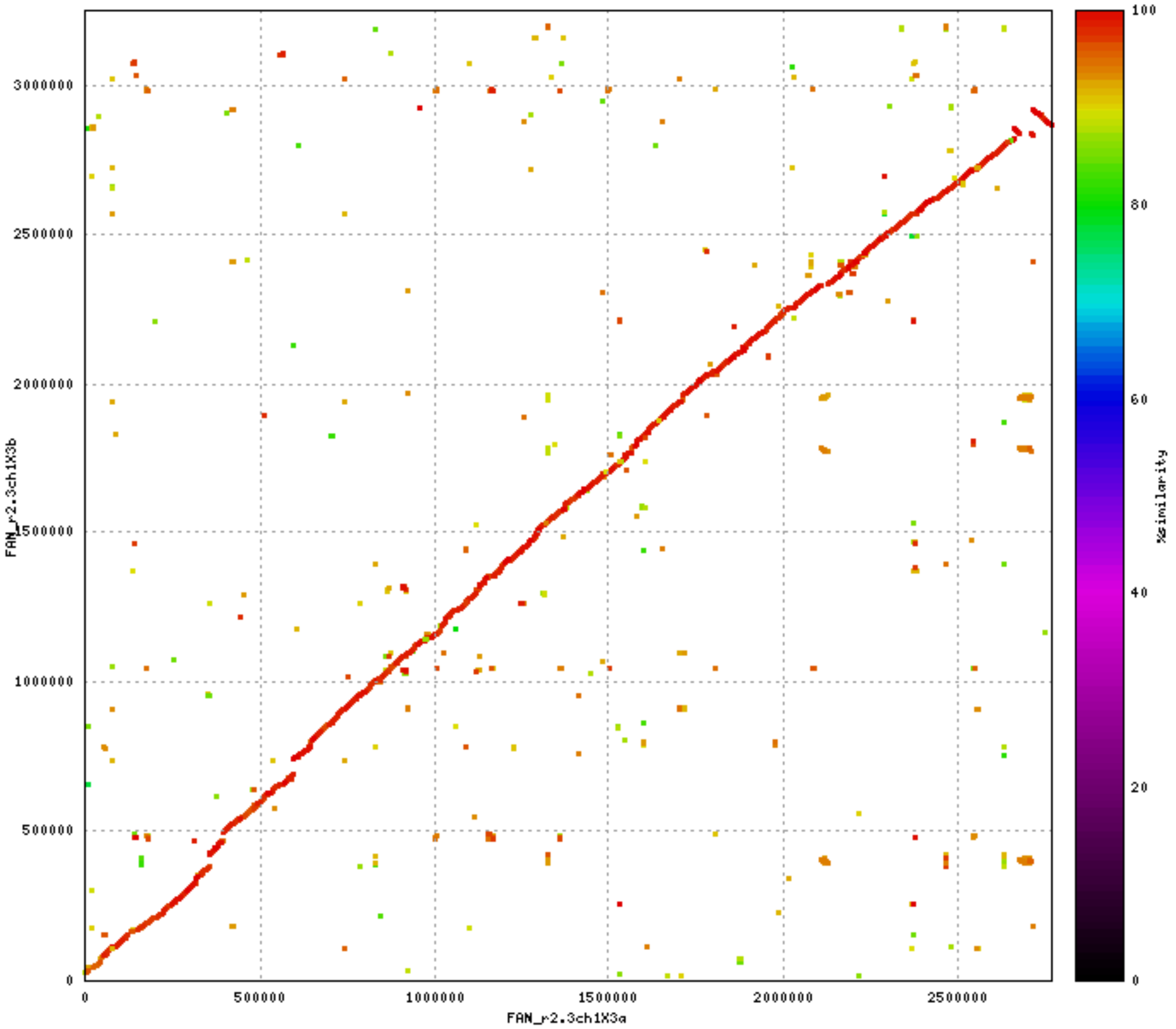

FAN_r2.3ch1X3b
FAN_r2.3ch1X3a
%similarity
0
500000
1000000
1500000
2000000
2500000
3000000
0
20
40
60
80
100

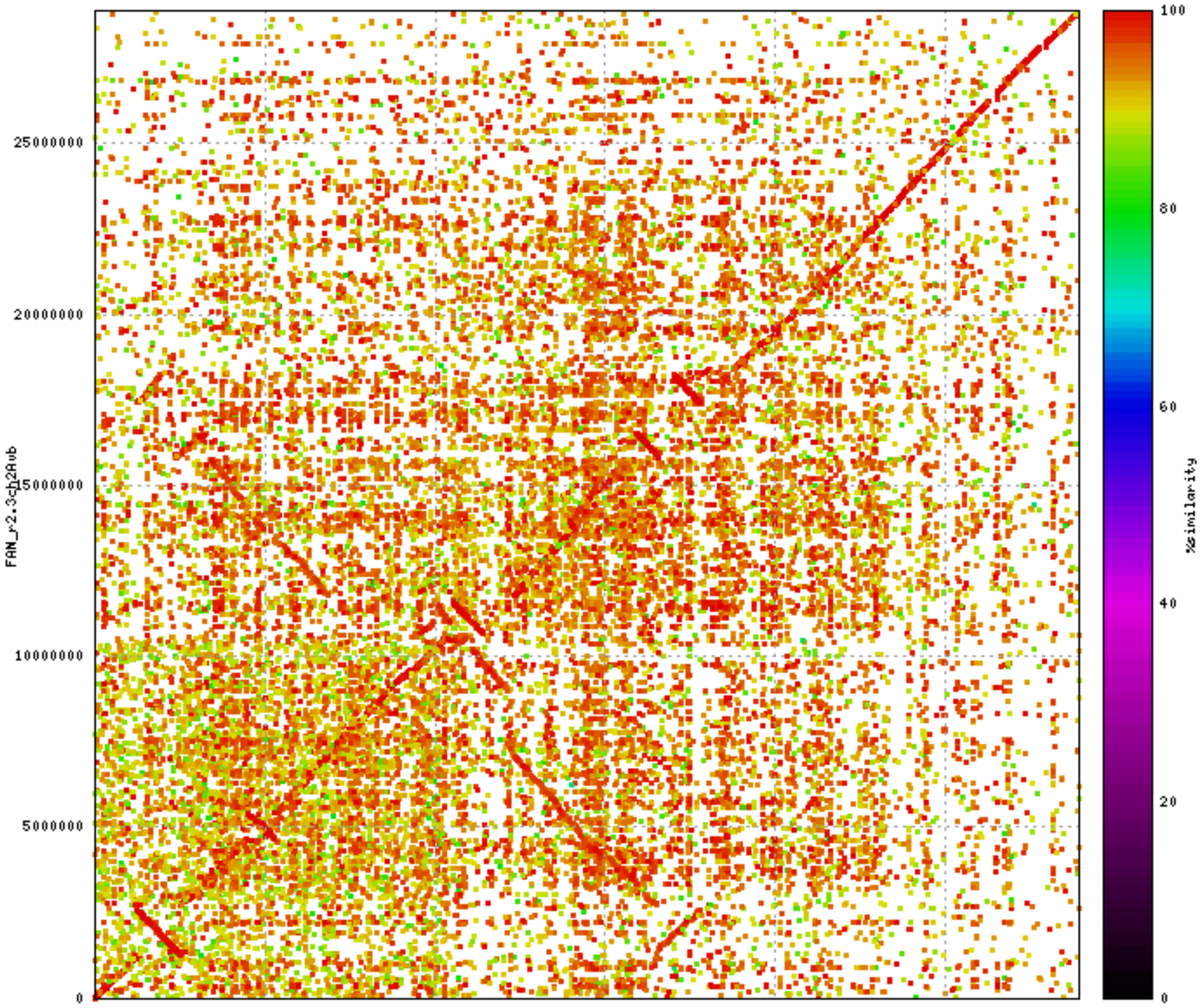

FAN_r2.3ch2Avb
FAN_r2.3ch2Ava
%similarity
0
5000000
10000000
15000000
20000000
25000000
0
20
40
60
80
100

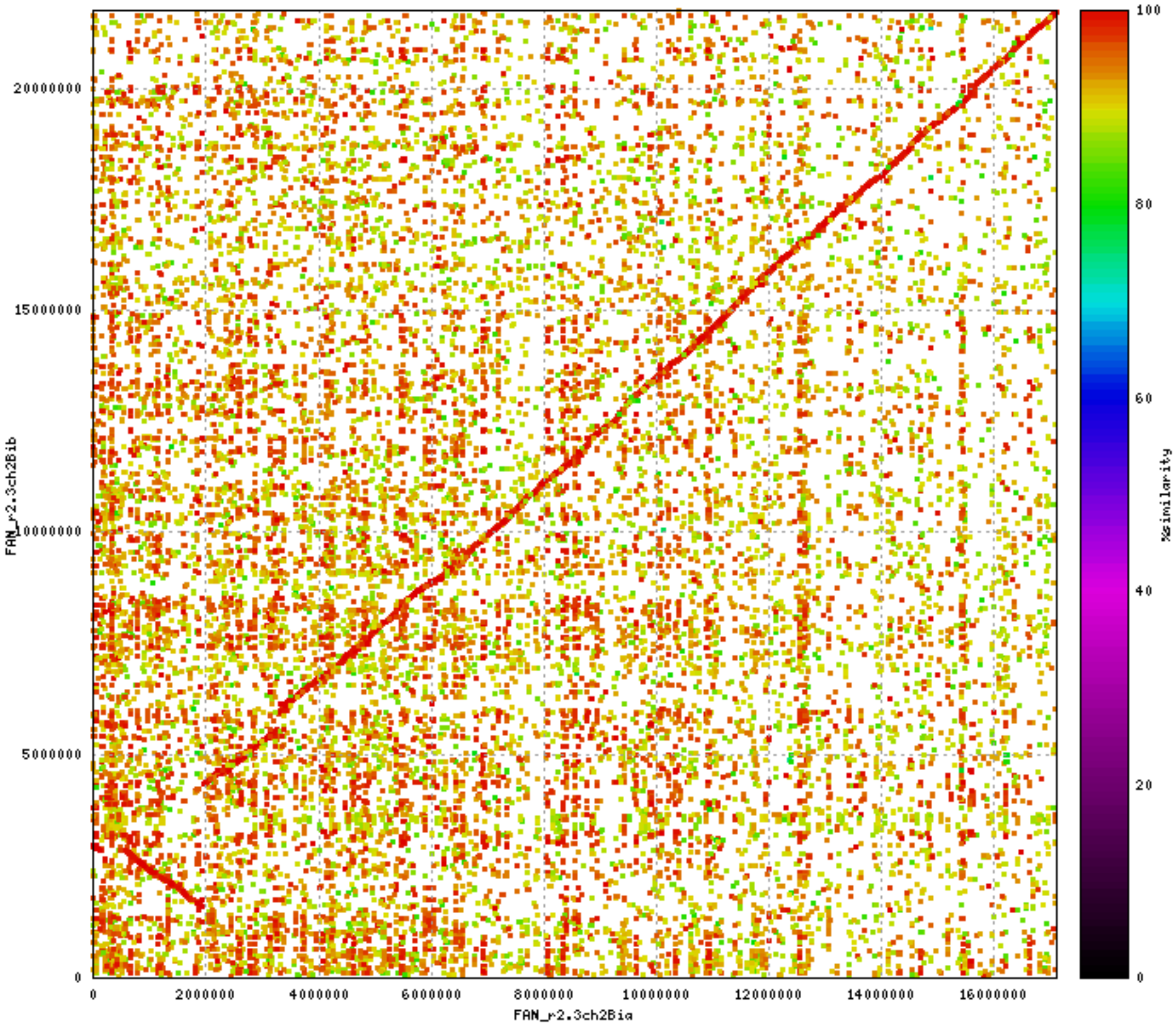

FAN_r2.3ch2Bib
FAN_r2.3ch2Bia
%similarity
0
5000000
10000000
15000000
20000000
2000000
4000000
6000000
8000000
10000000
12000000
14000000
16000000
20
40
60
80
100

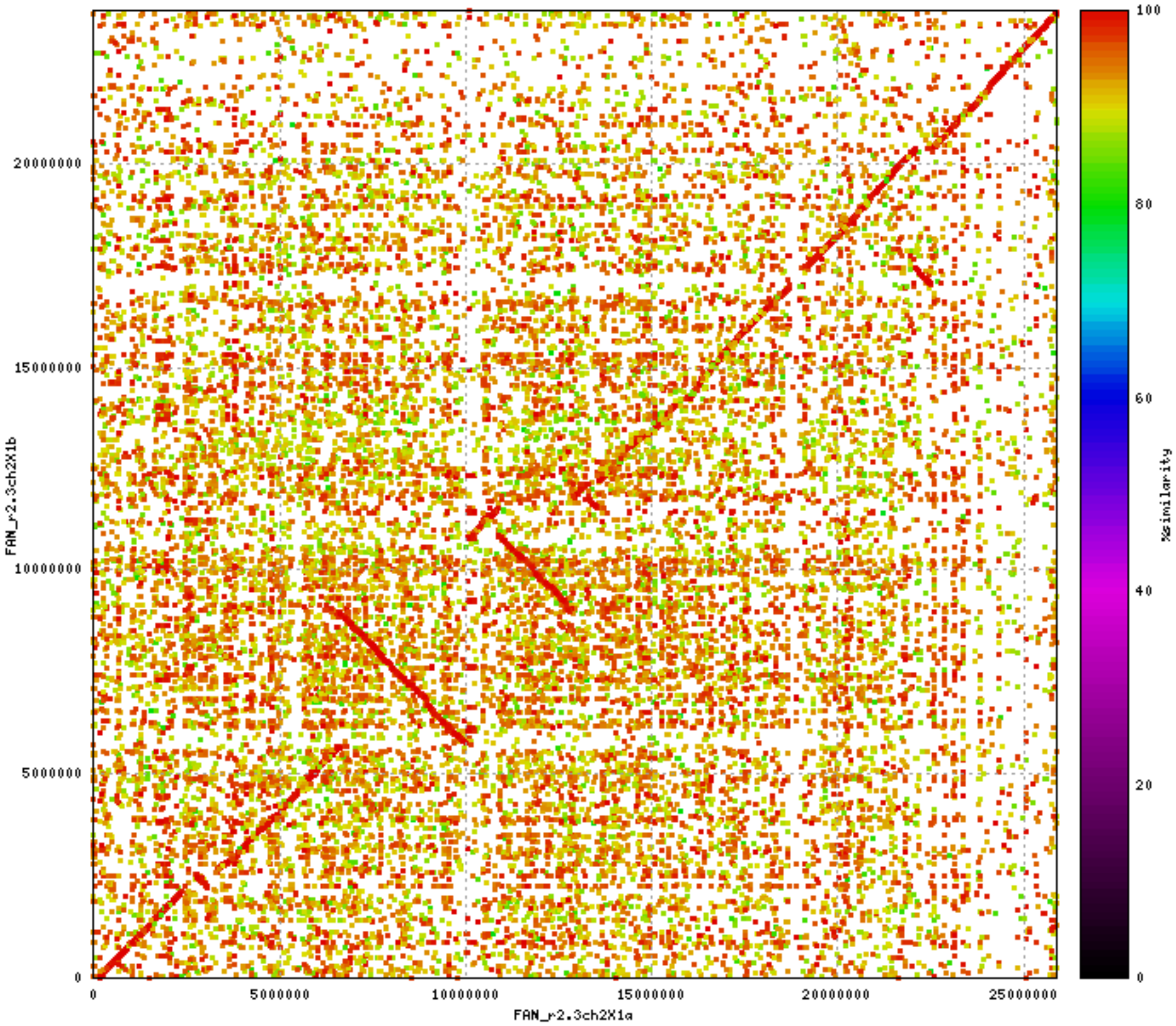

FAN_r2.3ch2X1b
FAN_r2.3ch2X1a
%similarity
0
5000000
10000000
15000000
20000000
25000000
0
20
40
60
80
100

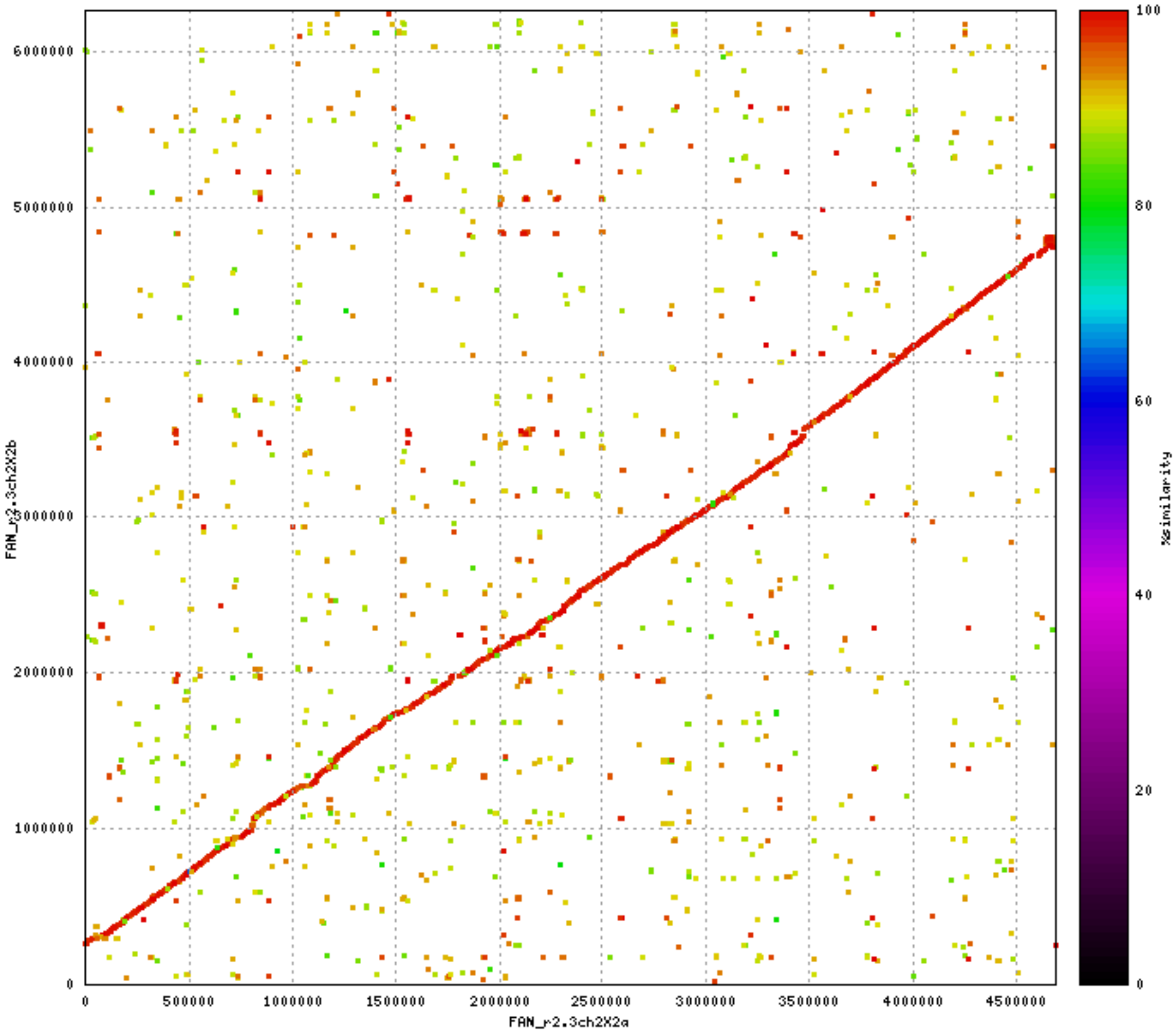

FAN_r2.3ch2X2b
FAN_r2.3ch2X2a
%similarity
0
500000
1000000
1500000
2000000
2500000
3000000
3500000
4000000
4500000
1000000
2000000
3000000
4000000
5000000
6000000
20
40
60
80
100

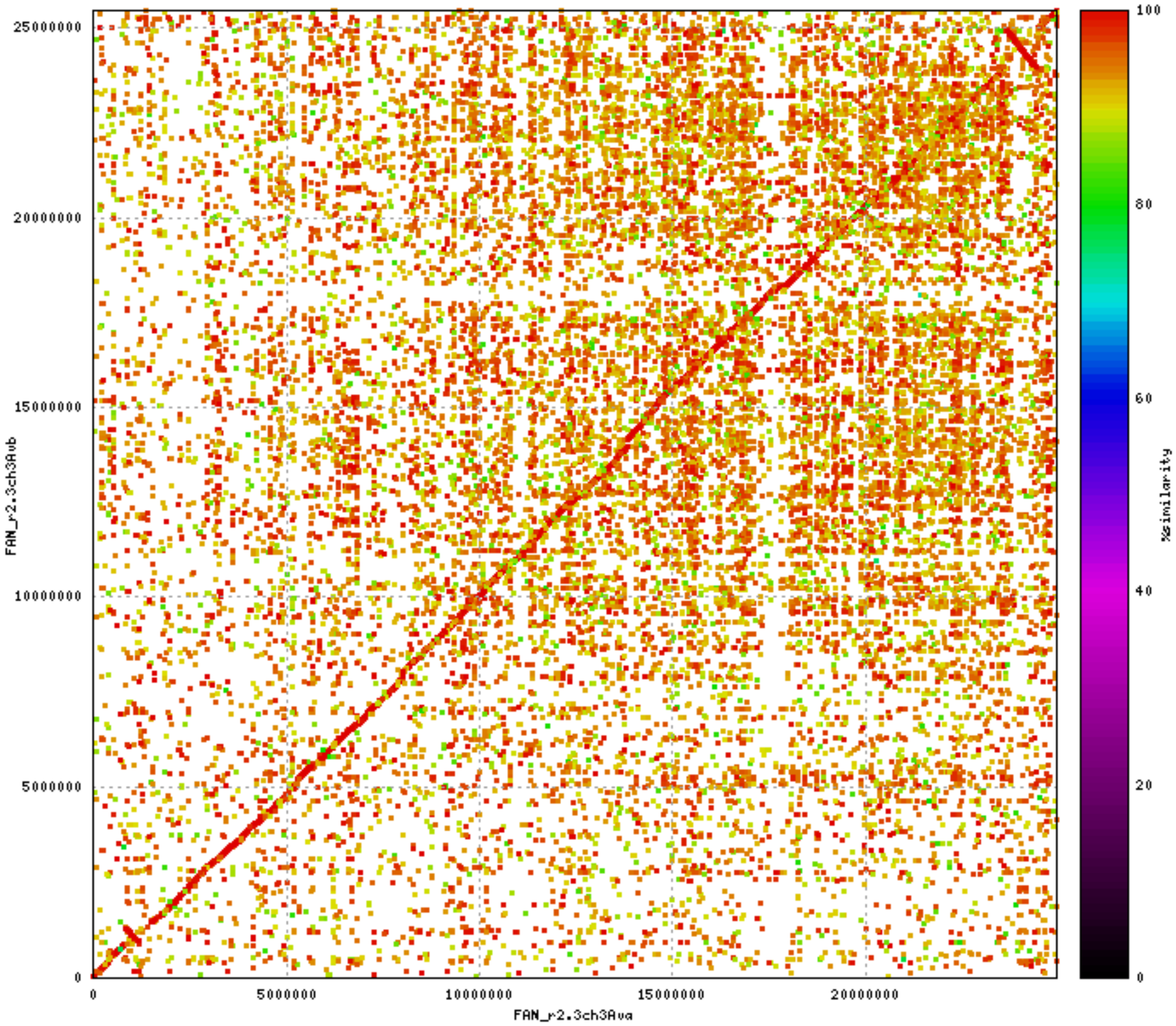

25000000
20000000
15000000
10000000
5000000
0
FAN_r2.3ch3Avb
0
5000000
10000000
15000000
20000000
FAN_r2.3ch3Ava
100
80
60
40
20
0
%similarity

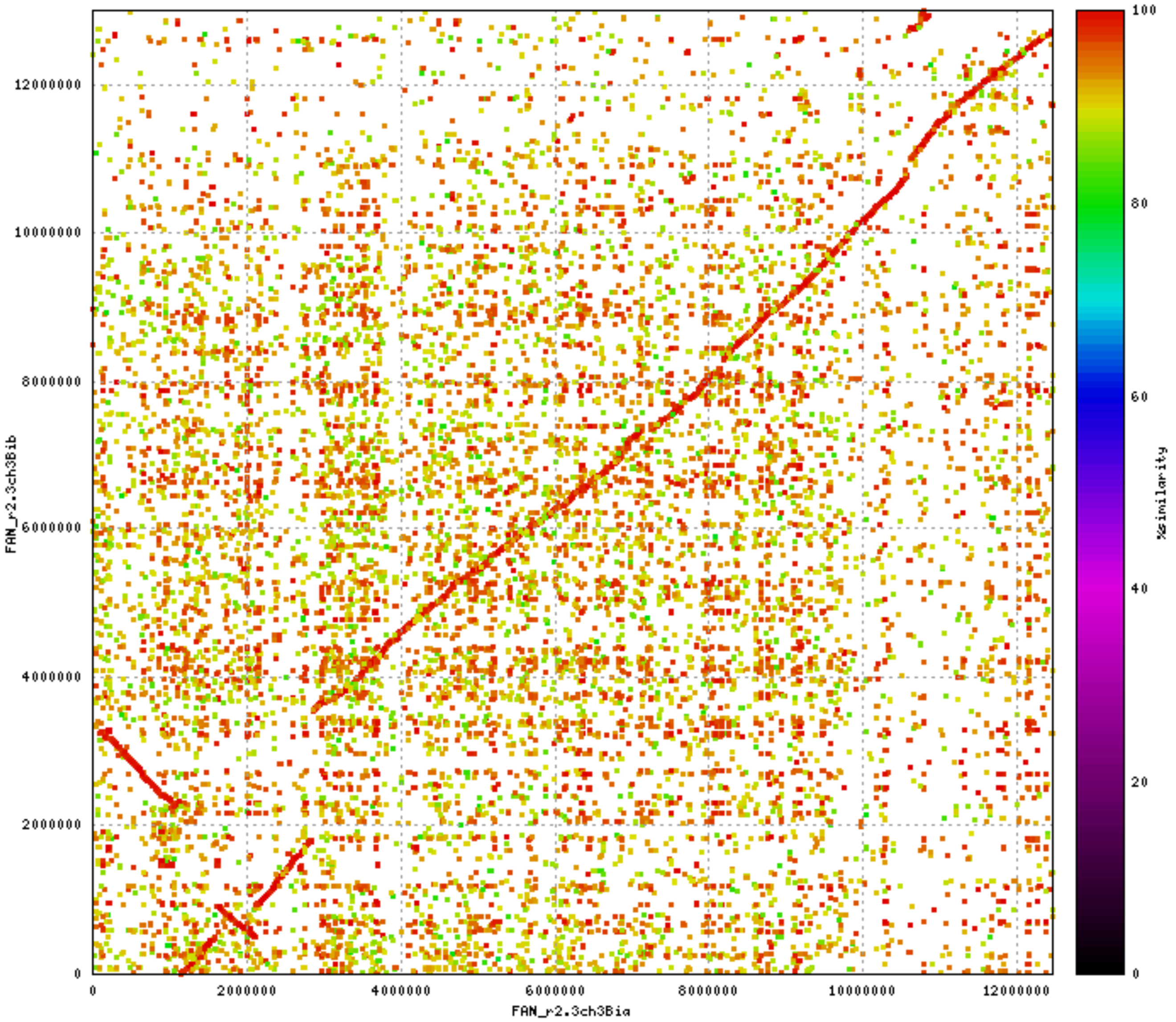

FAN_r2.3ch3Bib
FAN_r2.3ch3Bia
%similarity
0
2000000
4000000
6000000
8000000
10000000
12000000
0
20
40
60
80
100

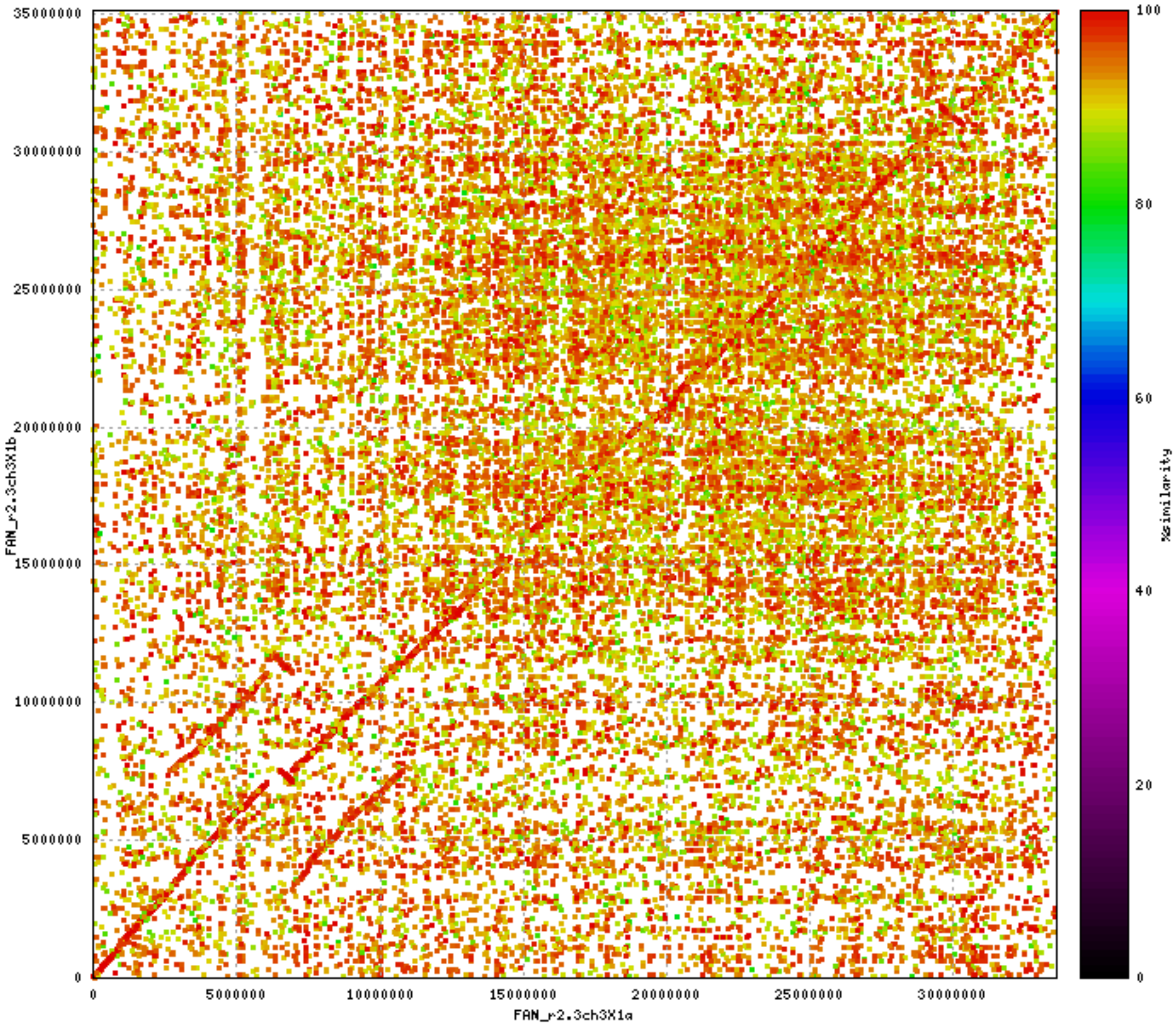

35000000
30000000
25000000
20000000
15000000
10000000
5000000
0
FAN_r2.3ch3X1b
0
5000000
10000000
15000000
20000000
25000000
30000000
FAN_r2.3ch3X1a
100
80
60
40
20
0
%similarity

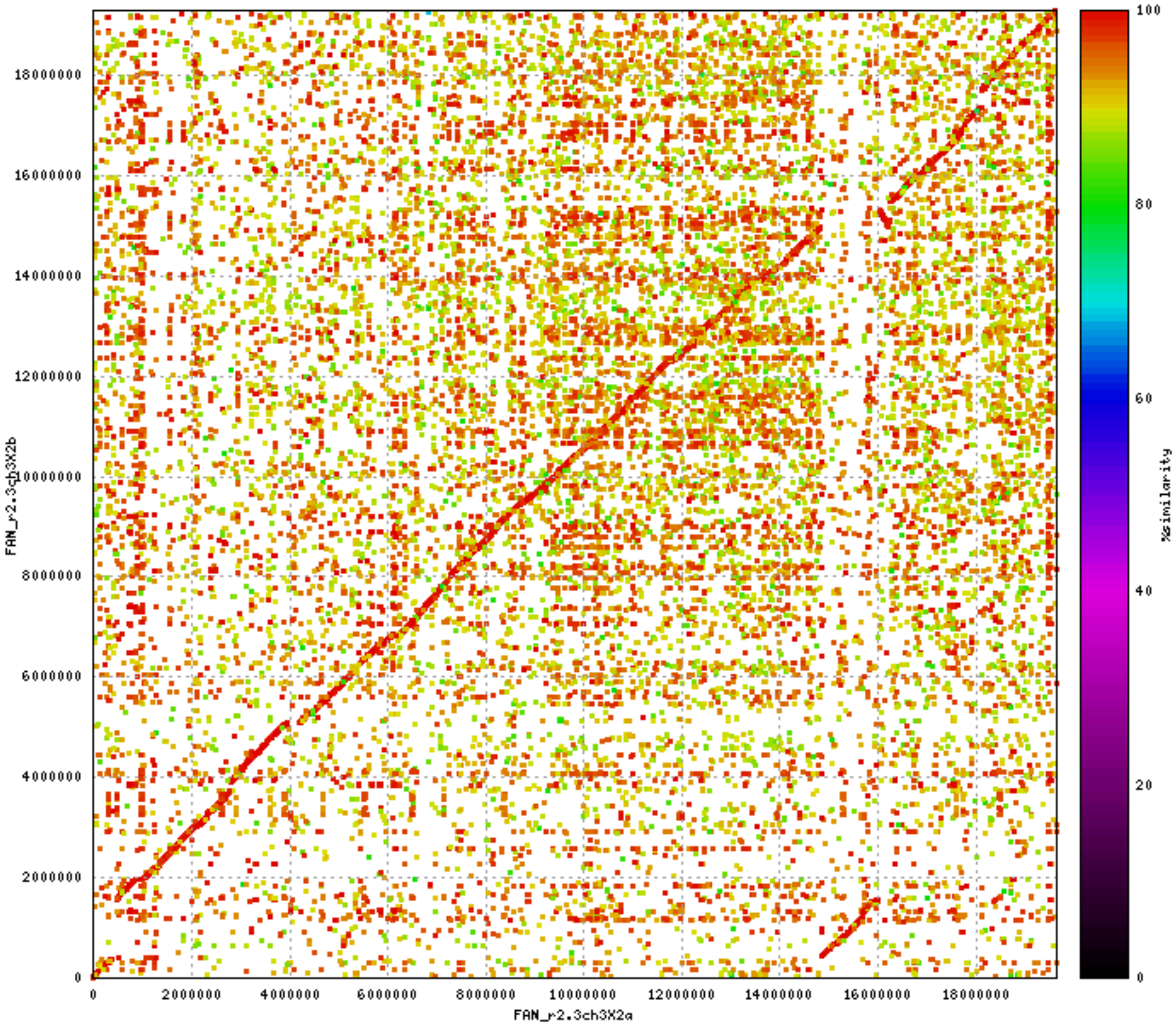

FAN_r2.3ch3X2b
FAN_r2.3ch3X2a
%similarity
0
2000000
4000000
6000000
8000000
10000000
12000000
14000000
16000000
18000000
0
20
40
60
80
100

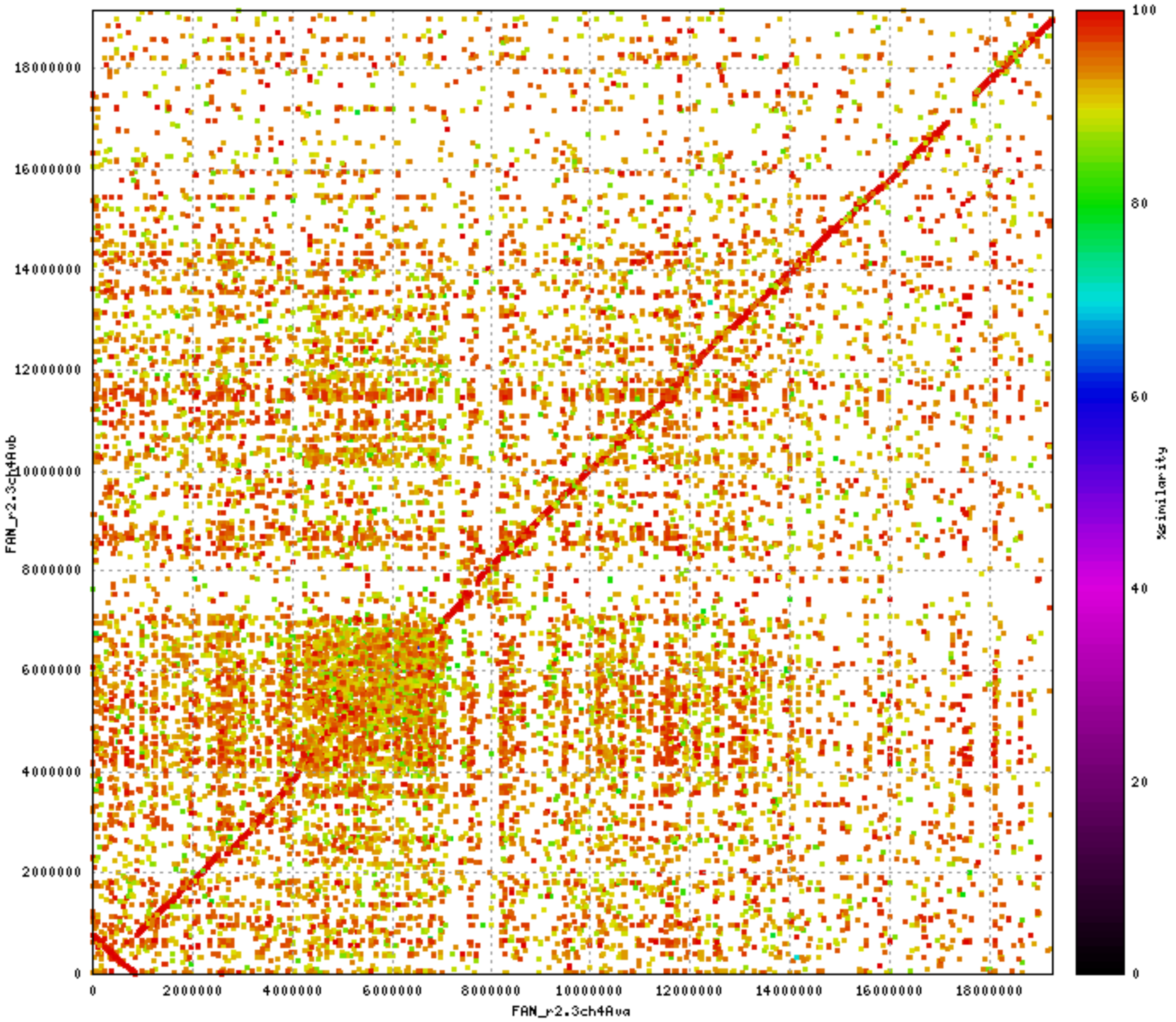

FAN_r2.3ch4Avb
FAN_r2.3ch4Ava
%similarity
0
2000000
4000000
6000000
8000000
10000000
12000000
14000000
16000000
18000000
20
40
60
80
100

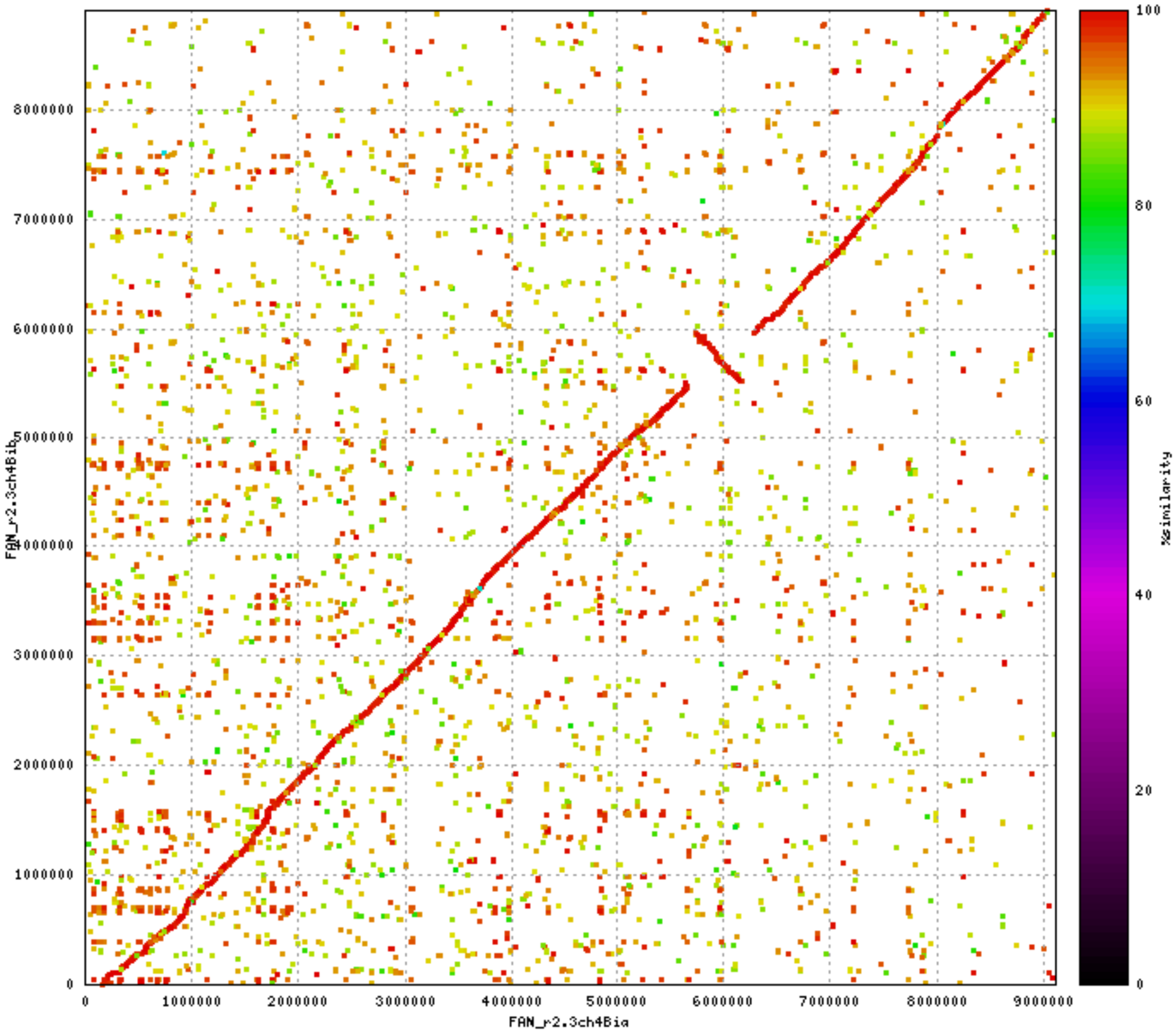

FAN_r2.3ch4Bib
FAN_r2.3ch4Bia
0
1000000
2000000
3000000
4000000
5000000
6000000
7000000
8000000
9000000
%similarity
0
20
40
60
80
100

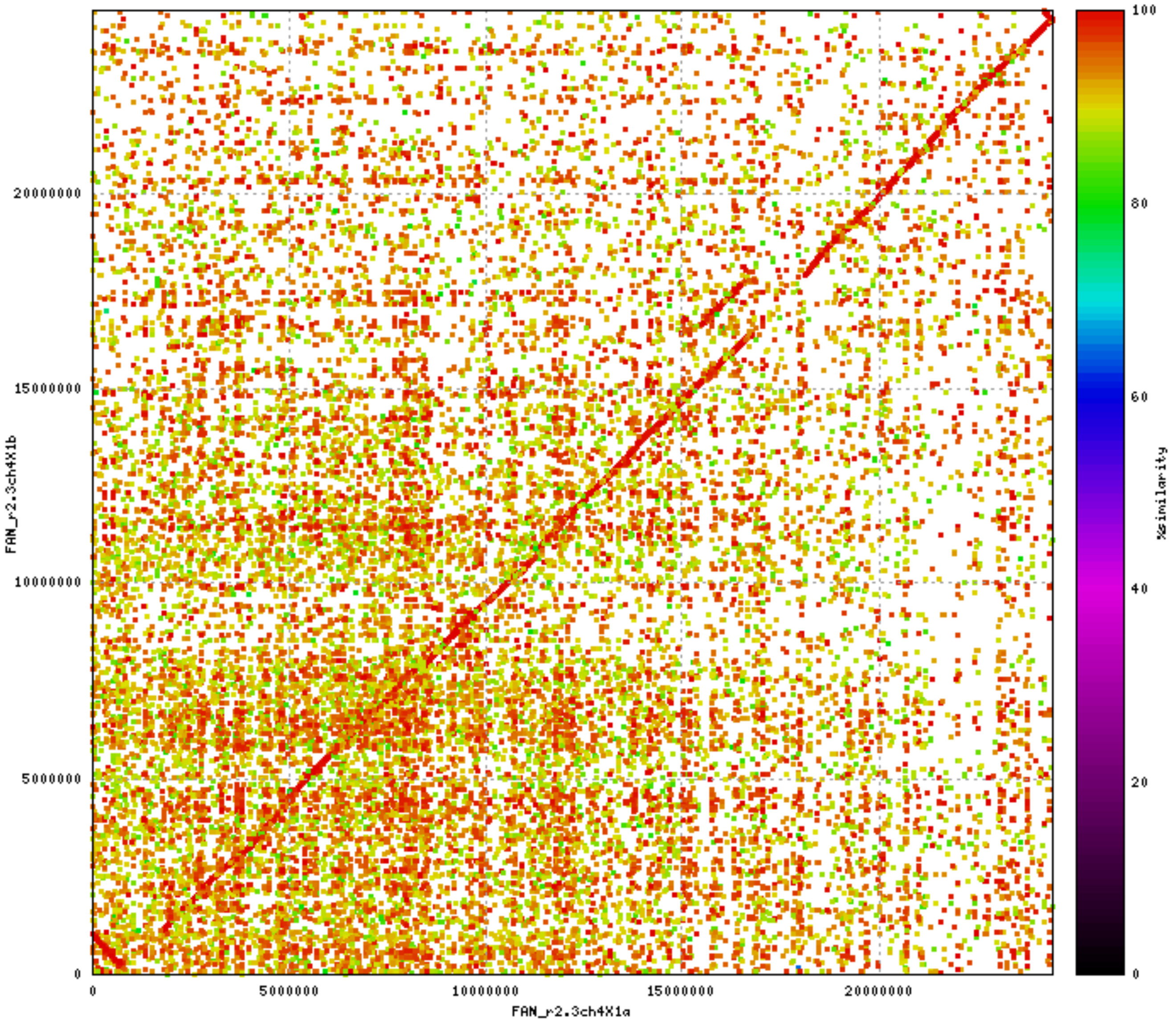
FAN_r2.3ch4X1b
FAN_r2.3ch4X1a
%similarity
0
5000000
10000000
15000000
20000000
0
20
40
60
80
100

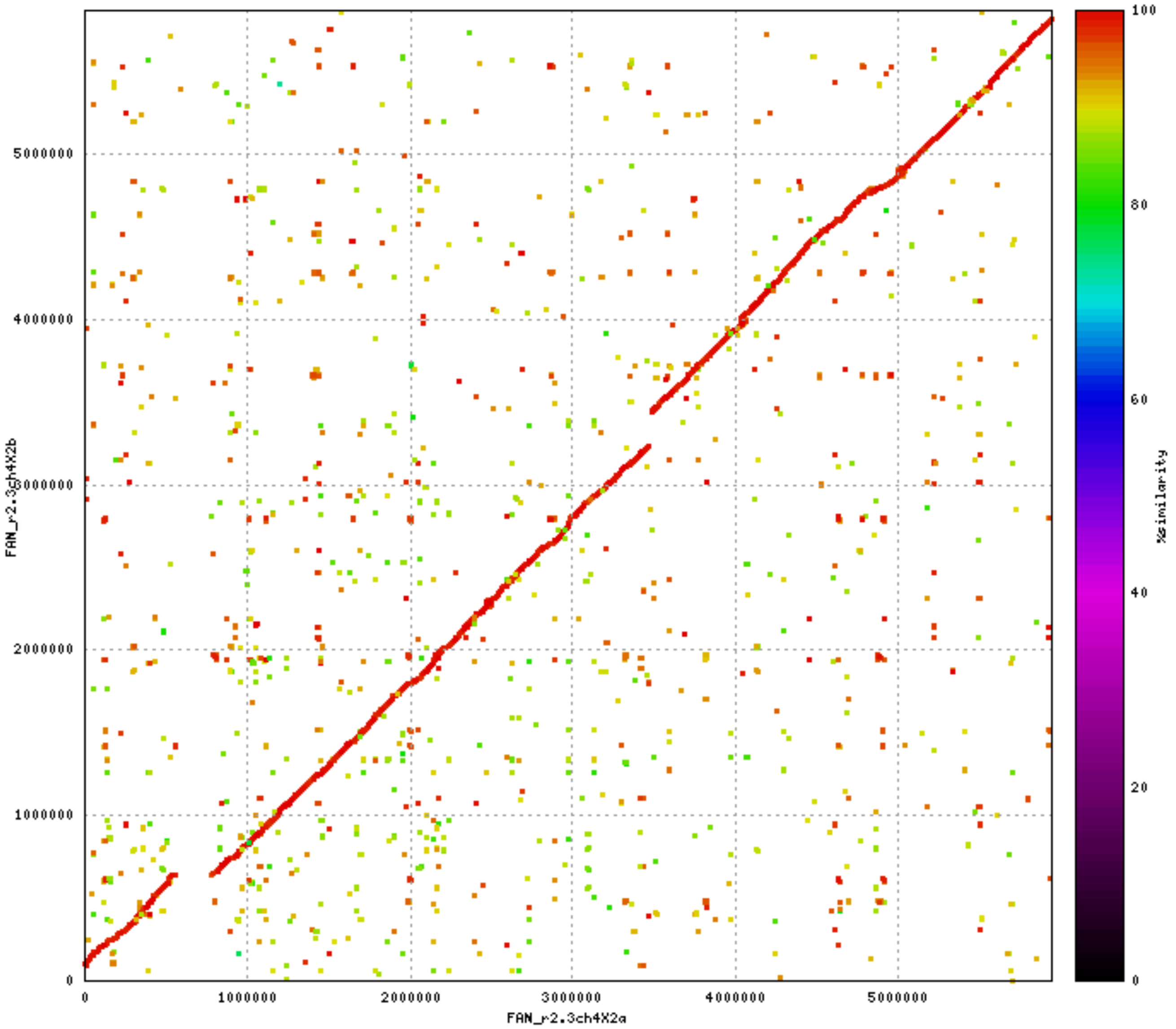

FAN_r2.3ch4X2b
FAN_r2.3ch4X2a
0
1000000
2000000
3000000
4000000
5000000
%similarity
100
80
60
40
20
0

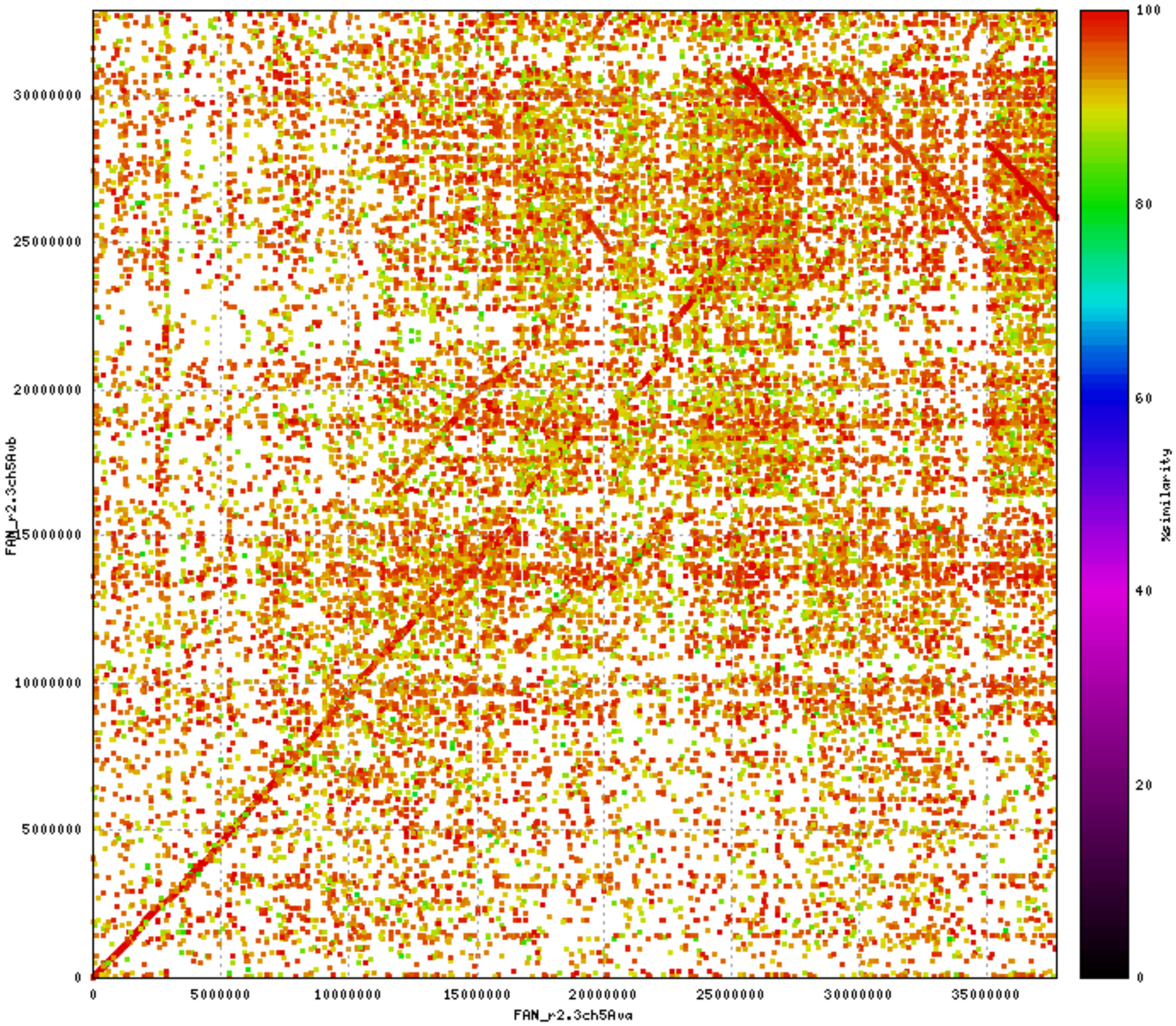

FAN_r2.3ch5Avb
FAN_r2.3ch5Ava
%similarity
0
5000000
10000000
15000000
20000000
25000000
30000000
35000000
0
20
40
60
80
100

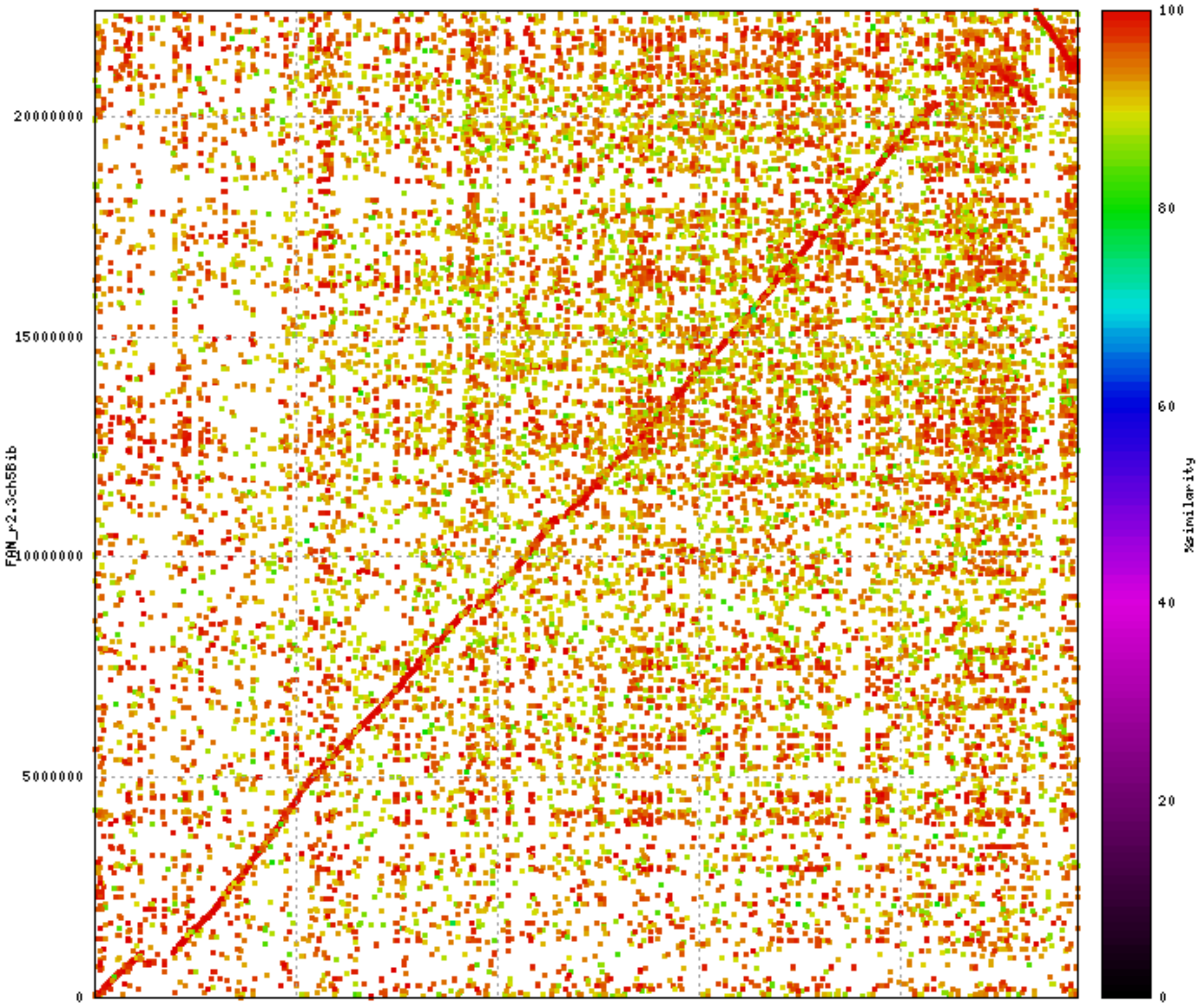

0
5000000
10000000
15000000
20000000
FAN_r2.3ch5Bia
FAN_r2.3ch5Bib
100
80
60
40
20
%similarity

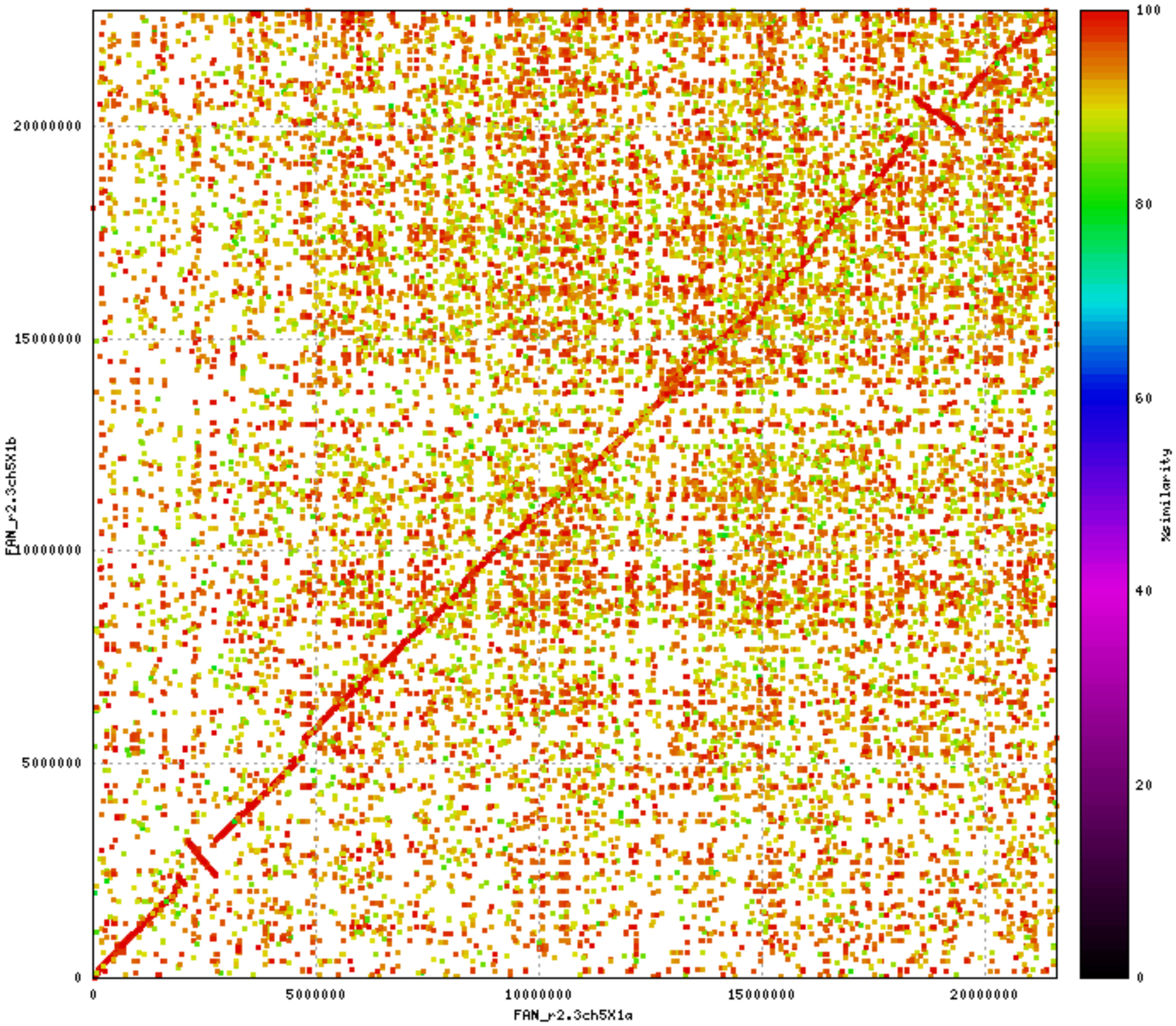

FAN_r2.3ch5X1b
FAN_r2.3ch5X1a
%similarity
0
5000000
10000000
15000000
20000000
0
20
40
60
80
100

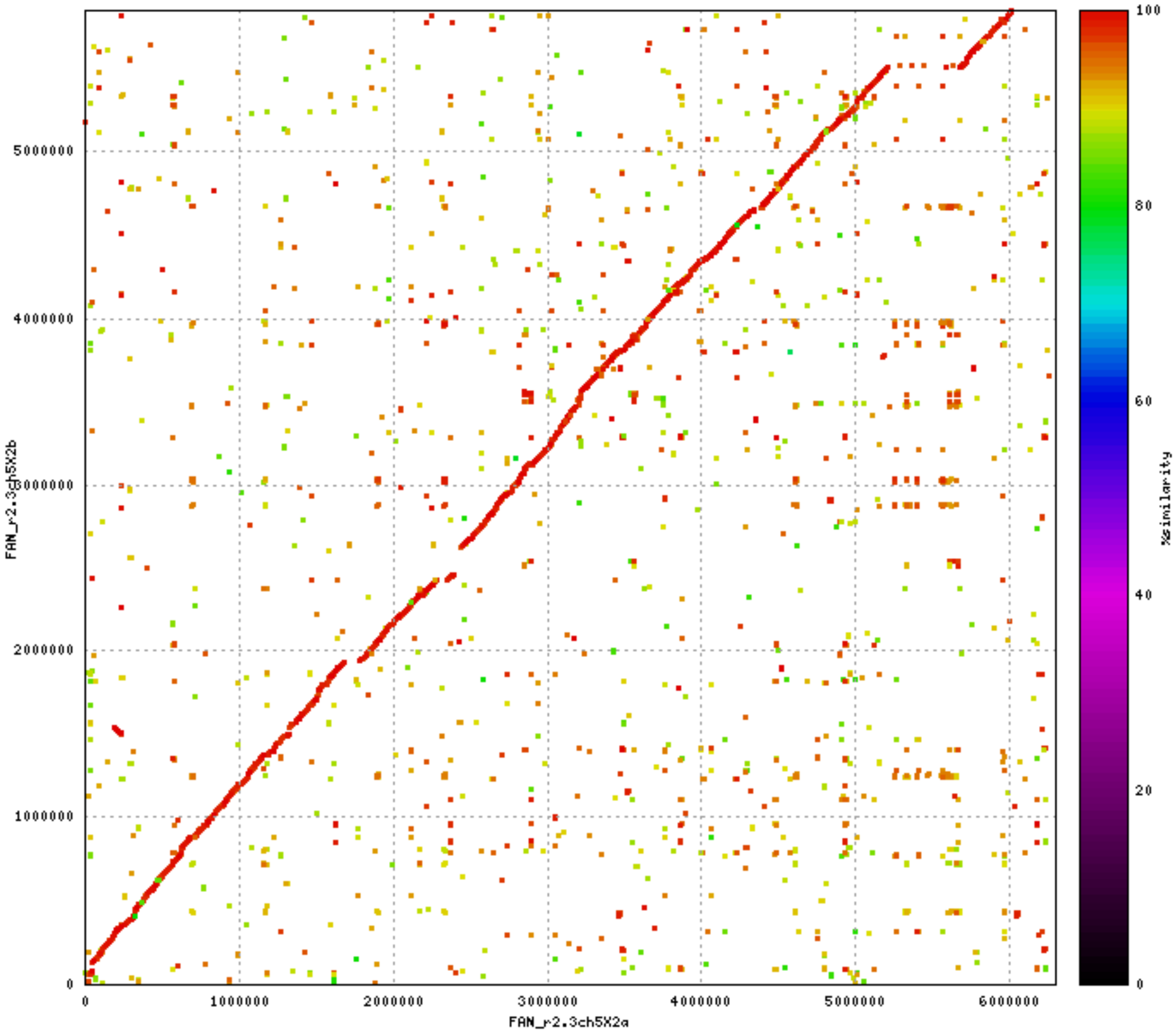

FAN_r2.3ch5X2b
FAN_r2.3ch5X2a
0
1000000
2000000
3000000
4000000
5000000
6000000
%similarity
100
80
60
40
20
0

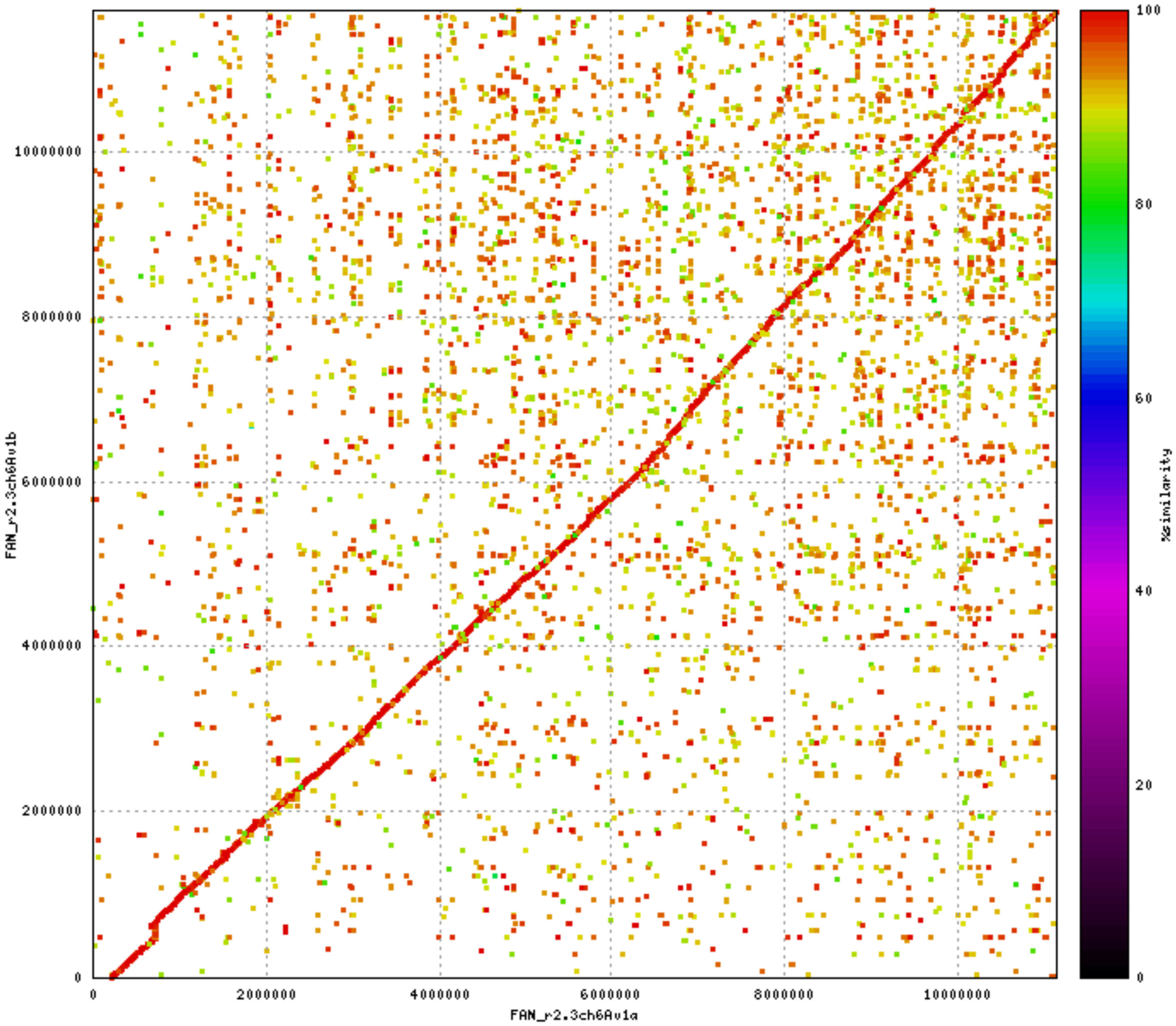

FAN_r2.3ch6Av1b
FAN_r2.3ch6Av1a
0
2000000
4000000
6000000
8000000
10000000
%similarity
100
80
60
40
20
0

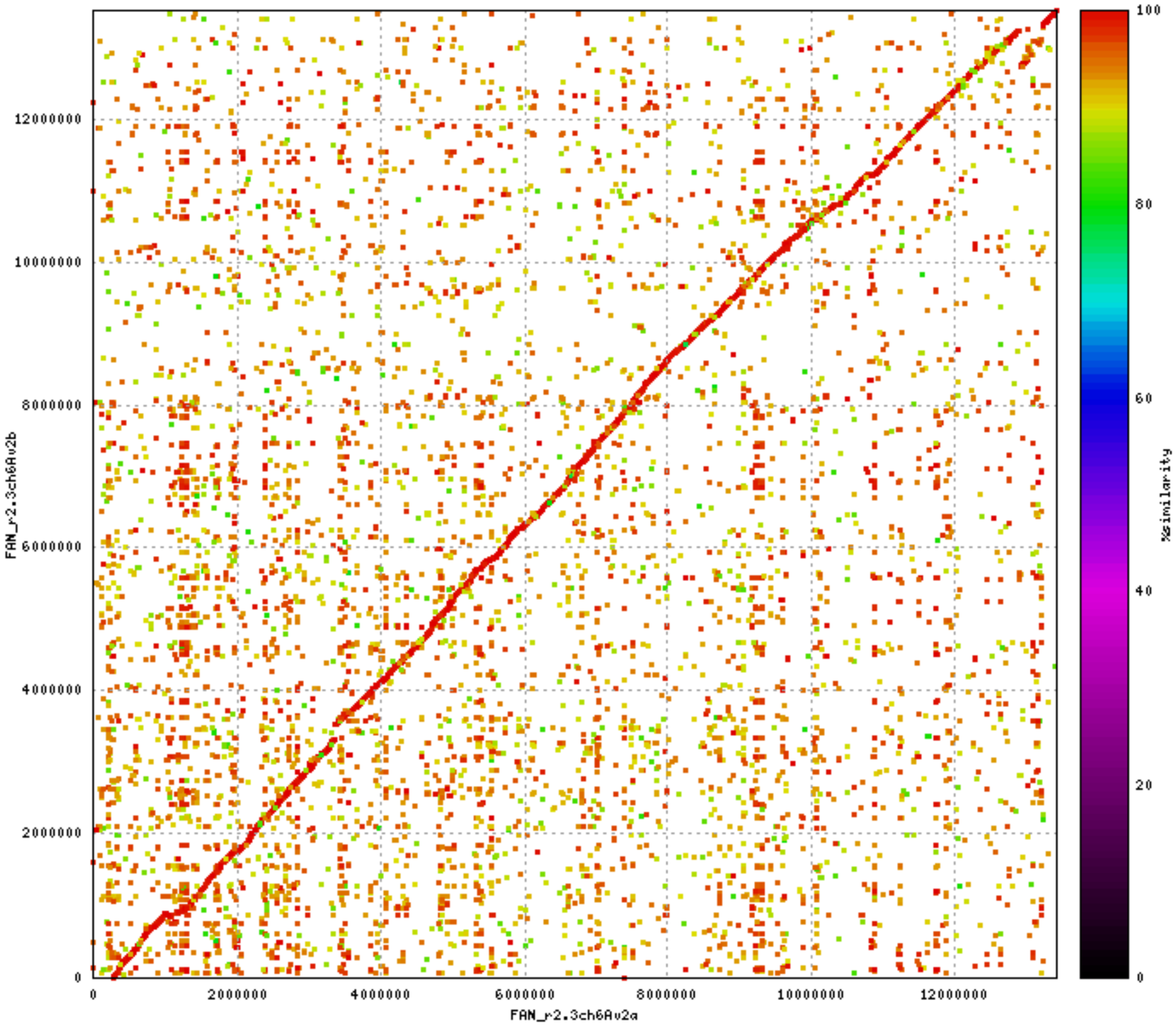

FAN_r2.3ch6Av2b
FAN_r2.3ch6Av2a
%similarity
0
2000000
4000000
6000000
8000000
10000000
12000000
0
20
40
60
80
100

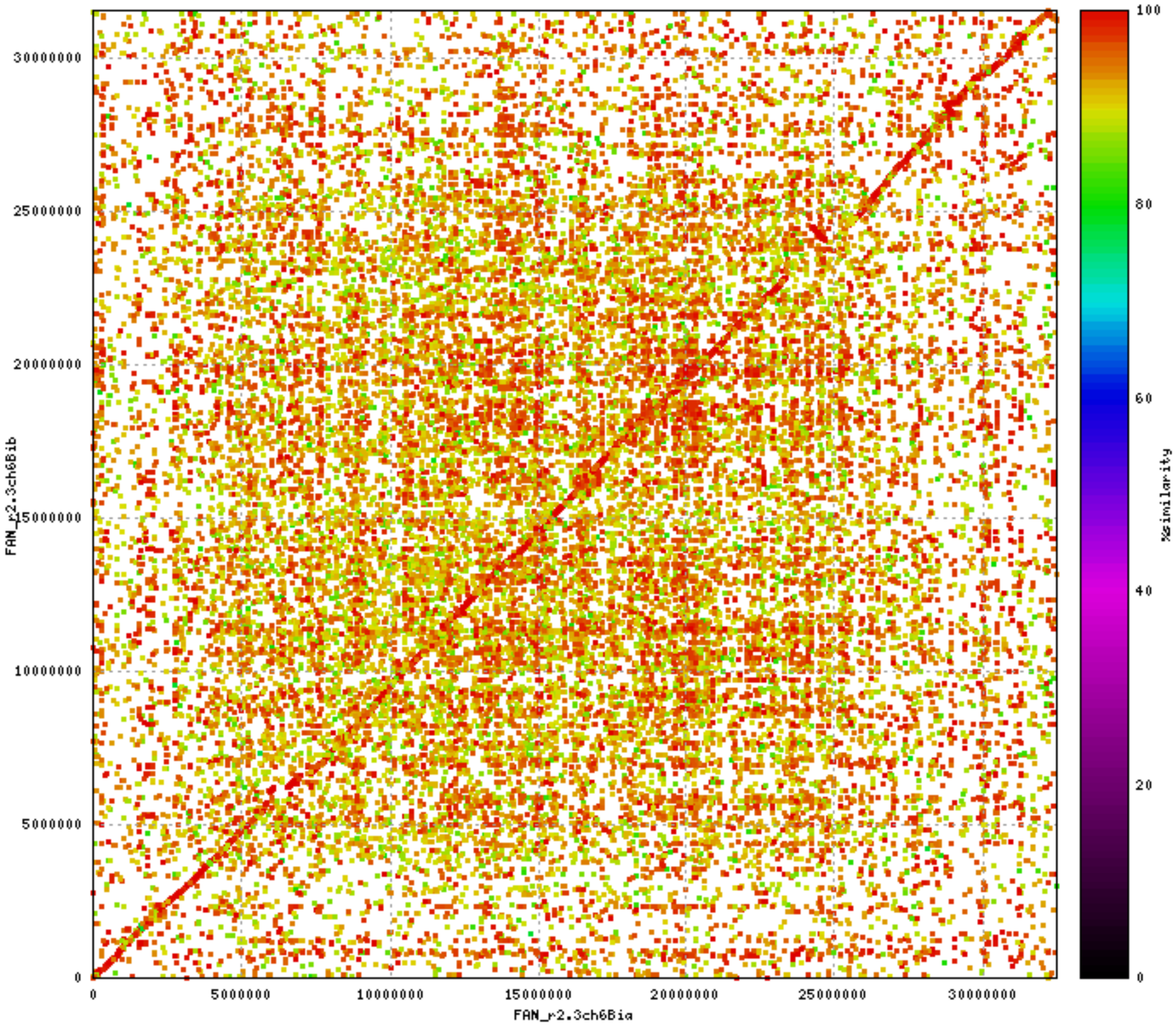

FAN_r2.3ch6Bib
FAN_r2.3ch6Bia
%similarity
0
5000000
10000000
15000000
20000000
25000000
30000000
20
40
60
80
100

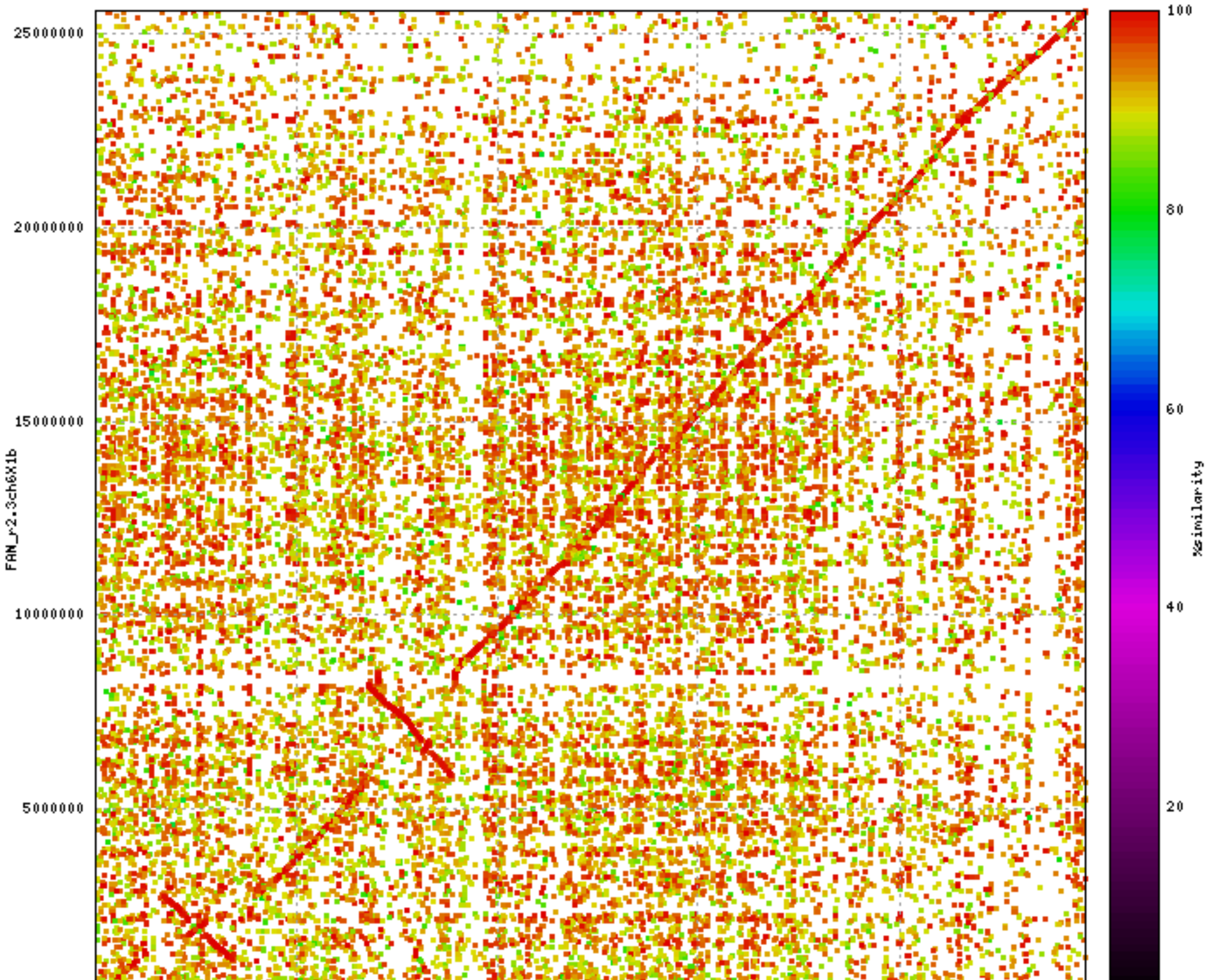

25000000
20000000
15000000
10000000
5000000
0
FAN_r2.3ch6X1b
0
5000000
10000000
15000000
20000000
FAN_r2.3ch6X1a
100
80
60
40
20
0
%similarity

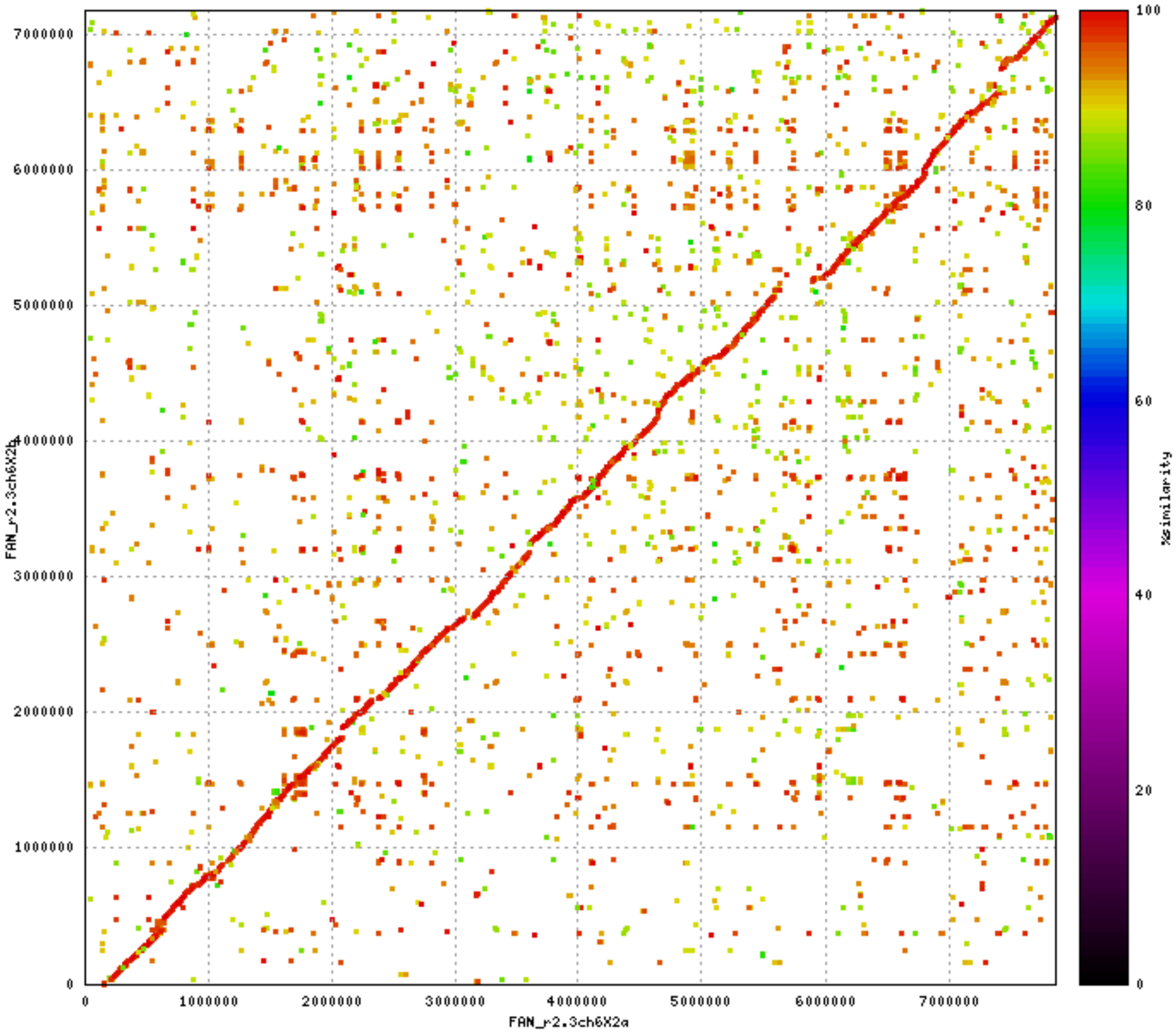

FAN_r2.3ch6X2b
FAN_r2.3ch6X2a
0
1000000
2000000
3000000
4000000
5000000
6000000
7000000
%similarity
0
20
40
60
80
100

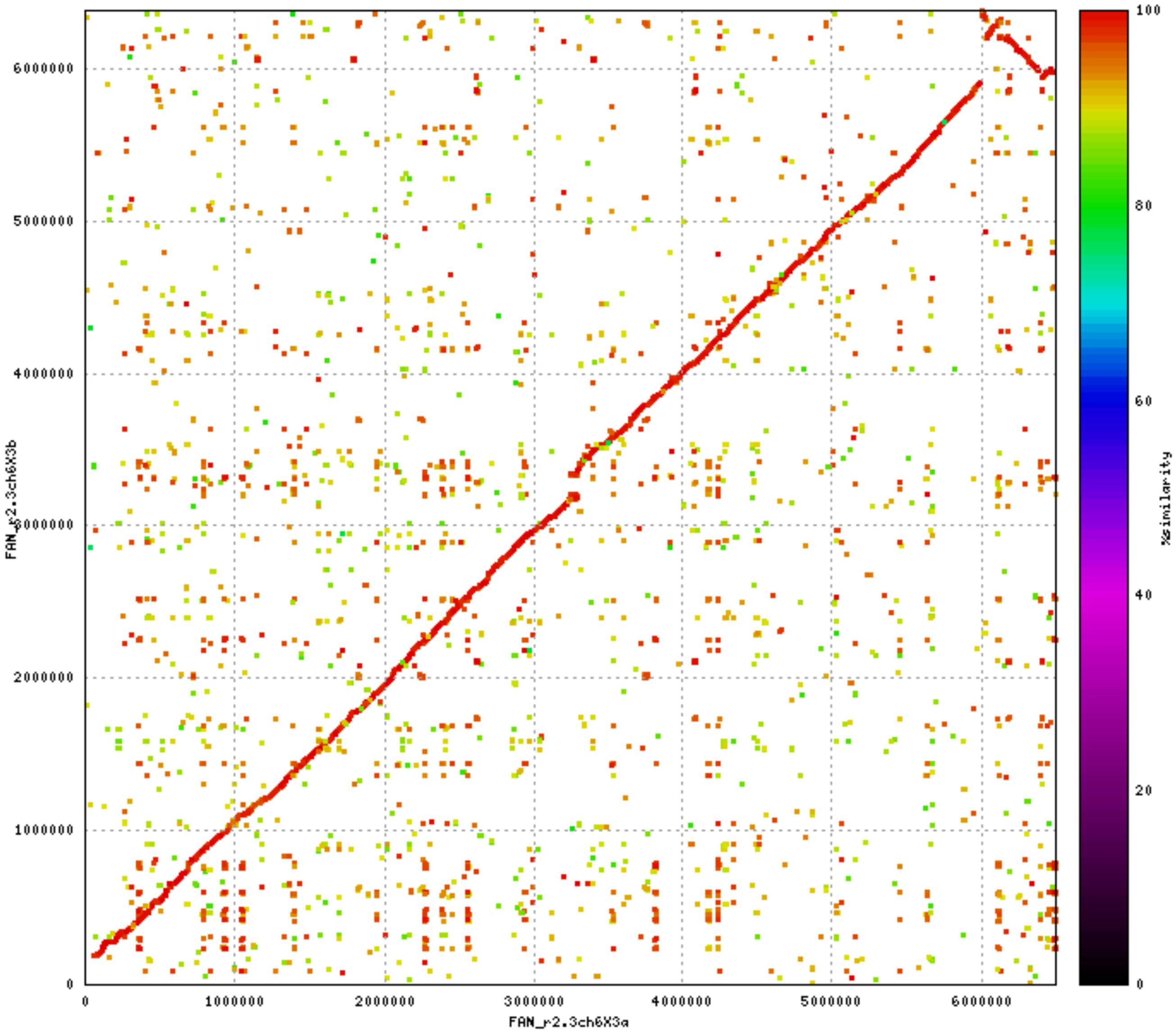

0
1000000
2000000
3000000
4000000
5000000
6000000
FAN_r2.3ch6X3a
FAN_r2.3ch6X3b
100
80
60
40
20
%similarity

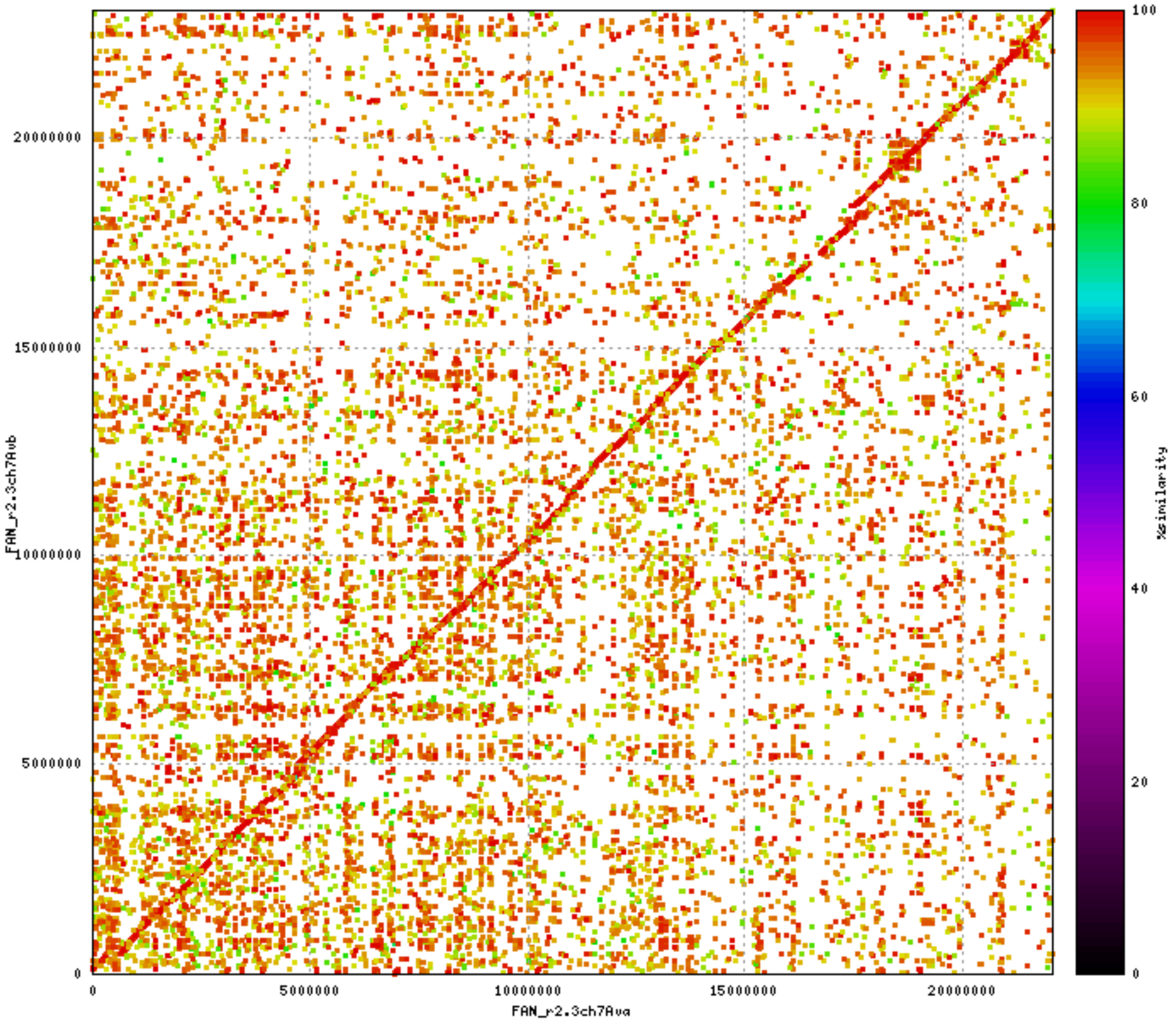

0
5000000
10000000
15000000
20000000
FAN_r2.3ch7Avb
FAN_r2.3ch7Ava
100
80
60
40
20
0
%similarity

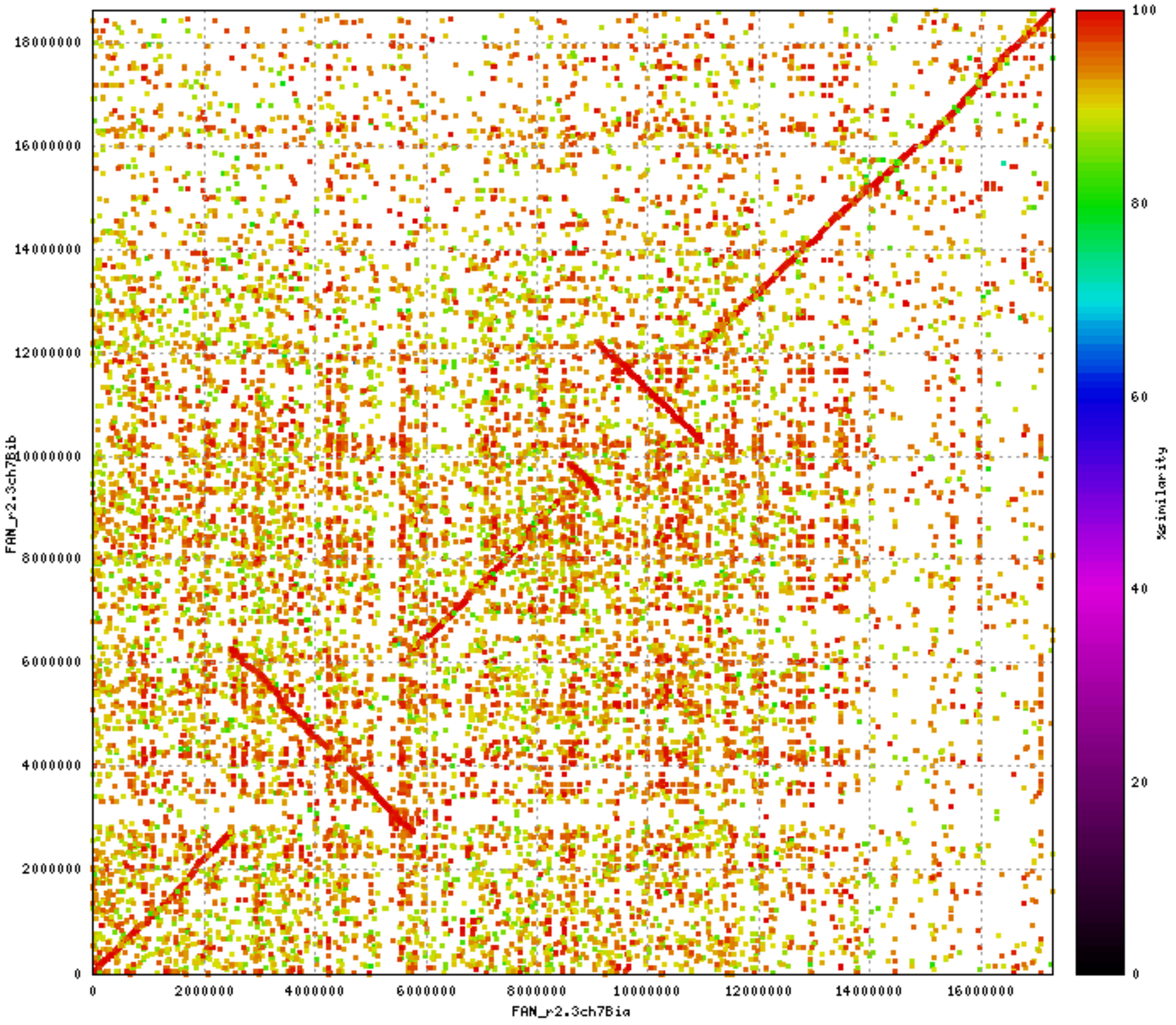

FAN_r2.3ch7Bib
FAN_r2.3ch7Bia
%similarity
0
2000000
4000000
6000000
8000000
10000000
12000000
14000000
16000000
18000000
0
20
40
60
80
100

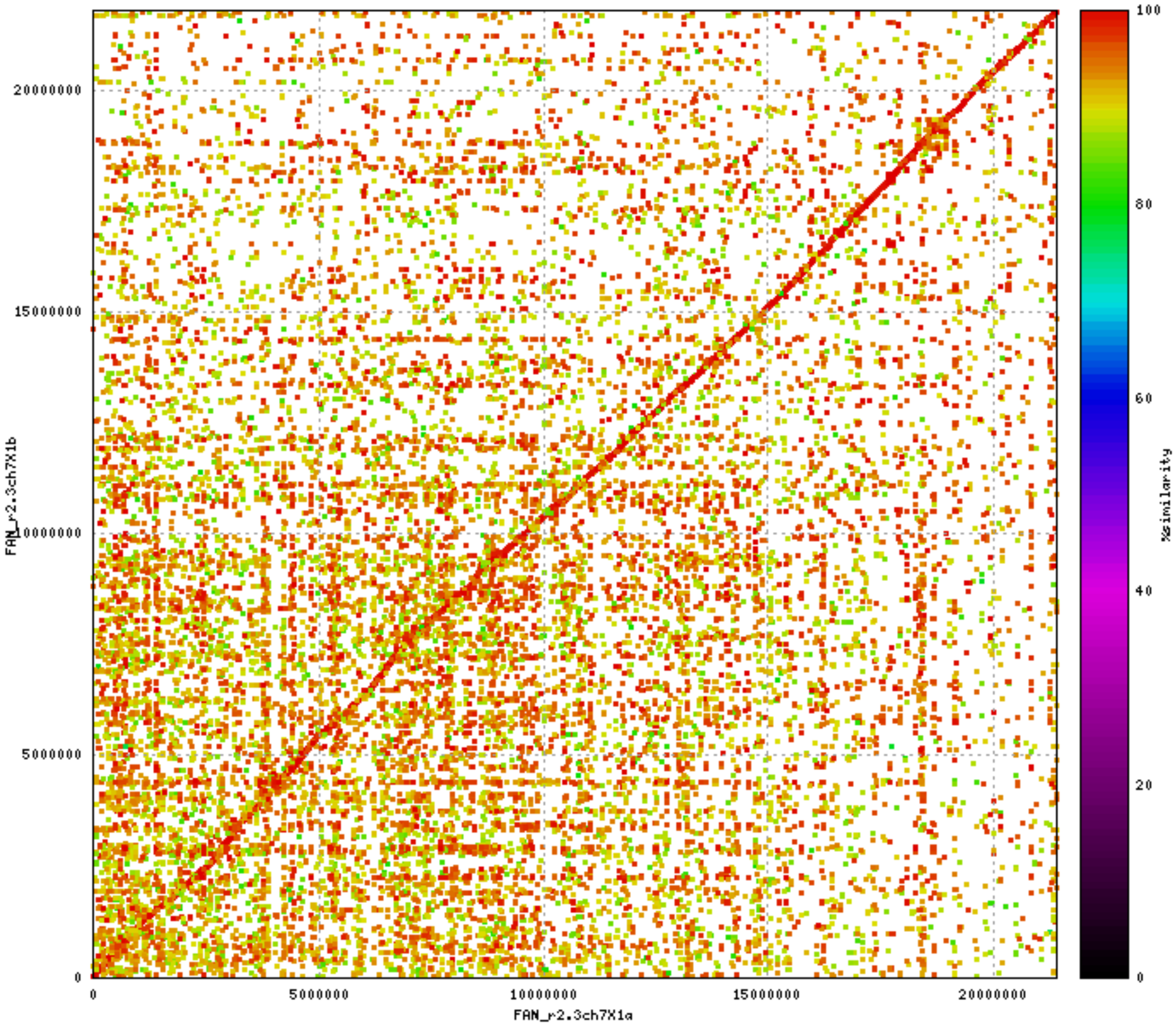
20000000
15000000
10000000
5000000
0
0
5000000
10000000
15000000
20000000
FAN_r2.3ch7X1b
FAN_r2.3ch7X1a
100
80
60
40
20
0
%similarity

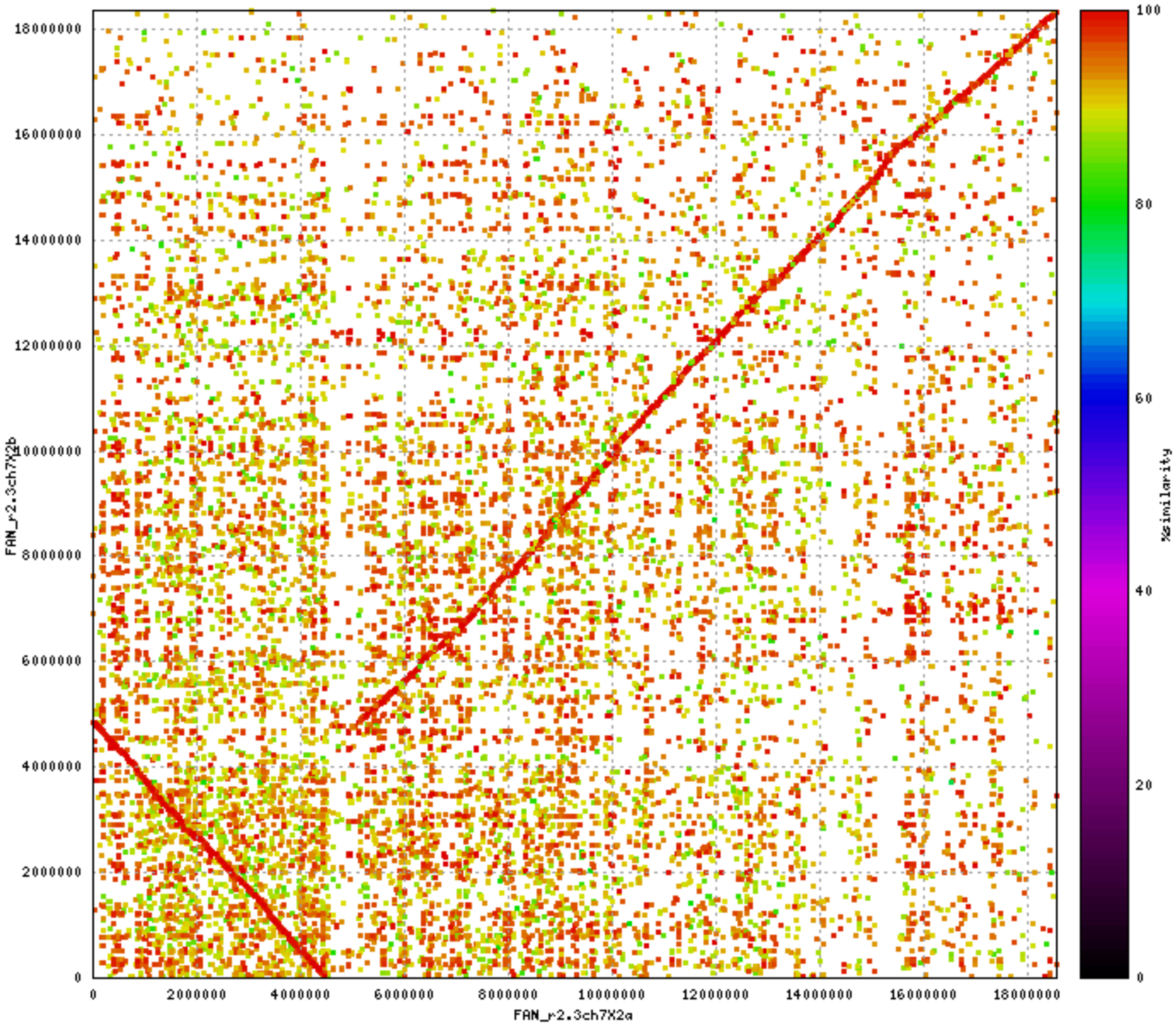

FAN_r2.3ch7X2b
FAN_r2.3ch7X2a
%similarity
0
2000000
4000000
6000000
8000000
10000000
12000000
14000000
16000000
18000000
0
20
40
60
80
100
